# Probing inter-areal computations with a cellular resolution two-photon holographic mesoscope

**DOI:** 10.1101/2023.03.02.530875

**Authors:** Lamiae Abdeladim, Uday K. Jagadisan, Hyeyoung Shin, Mora B. Ogando, Hillel Adesnik

**Affiliations:** Department of Neuroscience, University of California, Berkeley; The Helen Wills Neuroscience Institute

## Abstract

Brain computation depends on intricately connected yet highly distributed neural networks. Due to the absence of the requisite technologies, causally testing fundamental hypotheses on inter-areal processing has remained largely out-of-each. Here we developed the first two photon holographic mesoscope, a system capable of simultaneously reading and writing neural activity patterns with near single cell resolution across large regions of the brain. We demonstrate the precise photo-activation of spatial and temporal sequences of neurons in one or multiple brain areas while reading out the downstream effect in several other regions. We thus establish mesoscale two photon holographic optogenetics as a new platform for mapping functional connectivity and causal interactions across distributed cortical areas with high resolution.

## Introduction

The neural computations that support sensation, cognition and behavior depend on the precise coordination of neural activity patterns within and across widely separated brain areas. Recent advances in optogenetics and three-dimensional (3D) light patterning (Adesnik and Abdeladim 2021, Ronzitti 2017) have enabled investigators to causally probe how the activity of specific neural ensembles impact computation and drive behavior (Chettih 2019, Carillo-Reid2016, Oldenburg 2024, Marshel 2019, Dalgleish 2020, Russel 2019, Daie 2021), but only in very small, circumscribed regions of the brain on the order of <1-2mm^2^. These techniques rely on phase modulation of the optical wavefront with a spatial light modulator (SLM) allowing the user to selectively target and photo-stimulate neural ensembles of interest. Despite the power of these new read/write optogenetic approaches, the inability to apply them to distributed brain networks has prevented investigators from causally probing the logic and principles of inter-areal communication that are central to brain function. A new technology that could overcome this technical barrier would allow neuroscientists to address key outstanding questions for the first time, which could have profound importance for understanding brain function in health, disease and neural development.

The recent introduction of mesoscale two photon (2p) microscopes, which can sample neural activity with near micron precision across up to 25mm^2^ of brain tissue, have vastly increased the ability of investigators to acquire physiological data on distributed neural populations in behaving animals (Sofroniew 2016, Stirman 2016, Yu 2021, Rumyantsev 2020, Kim and Shnitzer 2022, Ota 2022, Machado 2022). While these 2p mesoscopes can monitor cellular activity, they have no ability to perturb it, preventing the user from probing causal relationships between the activity in connected brain areas and behavior. Thus, if it were possible to develop a 2p mesoscope that can not only measure but also manipulate neural activity with cellular resolution, neuroscientists could use such a system to address longstanding mysteries of long-range neural communication in the brain.

However, there are multiple significant technical challenges to achieving high resolution holographic optogenetics in a 2p mesoscope. Existing commercial 2p mesoscopes were not designed for the integration of holographic systems, requiring a significant redesign of the mesoscope build. A particularly outstanding challenge in achieving high-resolution patterned photostimulation across a mesoscopic field-of-view (FOV) is the physical limits imposed by spatial light modulators (SLMs) designs which ultimately constrain the accessible photostimulation FOV. We first designed and optimized a flexible 3D holographic system fully integrated onto a 2p random-access (2p-RAM) mesoscope that enables near single-cell resolution holographic photo-stimulation of neural ensembles in the brain across a millimetric FOV. We tested and validated the optical capabilities of this new read/write platform to measure and recreate highly specific patterns of activity in the brain. In particular, we showed that we can decode the identity of the specific photo-stimulus purely from the modulation of activity of postsynaptic neurons in downstream areas, and that we can holographically recreate and transmit visual-like information across areas in mouse visual cortex. Next, we expanded the photostimulation FOV by an order of magnitude by integrating a random-access scan module to the 2P holographic path. The compounded constraints of the 4f architecture of 3D-SHOT, the need to maintain conjugation and coplanarity, the steric constraints of large 3-inch optics, the limited off-the-shelf optical components, and the overall space constraints on the added breadboard constituted significant challenges that had to be jointly overcome to both maintain the specifications of the system at the central FOV and expand the holographic FOV. We further showed how one can use this system to probe functional interactions between distant brain regions. Finally, we demonstrated for the first time near simultaneous photostimulation of specific neural ensembles across distinct remote cortical areas. Together, these data illustrate how our 2P holographic mesoscope enables the user to probe causality both at the local and the interareal level with near single-cell resolution.

## Results

### A 2P-RAM mesoscope with temporally focused 2P holographic illumination

We designed a mesoscale all-optical read/write platform around a two-photon random access (2P-RAM, Sofroniew 2016) system (**Fig 1a-c, Supp. Fig 1**) that is commercially available (Thorlabs, Inc.). The key features of this imaging system that make it a powerful tool for studying inter-areal communication are its nominal 5x5mm FOV, a fast remote focusing system for FOV curvature correction and multiplane imaging, and four automated axes of motion for positional flexibility (**Fig1c**). We sought to develop a mesoscale holographic optogenetics system that achieves near cellular resolution photo-stimulation despite the lower numerical aperture of the optical path and that could likewise address the imaging plane curvature. Furthermore, we aimed for an optical design that does not compromise the 4-axis movement capabilities of the 2P-RAM system, since they are essential for executing a diverse array of neurobiological experiments. We therefore designed a compact and re-dimensioned 3D holographic module that employs temporal focusing to confine excitation axially (3D-SHOT, three-dimensional sparse holographic optogenetics with temporal focusing Mardinly 2018, Pégard 2017). To preserve all translational axes of the entire system, we built the holographic module on an extension breadboard attached to the movable main frame of the microscope (**Fig1b, Supp. Fig 1, Supplementary note 2**). In 3D-SHOT, an initial blazed grating generates a temporally focused disc of light that is magnified and replicated in multiple locations within a 3D volume by a spatial light modulator (SLM). The 3D light pattern is then relayed onto the objective image space through a set of 4f-systems (**Fig 1c**). Given the large distance on the mesoscope between the photostimulation laser output and the 3D-SHOT module, we added a non-magnifying telescope after the laser to control the divergence of the beam impinging on the diffraction grating. Preserving the alignment invariance of the holographic path with the motorized movements of the mesoscope is a key challenge. To address this (**Fig1c**), we first combined the imaging and photostimulation beam before entering the multi-stage periscope so that the photostimulation beam maintains invariance with X and Y frame translations. Second, we replaced the last periscope mirror with a long-pass dichroic that directs the imaging beam (920nm) onto the vertical breadboard and the photostimulation beam (1030nm) onto the extension breadboard so that the latter maintains invariance with rotation. Third, we recombined both beams with a short-pass dichroic placed right before the mesoscope tube lens on the vertical breadboard. Consequently, the holographic module is invariant within all dimensions controlled by the motorized microscope frame, X, Y translations and rotation. We recovered axial translation by adding a motorized stage for animal positioning along the vertical axis, additionally mounted on a goniometer to ensure that imaging and photo-stimulation are fully orthogonal to the cranial window surface.

**Figure 1.**
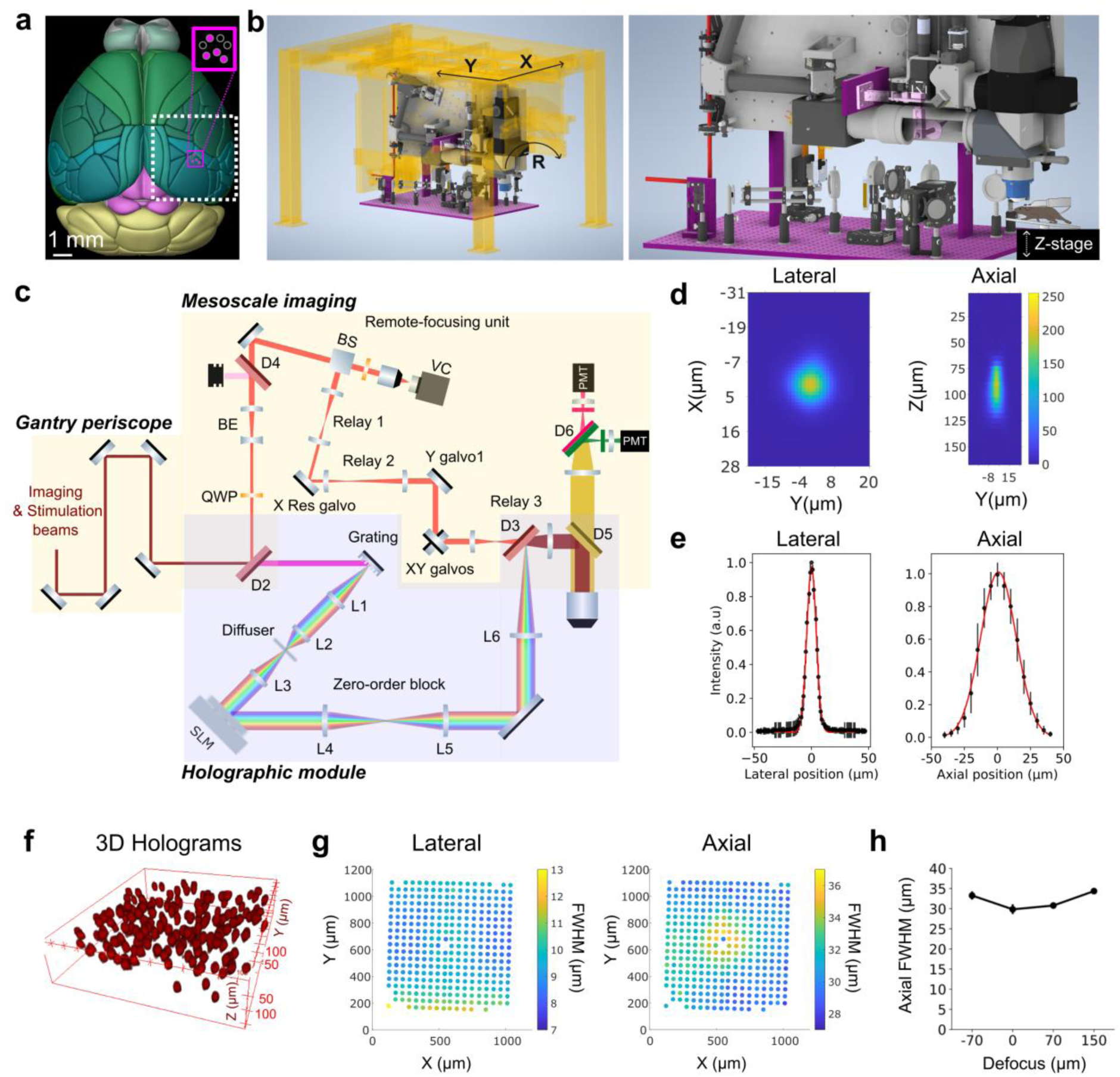
A 2P-RAM mesoscope with temporally focused 2P holographic photostimulation. **a)** Schematic dorsal view of the mouse brain with putative functional areas demarcated (Allen Brain Explorer). The holographic mesoscope allows targeted photostimulation in a 1x1 mm FOV (magenta square) combined with simultaneous mesoscale imaging across a nominal 5x5 mm imaging FOV (dotted white). **b)** Left: CAD model of the holographic mesoscope platform. Right: blow up of the holographic module attached to the main frame of the mesoscope. The 3D-SHOT holographic module consists primarily of the highlighted magenta parts and the optics on the highlighted extension breadboard. **c)** Simplified schematic of the optical path. QWP: quarter-wave plate, BE: beam expander, Ln: achromatic doublet, Dn: dichroic mirror. **d)** Example lateral and axial 2P fluorescence intensity profiles of a single, temporally focused holographic illumination spot. **e)** Optical lateral and axial point-spread-functions (PSFs). Error bars: standard error of the mean (S.E.M). Red curve: Gaussian fit, lateral full-width-half-maximum (FWHM) 9 µm, axial FWHM: 32 µm. **f)** Example point-cloud 3D holograms. **g)** 2D grid of single-spot holograms showing the extent of the holographic FOV and the distribution of lateral and axial resolution across the FOV. **h)** Axial PSF as a function of digital Z-defocus.

To quantify the optical resolution of the mesoscale holographic system, we first measured the spatial profile or ‘point spread function’ (PSF) of single two photon excitation spots (**Fig. 1d-e**) which measured 9µm ± 0.7 μm in the lateral dimension and 32µm ± 1.6 μm in the axial dimension (mean ± s.d, n=538 holograms arranged in a 2D grid, see also **Fig. 1g, Supp** **Fig 2**). Although these values are slightly larger than those previously reported for 3D-SHOT in a conventional 2p microscope, they are consistent with the lower NA of the mesoscope objective (**Supplementary note 1**). We then determined the extent of our holographic field-of-view (FOV) which is equivalent to the total accessible FOV range of the SLM at the system’s magnification. We set the FOV limits to the farthest accessible holograms without aliasing which corresponded to a total lateral FOV of ∼950 μm × 990 μm (**Fig. 1g**), which closely approximates the theoretical SLM FOV at this magnification (see **Methods**). We compensate for the inherent decrease in diffraction efficiency laterally across the FOV by adjusting the power delivered to each spot according to its spatial location (**Supp** **Fig. 2**). Notably, we observed that while holograms farther from the FOV center had expectedly lower intensity profiles (due to lower diffraction efficiencies), off-axis holograms remained only slightly affected by off-axis aberrations (**Supp. Fig 3**). Next, we generated multiple holograms spatially arranged in a 3D point-cloud (**Fig. 1f**). We then measured the axial optical PSFs in 3D and found that they were mostly invariant within a digital defocusing range of 220 µm (**Fig. 1h**). A potential challenge for holographic targeting on the 2P-RAM mesoscope is the large microscope field curvature, 160 μm over the 5mm FOV, mostly introduced by the mesoscope objective. We hence quantified the flatness of our holographic FOV, by measuring the axial locations of holograms generated with equi-Z SLM coordinates and found the maximal peak-to-valley to be ∼ 40 𝜇𝑚 which is well within the addressable defocus range of the SLM and can be corrected for without compromising 3D targeting (**Supp Fig2**). Another potential challenge for multiphoton holographic optogenetics on the 2P-RAM mesoscope is the introduced dispersion from the 4f lenses and the large mesoscope optics. We estimated the total group delay dispersion (GDD) introduced by the holographic path to be ∼15000 𝑓𝑠^2^ (**Supplementary table 2**). For relatively broad pulses commonly used in multiphoton optogenetics (∼400𝑓𝑠), the broadening of the pulse width at the objective focus is minimal, thus discarding the need for a pulse compression unit on the holographic path.

**Figure 2.**
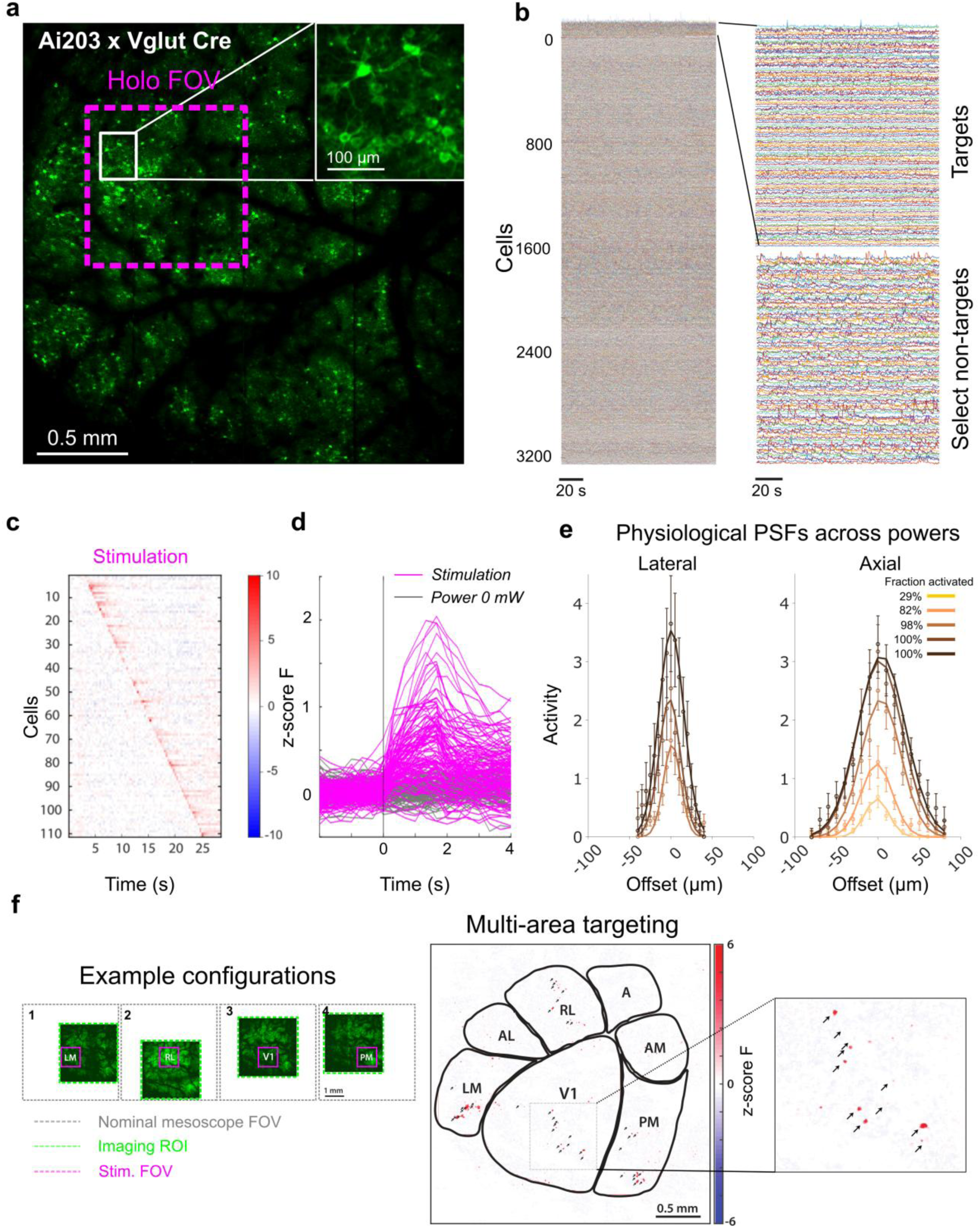
Cellular resolution two-photon holographic optogenetics at the mesoscale. **a)** A 2p mesoscale image of widespread co-expression of GCaMP7s and ChroME over a large field-of-view (FOV) in an Vglut1-Cre;Ai203 transgenic mouse. Magenta square indicates the accessible holographic FOV within an example mesoscale imaging FOV of 2.4 mm × 2.4 mm recorded at 4.5Hz framerate. **b)** Example single trial calcium traces from a set of photo-stimulation targets within the holographic FOV and thousands of traces from non-targets recorded across the mesoscale FOV. **c)** Calcium responses of targeted cells during single-cell sequential targeting on stimulation trials (right). Photostimulation with 5x5ms pulses at 30Hz at 75mW. Traces averaged over 5 trials. **d)** Stimulus-aligned target cell responses across the holographic FOV (n=111 cells). Single-trial traces. **e)** *In vivo* optically measured lateral (left, n=20 cells) and axial (right, n=32 cells) physiological point-spread functions as a function of stimulation power. Absolute powers used are 0.5mW, 1mW, 2mW, 4mW and 8mW in a CaMK2-tTA;tetO-GCaMP6s mouse, corresponding to a fraction of activated cells among targeted cells of 29%, 82%,98%,100% and 100% respectively (see supp. Fig. 8). The filled circles and error bars are the actual mean +/- s.e.m. across cells, and the solid traces are weighted gaussian fits. For lateral PPSFs, fitted FWHMs are 28.5µm, 31.9µm and 38.4µm for 2,4 and 8mW respectively. For axial PPSFs, fitted FWHMs are 35.4µm, 46µm, 59.8µm, 66.4µm and 70.4µm 0.5, 1, 2,4 and 8mW respectively. **f)** Demonstration of sequential area mesoscale holographic stimulation. Left: 4 example configurations to holographically target 4 different visual areas within the same session and record from surrounding visual areas. Imaging ROIs were selected for a 3 Hz imaging rate at 1µm/pixel. Photostimulation with one single 250ms pulse 80mW. Center: Activation map of holographically targeted neurons in 4 visual areas (V1, PM, RL and LM). Black arrows point to target locations. Right: Zoomed in inset of the activation map in V1.

Finally, we assessed holographic targeting stationarity with X, Y translations and rotation, which is essential for accurate optical generation of neural activity patterns despite movement of the mesoscope relative to the brain. We found that the error between original hologram location and at displaced stage positions remained less than 3 µm over a ± 3mm translation range and a ± 2.5° rotation angle range demonstrating alignment invariance with mesoscope frame movements (**Supp. Fig 4**).

### *In vivo* 2P photostimulation on the 2P-RAM mesoscope

Next, we tested our system for *in vivo* all-optical control of neural ensembles in awake mice. We tested the system on mice expressing soma-targeted opsin either transgenically or via viral injection. In Vglut1-Cre;Ai203, mice (Bounds, 2021) which express the potent opsin ChroME and the sensitive calcium indicator (GCaMP7s) in cortical excitatory neurons (**Fig. 2a**) we were able to photoactivate selected individual cells within the holographic FOV while recording from thousands of neurons within the mesoscale imaging FOV (**Fig. 2b-e**). Sequential photostimulation at 75mW/spot (10 × 10 ms pulses at 50Hz) elicited significant time-locked responses in 70% of targeted cells (**Fig. 2c**,**d****, Supp** **Figure 5**). This demonstrates effective holographic photostimulation despite the slightly larger optical axial resolution. Next, using viral transduction with a soma-targeted ChRmine-expressing AAV, we further demonstrated that we could reliably and simultaneously photo-activate large groups of neurons (**Supp. Fig 6,7**). To carefully quantify the effective spatial resolution of holographic photo-stimulation, we recorded physiological point-spread-functions (PPSFs) all optically in awake animals in a wide range of conditions (**Fig.2e, Supp 8,9,10**). Physiological point-spread functions can vary wildly depending on multiple parameters both optical and linked to the preparation (**see Supp. Note 2**). We measured lateral PPSFs between 23 µm and 38 µm across conditions (n=23 cells, pooled across ChroME and ChRmine cells), and axial PPSFs between 35µm and 77µm across conditions (n=35 cells, pooled across ChroME and ChRmine cells). This indicates that despite the lower NA inherent to the mesoscope objective, the achieved effective axial resolution *in vivo* is within the range of reported PPSFs in 2p optogenetics studies (80µm in Daie et al. 2021, 63µm in Fisek et al. 2023, 47 µm in Paker et al. 2015) and that in certain conditions near-single cell resolution can be achieved with our system (**Supp. Fig 8,9**).

**Figure 3.**
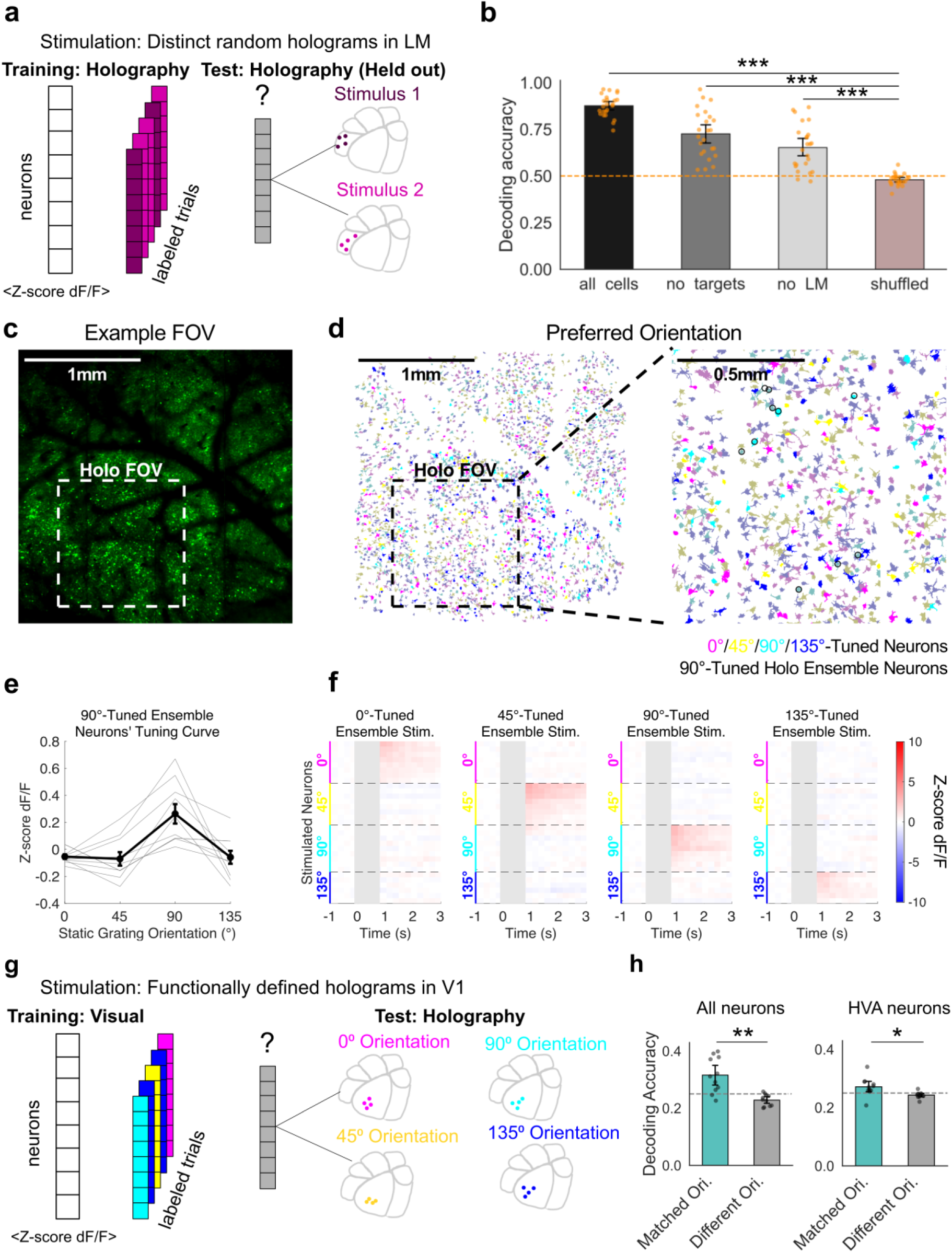
Specific transmission of information at the mesoscale upon targeted local 2P photostimulation. **a)** Schematic of a decoder trained to discriminate different holographically induced patterns of activity from downstream responses. Multi-target (3-20 cells) stimulation in LM (n=4 sessions, 2 mice expressing GCaMP7s and ChroME), with 10×10ms pulses at 16Hz and 50mW. Recording rate: 4.5Hz. **b)** Decoding accuracy of a classifier trained to discriminate the holographically induced patterns in LM from different groups of non-targeted cells in the FOV. All cells: decoding from all neurons in the mesoscale FOV. No targets: decoding from all neurons except the targeted cells. No LM: decoding from all neurons outside area LM. Shuffled: decoding from all neurons outside area LM but with training set labels shuffled. The cross-validated decoding accuracy across N = 4 sessions for all cells, no targets, no LM and shuffled categories was, respectively, 0.88 ± 0.01, 0.73±0.03, 0.65±0.02 and 0.48±0.01 (mean ± SEM), the chance level being at 0.5.Wilcoxon signed-rank test across N = 4 sessions: All cells/ no targets: 4.89×10^-5^, all cells/ no LM: 1.67× 10^-6^, no targets/ no LM: 0.011, all cells/ shuffled: 1.67×10^-7^, no targets/ shuffled: 1.32×10^-9^, no LM/shuffled: 5.96×10^-7^. All groups (all cells, no targets, no LM) show highly significant decoding performance (p-value <0.001) compared to the shuffled group. **c)** Example imaging FOV used in this experiment (2.4 × 2.4 mm^2^ at 4.5Hz recording rate, Vglut1-Cre;Ai203 mouse with neurons co-expressing GCaMP7s and ChroME). The holographic FOV was positioned over V1. **d)** Same FOV as **a**, with the cells color coded by their preferred orientation (0/45/90/135° in magenta/yellow/cyan/blue, respectively). Significantly tuned cells are labelled with solid colors while other cells are grayed out (p<0.05 Wilcoxon rank-sum test for preferred vs orthogonal orientations). Right: enlarged image of the holographic FOV with thin circles indicating the holographic targets that were activated by the 90°-tuned ensemble stimulation. **e)** Orientation tuning curves of a group of neurons that were then co-activated as the 90°-tuned ensemble stimulation in f). Photostimulation with 10×10ms pulses at 16Hz and 50mW. **f**) Holographically evoked activity in a population of neurons that were grouped into four distinct orientation-specific ensembles. Imaging frames recorded during the holographic stimulation are grayed out due to the holographic stimulation artifact. **g)** Schematic of a decoder trained on visual trials and tested on holographic trials. **h)** Decoding performance in assigning holographic trials (functional specific holographic perturbations) to matching visual trials (oriented gratings). Decoding accuracy was measured as the proportion of holography trials that were classified as the grating orientation corresponding to the photo-stimulated ensemble’s orientation preference. Given that 4 orientations are probed, chance level is at 0.25. Left: Decoder trained and tested on all neurons in the FOV (0.316 ± 0.0185 mean ± SEM matched orientation, 0.228 ± 0.0062 mean ± SEM different orientation, p = 0.0098 Wilcoxon signed-rank test across N=11 sessions). Right: Decoder trained and tested only on neurons in downstream extrastriate areas, excluding V1 (0.271 ± 0.0096 mean ± SEM matched orientation, 0.243 ± 0.0032 mean ± SEM different orientation, p = 0.04 Wilcoxon signed-rank test across N=11 sessions).

**Figure 4.**
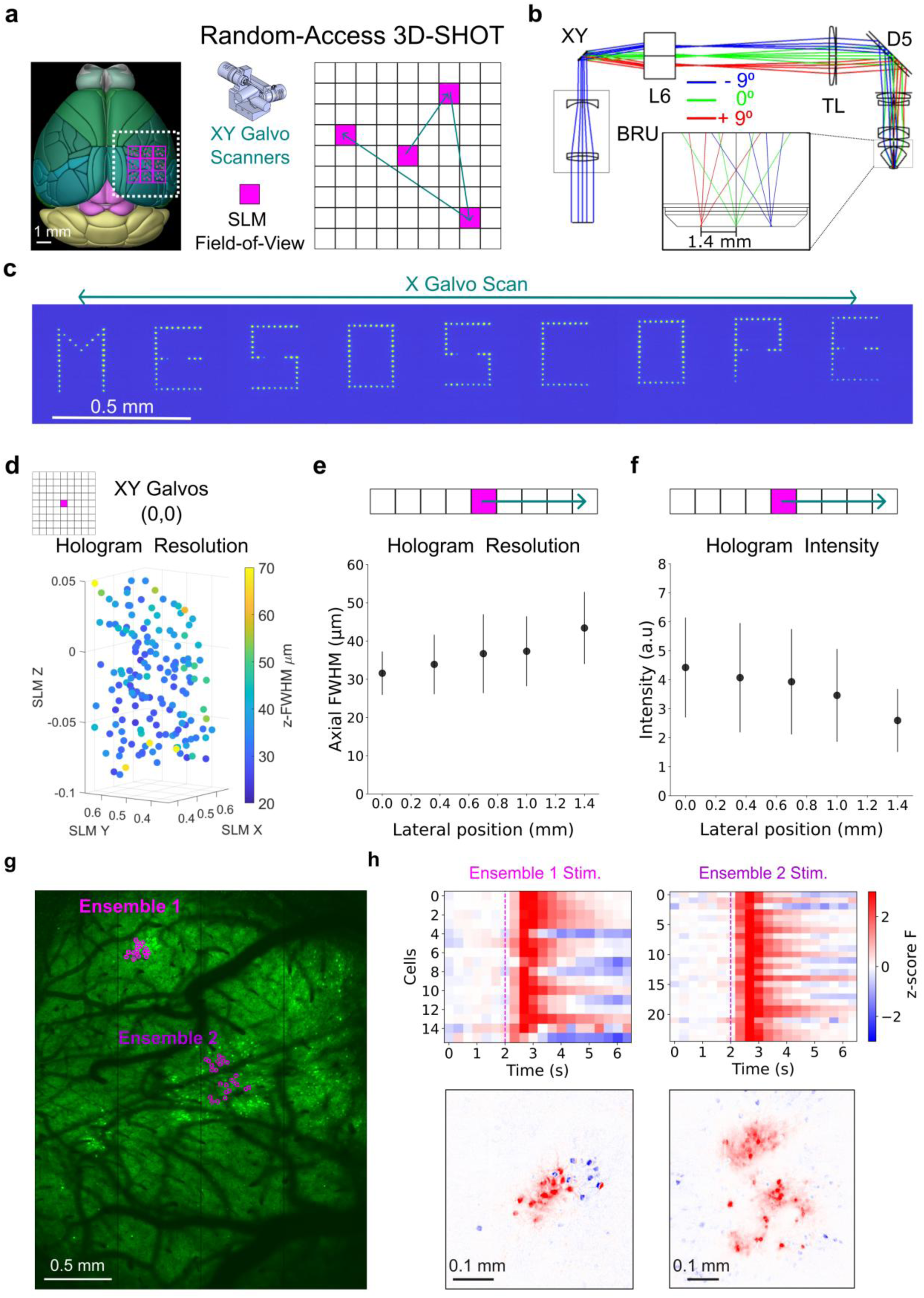
Mesoscale two-photon optogenetics with random-access 3D-SHOT. **a)** Principle of random-access 3D-SHOT. XY Galvanometer scanners are used to deflect the center of the SLM FOV thus effectively expanding the accessible holographic stimulation area. Mouse brain schematic adapted from Allen Brain Explorer. **b)** Zemax optical model of the added scanning module implemented for FOV expansion. BRU: Beam Reducer Unit. Compact Galilean telescope with a 2x beam reduction factor. XY: Large aperture 30mm galvanometer scanners. L6: f-theta scan lens. ± 9° optical deflection enables a ± 1.4mm shift of the center of the SLM FOV in each direction. **c)** Holographic multi-dot fluorescence patterns generated with random-access 3D-SHOT. Each letter was generated at a distinct X galvanometer position, ranging from –2 V (–9° optical deflection) to +2 V (+9°), in 0.5 V increments. Holographic letter patterns were recorded independently using a substage camera, which was translated by 350 µm between each recording. All patterns were stitched together to reconstruct the full 2.8 mm scan range. Occlusions at the center of targeted SLM FOVs correspond to the position of the zero-order block that is used to block non-diffracted light. **d)** Axial optical point-spread-function measured on a 3D distribution of spots (n=154) in the central holographic FOV (X galvo = 0°, Y galvo = 0°) within a 400µm×400µm×150µm volume. Mean axial FWHM = 34µm. **e)** Mean axial resolution measured across 3D distribution of spots at different galvanometer scan positions. See **supp. Fig. 14** for more extensive characterizations. Error bars indicate standard deviation values. **f)** Mean single-hologram intensity measured across 3D distribution of spots at different galvanometer scan positions. See **supp. Fig. 14** for more extensive characterizations. Error bars indicate standard deviation values. **g)** Mesoscopic 3.1mm×2.4mm FOV of a CaMK2-tTA;tetO-GCaMP6s mouse visual cortex injected with a pAAV-hSyn-GCaMP6m-p2A-ChRmine-Kv2.1-WPRE virus at discrete locations. Photostimulation targets are segmented from the mesoscale FOV and assigned specific combinations of targeting phase masks and XY galvanometer positions (see Methods). **h**) Top: Post-stimulus time histograms (PSTHs) of activated neural ensembles represented in g) and arbitrarily located in the mesoscopic FOV. Bottom: Spatial fluorescence maps showing the mean pixel-by-pixel baseline subtracted fluorescence intensity across 2.3 s (7 frames) post-stimulation. Scale range: -2×10^-3^ (blue) to +2×10^-3^ (red).

**Figure 5.**
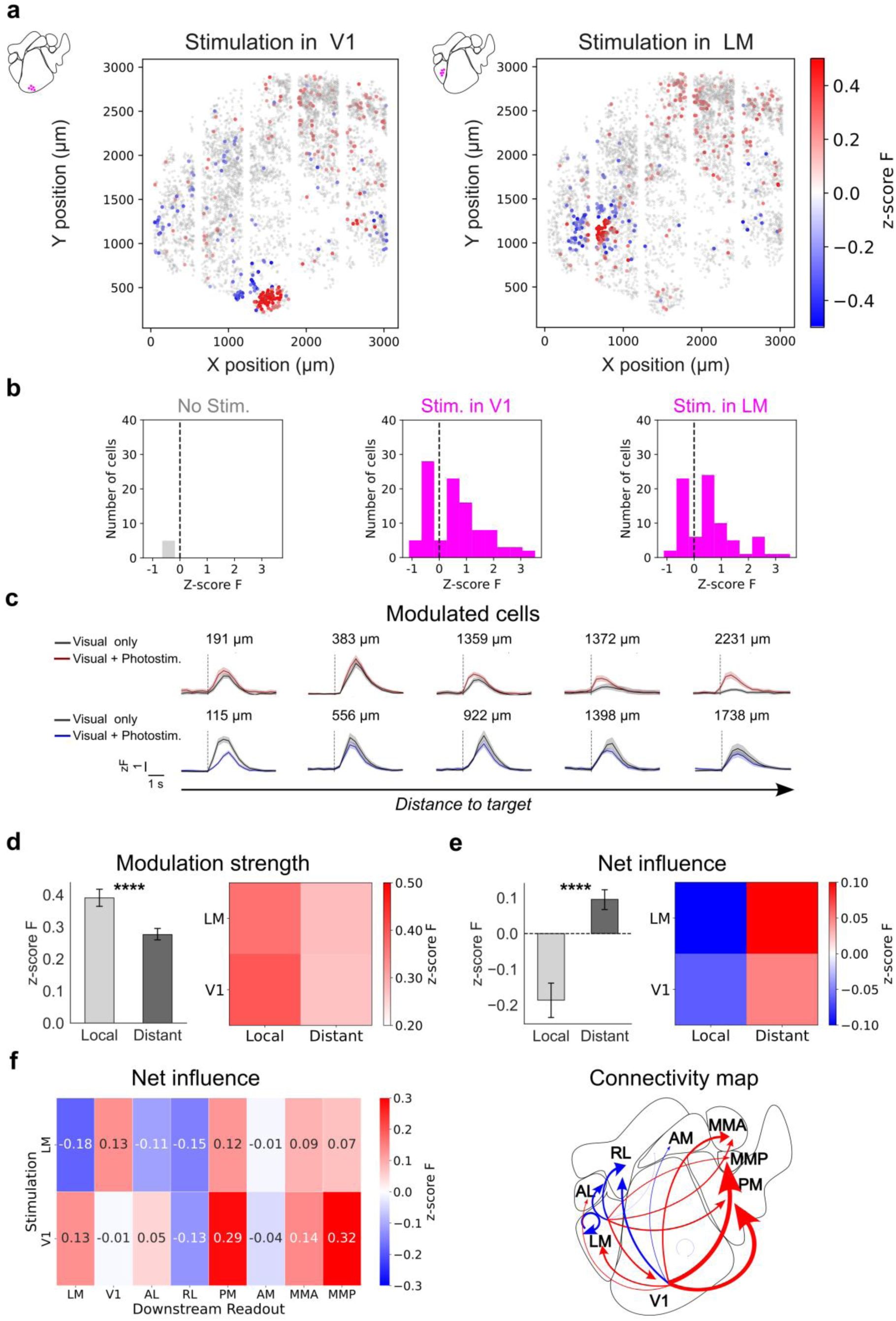
Mesoscale causal probing of connectivity across cortical areas. **a)** Example mesoscale holographic ‘influence’ maps measured in response to activation of an ensemble of V1 neurons (left) and an ensemble of LM neurons (right). Significantly modulated cells (including targets) are colored according to the amplitude difference between their calcium responses in photostimulation trials and their calcium responses in control (no photostimulation) trials. Non-modulated cells are displayed in gray. Experiment performed on a mesoscopic 3.1mm×3.1mm FOV recorded at 3Hz, in a CaMK2-tTA;tetO-GCaMP6s transgenic mouse injected locally with viral pAAV-hSyn-GCaMP6m-p2A-ChRmine-Kv2.1-WPRE at discrete locations. Photostimulation at 10×10ms pulses at 30Hz at Power=9mW. **b)** Distribution of post-stimulation modulation in targets (n=10) and follower cells. Post-stimulation modulation is defined as the difference between responses in photostimulation and no-power control trials. Follower cells are defined as reliably modulated cells across non-overlapping trial sets (see Methods). **c)** Example average calcium responses of individual modulated cells. Red/Blue: average response on photostimulation trials (n=100trials). In this example, photostimulation is combined with visual stimulation (gaussian modulated noise, see Methods). Black: average response on control trials (visual stimulation with no holographic photostimulation, n=100 trials). **d)** Modulation strength of the holographic perturbation on the non-targeted network. Modulation strength is defined as the average absolute value of the mean response across all follower cells, regardless of response polarity. Both excitatory and inhibitory modulations are therefore counted as positive values (see Methods). For LM photostimulation, the local network corresponds to all follower cells within LM and farther than 100µm from the targets. The distant network corresponds to all follower cells outside of LM. For V1 photostimulation, all target holograms were located in posterior V1. The local network corresponds to all follower cells within 100µm-1mm from the targets. The distant network corresponds to all follower cells outside V1. Local modulations strength: 0.39± 0.01 (mean± SEM), n=287 cells, 6 sessions, 2mice. Distant modulations strength: 0.28± 0.01 (mean± SEM), n=607 cells, 6 sessions, 2mice. Mann-Whitney p-value: 7.14×10^-28^ . Stimulated ensemble size varied between 10 and 60 cells stimulated per hologram per session. For each session, the hologram size was matched between LM and V1 stimulations. **e)** Net influence of the holographic perturbation on the non-targeted network. Net influence is defined as the average signed value of the mean response across all follower cells (see Methods). Local net influence: -0.19± 0.02 (mean± SEM), n=287 cells, 6 sessions, 2mice. Distant net influence: 0.1± 0.01 (mean± SEM), n=607 cells, 6 sessions, 2mice. Mann-Whitney p-value: 4.97×10^-26^. **f)** Inter-areal causal mapping of net influence across cortical areas computed from post-stimulation responses of follower cells. N=6 sessions, 2 mice. Area boundaries are defined using online two-photon retinotopic mapping (see **supp. Fig. 17** and Methods). Left: connectivity matrix of net influence. Right: inter-areal connectivity map. Arrow size is proportional to net influence. Blue or red indicate either inhibitory or excitatory influence respectively.

To illustrate how one can use the 2P holographic mesoscope to address key questions related to inter-areal processing despite the holographic FOV limitation, we sequentially targeted ensembles of neurons in different visual areas while recording the same set of visual cortical areas via translation of the mesoscope, taking advantage of the system’s targeting invariance to spatial translation. We could thus sequentially activate neural ensembles from four different cortical areas (V1, PM, RL, and LM) while simultaneously recording activity from six surrounding visual areas (**Fig. 2f**). In each of the four configurations, the system was translated to match the target stimulation area to the holographic FOV. The imaging FOV was then flexibly selected within the accessible 5mm ×5mm FOV to maintain a consistent FOV across the 4 configurations, as shown in Fig2f by the similarity in vasculature patterns between the 4 configurations. Photo-stimulation in each cortical area elicited robust response in the holographically targeted cells even while we maintained acquisition of physiological data from the entire FOV. This demonstrates that a 3D-SHOT based holographic mesoscope can achieve, in the same mouse, ensemble stimulation in any chosen visual area while concurrently recording neural activity from thousands of neurons in surrounding visual areas.

### Specific transmission of information across cortical areas

One of the major advantages of mesoscale two-photon holographic optogenetics should be the ability to specifically perturb the cortical network in one area and readout the impact of the perturbation in the population activity of thousands of neurons in multiple downstream areas, something not possible with conventional 2p microscopes. To test this, we stimulated ensembles of neurons in visual area LM while recording from neurons across 4-6 other visual cortical areas in a total 2.4 mm × 2.4 mm FOV at 4.5Hz (**Fig. 3a,b**). We then asked if activating different neural activity patterns in one cortical area (LM) induces differentiable patterns of modulation in downstream areas. To answer this question, we sought to decode, trial by trial, the identity of the holographically induced pattern in LM from the neural responses in the other visual areas (**Fig. 3a, Supp** **Fig. 11**). Indeed, we could properly predict the holographic pattern on each trial significantly above chance levels, despite excluding all measurements taken from LM itself (**Fig. 3b**). This indicates that photostimulation of even relatively modestly sized neural ensembles is sufficient to drive distributed and informative patterns of neuromodulation downstream at the cellular scale.

A key benefit of two-photon holographic optogenetics is the ability to co-activate ensembles of neurons based on their functional properties which may be critical for inter-area communication. We thus asked if we could transmit specific visual-like information across cortical areas by co-activating groups of visual cortical neurons that share orientation preference (**Fig. 3c-h**). In the mouse visual cortex, orientation preference shows minimal spatial clustering (Ohki and Reid 2007, Bonin et al., 2011, Ringach et al., 2016) and thus co-activating an ensemble of co-tuned neurons requires highly precise optical targeting. To test this capability, we measured orientation tuning curves of V1 neurons and then constructed various ensembles of neurons with shared orientation preference for 2p holographic stimulation (**Fig. 3d,e**). Indeed, we could successfully photo-activate groups of co-tuned neurons in V1 (**Fig 3f, Supp. Fig 11**) while recording activity from surrounding areas. To test whether our holographic stimulation generated functionally specific patterns of neural activity, we trained a decoder to discriminate the visual responses evoked by static gratings across four orientations (**Fig. 3g, Supp. Fig 12**). In the test phase, we probed this decoder, trained only on visual responses, to classify neural activity patterns evoked purely by photostimulation of co-tuned ensembles, i.e., in the absence of any visual stimulus. The decoder was able to classify holographic photo-stimuli above chance across both photostimulation targets and across all recorded neurons (**Supp. Fig 12, Fig. 3h**). Importantly, even when only decoding from neurons in downstream extrastriate areas, the decoder correctly classified holographic trials according to the orientation preference of the photostimulated ensemble, slightly but significantly above chance (**Fig. 3h**). This result demonstrates that the system both writes in and transmits visually relevant information across cortical areas.

### Photostimulation across a mesoscopic field-of-view with random-access 3D-SHOT

Activating neural ensembles in one area and recording from adjacent areas opens a new class of causal paradigms and is sufficient for many experiments. However, creating comprehensive inter-areal functional connectivity maps requires stimulating groups of neurons in different cortical areas within the same experiment, preferably rapidly and randomly interleaved across trials. The initial holographic mesoscope we described above does not achieve this, as the photostimulation area is fixed within the FOV owing limitations of the spatial light modulator. To overcome this challenge, we sought to engineer a random-access holographic photostimulation path that would allow flexible jumps to distant neural targets. Our solution incorporated XY galvanometer scanners conjugated to the back aperture plane, enabling flexible and fast repositioning of the SLM FOV anywhere within the mesoscopic FOV (**Fig. 4a**). Implementing this solution using exclusively off-the-shelf optics proved challenging due to the limited availability of large-aperture components and the need to satisfy multiple design constraints simultaneously. First, the galvanometers needed to be conjugated to both the objective back aperture and the spatial light modulator (SLM). Second, due to the large beam size at the back aperture, even when using oversized galvanometers, the beam still required significant reduction, and this beam reduction had to maintain the resolution and optical specifications of the 3D-SHOT path at the center of the FOV. Third, there were substantial steric constraints, given the use of large 3-inch optics within the compact space of the mesoscopic breadboard. Finally, the imaging and holographic paths had to be not too far from coplanarity to maintain the 3D capability of the system. We thus designed an optical configuration that combined a compact pre-scanning beam reducer in collimated space, large 30mm XY galvanometer scanners, and an f-theta scan lens with a shortened focal length to restore the beam size at the back aperture (**Fig. 4b, Supp. Fig13**). All optical components could be obtained as stock items from major optics vendors, avoiding the need for customized optical elements.

To maintain high diffraction efficiency across the FOV, we tiled the mesoscopic FOV into 81 quadrants of 350 µm × 350 µm, each centered on a unique (X, Y) galvanometer position. This strategy enabled consistent patterned photostimulation across a 3.2 mm span (**Fig. 4c**), representing nearly a tenfold expansion of the accessible photostimulation field. We validated that the random-access holographic configuration retained the axial resolution performance of the mesoscale 3D-SHOT system, as presented above, at the center of the FOV (mean axial FWHM = 34µm, n =154 holograms, **Fig. 4d**). We characterized the axial resolution and the hologram intensity as the galvo is scanning across the mesoscopic FOV (**Figs 4e,f** **and supp. Fig. 14**). Hologram intensity decreased by only ∼2.5-fold from the center to the corner of the 3.2 × 3.2 mm FOV, requiring only a 1.5-fold adjustment in photostimulation power to compensate. Hologram optical axial resolution decreased across the mesoscopic FOV, remaining below 50 µm over more than two-thirds of the area, with a worst-case edge average axial resolution of 56 µm. We characterized the axial field curvature across the mesoscopic FOV and found a maximum relative shift of 85 µm between the flattened imaging plane and the holographic planes, a variation that remains well within the SLM’s defocusing correction range (**supp. Fig. 14c**). Finally, as done above, we characterized the functional point spread function (PPSF) at an off-center galvanometer position. While near single-cell resolution was achieved at the lowest photostimulation powers, effective functional axial resolution degraded more rapidly at higher powers (∼ 82µm FWHM at the highest power probes beyond saturating power, **supp. Fig. 16**). The measured values remained at the upper end, but within the range of reported PPSF values for patterned two-photon optogenetics. We next demonstrated that random-access 3D-SHOT could reliably activate holograms from arbitrary locations within the 3.2 × 3.2 mm² mesoscopic photostimulation field of view (**Fig. 4g,h**). By dynamically updating the holographic phase mask for each selected galvanometer (X, Y) position, we successfully targeted and stimulated user-defined groups of neurons across distinct cortical areas within the same experiment, in a randomized trial-by-trial manner.

### Causal mapping of functional connectivity across cortical areas

One of the major advantages of mesoscale two-photon holographic optogenetics is the ability to map functional interactions between multiple areas simultaneously with near cellular resolution. To test this, we stimulated ensembles of neurons in either primary visual area V1 or visual area LM, while recording from neurons across more than seven other cortical areas (**Fig. 5a, supp** **Fig. 17**) in a total 3.1 mm × 3.1 mm FOV recorded at 3Hz. We detected downstream neurons both activated and suppressed upon photostimulation (**Fig. 5c**), located both within the same visual area as the targets and at distances of hundreds of microns to millimeters away. For connectivity analyses, we defined follower cells as putative strong and highly reliable connections, whereas modulated cells encompassed all significantly responsive cells to photostimulation, including both strong and weaker connections (see Methods). As a control, we compared the distribution of follower cells in photostimulation trials to that of statistical ‘follower cells’ identified in no-power (0 mW) control trials (Fig. 5b), confirming that holographic stimulation induces postsynaptic modulation above the false positive rate (**Fig. 5b, Supp** **Figs. 18,19**). Importantly, we found that these downstream modulations were consistent across trial splits, suggesting a deterministic impact of the photostimulation on the network despite the variability of ongoing spontaneous activity (**supp. Fig. 21**).

We first quantified the strength of holographic perturbations on both local and distant networks (**Fig. 5d**). We defined the local network as cells located in the photostimulation region 100-1000 microns from the nearest target, and distant neurons as all neurons in other areas. We computed the modulation strength as the sum of the absolute modulation amplitudes of follower cells divided by the number of follower cells (see Methods) to quantify the average amplitude of photostimulation-induced responses independently of polarity. We found that holographic perturbations produced significantly stronger modulation within the targeted cortical area compared to distant areas (Mann-Whitney p-value: 7.14×10^-28^), consistent with the high density of local recurrent connections in visual cortex (Rossi, Harris, and Carandini 2020; Inácio et al. 2025). Next, we examined the net modulation impact of holographic photostimulation in V1 and LM. In both areas, we observed a significant inhibitory effect on the local network (**Fig. 5e**), confirming and extending previous findings (Chettih and Harvey. 2019, Oldenburg et al., 2024, Rowland et al., 2023, LaFosse et al. 2024). Notably, in downstream areas the net effect in follower cells was excitatory (**Fig. 5e**). To more clearly reveal these long-range modulatory effects, we constructed functional connectivity maps based on the responses of follower cells to holographic perturbations per area, both for stimulation in V1 and in LM (**Fig. 5f**). While an exhaustive mapping of functional connectivity in the visual cortex is beyond the scope of this manuscript, these initial perturbation experiments suggest emergent reliable connectivity patterns (**supp. Fig. 21**). Activating neural ensembles in LM elicited strong inhibition in areas AL and RL, but drove excitation in V1 and PM. Similarly, stimulating V1 ensembles across sessions produced a net inhibitory effect in RL, while LM, M, and MMP emerged as strongly activated targets (**Fig. 5f**). The expected bidirectional connectivity between V1 and LM was recovered in the net effect analysis based on highly reliable follower cells and was also evident in absolute connectivity measures (**supp. Fig. 22**), suggesting that excitatory and inhibitory influences co-exist between these areas and may cancel out at the population level when averaging across large cell populations that include weaker connections (**supp. Fig. 20**).

### Near-simultaneous activation of remote neural ensembles in different cortical areas

Finally, we took advantage of the high speed of the galvanometer scanners and the rapid refresh rate of the SLM to synchronize the phase mask to the galvo position for every optogenetic stimulation pulse, enabling the interleaving of two remote holograms (**Fig. 6**). With a 10×10ms pulse stimulation at 10Hz, the two remote neural ensembles would be coactive within a 950ms time window. Using this strategy, we stimulated a single neural ensemble in V1, a separate single ensemble in LM, and then both ensembles near-simultaneously (V1+LM). The resulting average post-stimulus histograms showed that, in the latter case, both ensembles were effectively activated simultaneously (**Fig. 6b**). Lastly, we leveraged this experiment to demonstrate how this fast scan capability can enable new experimental paradigms for investigating inter-areal computations. In particular, it is unknown whether the functional impacts of two long range projection pathways that converge on a single downstream area will summate linearly or non-linearly. To address this question, we compared the net impact of photo-stimulating ensembles in V1 and LM simultaneously to the predicted sum of the effects when each was stimulated in isolation (**Fig. 6c**). We found that the net effect of near-simultaneous co-activation yielded modulations on individual downstream neurons that were less than the predicted sum, potentially indicative of normalization mechanisms (Carandini and Heeger, 2012). In V1-modulated cells, we also identified a small number of highly modulated ‘amplifier’ cells that showed the opposite trend and exhibited supra-linear summation. Overall, these findings illustrate how the holographic 2P mesoscope platform enables new classes of experiments that have until now been completely inaccessible with existing technologies.

**Figure 6.**
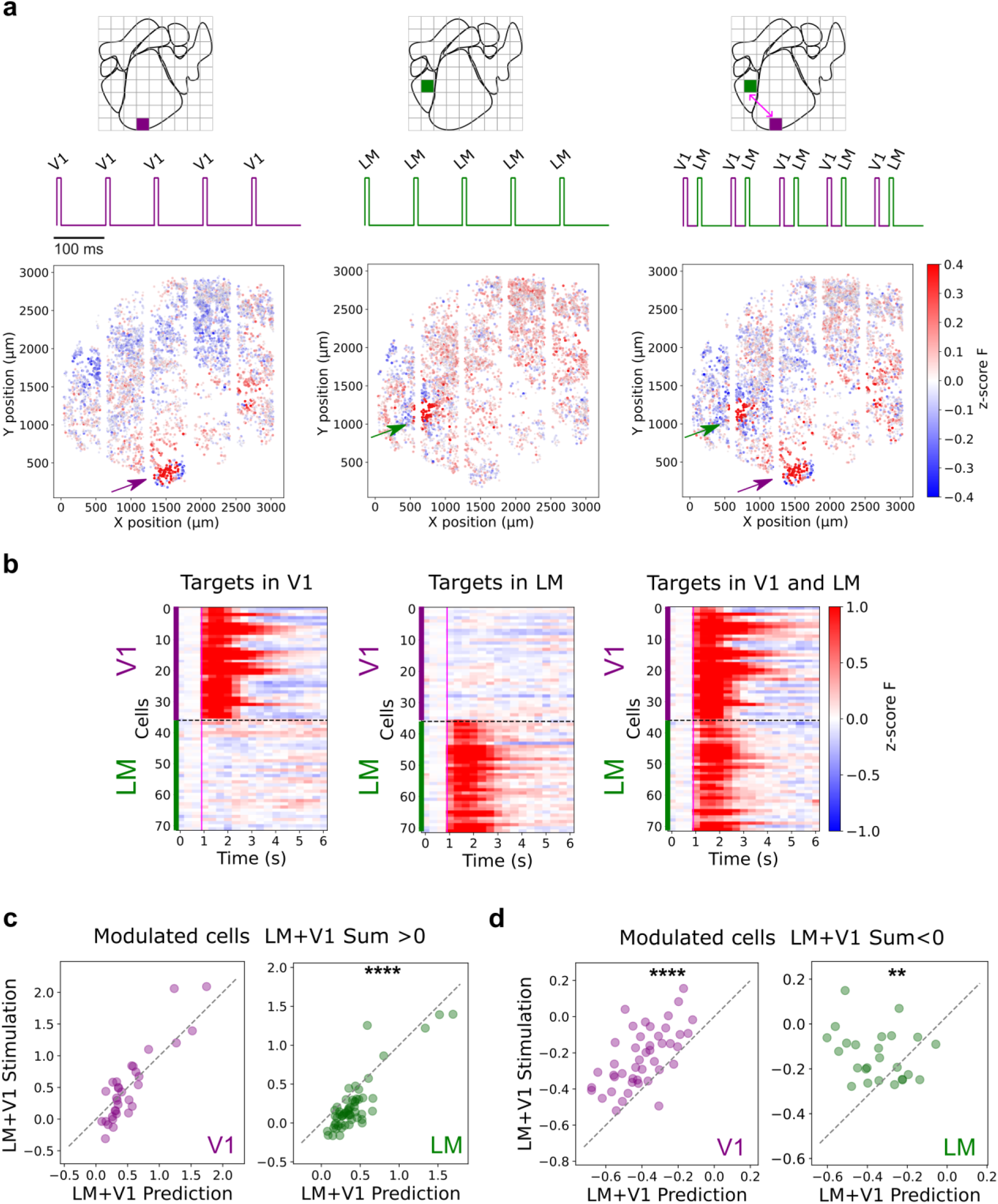
Near-simultaneous mesoscopic photostimulation of distant neural ensembles. **a)** Top: Principle of near-simultaneous mesoscopic photostimulation with hologram interleaving. 10Hz pulsed stimulation in V1 with Hologram 1 (left), 10Hz pulsed stimulation in LM with Hologram 2 (center), and fast interleaved 10Hz Hologram1 and Hologram2 stimulation (right) at 52.6 Hz galvanometer interleaving rate (i.e., 19ms between end of stimulation pulse for hologram 1 and start of stimulation pulse for hologram2). Bottom: Mesoscopic influence maps for single hologram perturbation in V1 (left), single hologram perturbation in LM (center) and joint interleaved holograms in V1 and LM (right). All cells detected in the FOV are displayed with a color displaying their relative response to photostimulation. Arrows indicate holographic perturbations. 35 cells photostimulated in LM and 35 cells photostimulated in V1. **b)** Average post-stimulus calcium responses when stimulating one single neural ensemble in V1 (left), one single neural ensemble in LM (center) and both neural ensembles near-simultaneously (right). Experiment performed on a mesoscopic 3.1mm×3.1mm FOV recorded at 3Hz, in a CaMK2-tTA;tetO-GCaMP6s transgenic mouse injected locally with viral pAAV-hSyn-GCaMP6m-p2A-ChRmine-Kv2.1-WPRE at discrete locations. Photostimulation at 10×10ms pulses at 10Hz at P=12.5mW. **c-d)** Scatter plots showing for every modulated cell by V1 perturbation (purple) or LM perturbation (green), the calcium response to the joint V1+LM holographic photostimulation against the predicted calcium response computed from summating responses to single V1 stimulation and LM stimulation. Given the non-normality of the distribution, responses are split whether the predicted sum V1+LM is positive (c) or negative (d). In c) Left: n=32 cells modulated cells from V1 stimulation, fraction of sublinear cells 0.59, p-value Wilcoxon signed-rank test: 0.50. Right: n=56 cells modulated cells from LM stimulation, fraction of sublinear cells 0.91, p-value Wilcoxon signed-rank test: 4.49×10^-9^. In d) Left: n=45 modulated cells from V1 stimulation, fraction of sublinear cells 0.044, p-value Wilcoxon signed-rank test: 3.74×10^-8^. Right: n=25 modulated cells from LM stimulation, fraction of sublinear cells 0.28, p-value Wilcoxon signed-rank test: 0.001815.

## Discussion

We designed, built and validated the first multiphoton holographic mesoscope. This technology is the first to offer targeted photostimulation approximating single-cell resolution, with simultaneous large-scale recording of neural activity across several cortical areas in a contiguous mesoscopic FOV. We demonstrated that engineering a scanning module within the photostimulation system increases the accessible photostimulation FOV by an order of magnitude and enables near-simultaneous activation of spatially remote neural ensembles. We established the new system’s ability to co-activate user-defined groups of neurons, selected based on their spatial or functional properties, while simultaneously recording neural activity from several downstream areas. Notably, we revealed that specific functionally relevant information can be holographically written in and propagated to downstream networks. We could identify individual downstream neurons that displayed time-locked excitatory or inhibitory responses hundreds or even thousands of microns away. We showed that this approach provides a foundation for constructing the first inter-areal cortical connectivity maps based on targeted causal perturbations. Overall, this demonstrates that one can use this new technology to probe how both local and long-range functional interactions relate to specific neuronal computations, all within the same experiment.

We developed the mesoscale read/write platform based on a commercially available 2p mesoscope (Thorlabs) that is a widely used mesoscope platform (Fahey 2019, Froudarakis 2020, Orlova 2021, Kanamori 2022, Pandey 2022) and that has enabled a number of other recent technological advances (Lu 2020, Demas 2021, Tsyboulski 2018). We expect that adapting a commercially available system will facilitate the adoption of the technology across both individual labs and new neurotechnology platforms. In SLM-based holographic systems, the size of the holographic FOV is fundamentally limited by the pixel count and pixel size of the spatial light modulator (SLM). Ongoing advances in high-resolution SLM technology — including increased pixel density and reduced pixel size—may eventually enable large-FOV photostimulation using a single SLM. However, such approaches will continue to be limited by reduced power efficiency at high diffraction angles and potential inter-pixel crosstalk, both of which can constrain the effective usable FOV. Our random-access 3D-SHOT strategy is inherently compatible with and can further amplify the benefits of these future SLM developments, enabling even larger and more flexible photostimulation fields. The concept of expanding the SLM FOV using galvanometer scanning has been previously explored for holographic imaging (Yang et al., 2015) and proposed conceptually for photostimulation (Sun et al., 2019). However, this is the first implementation of such an approach at the mesoscale, integrated within a constrained multi-4f holography and temporal focusing layout (3D-SHOT). These prior studies have demonstrated random-access holographic imaging over 0.15 mm² (Yang et al., 2015) and structured illumination across 1.64 mm². In contrast, our platform enables two-photon photostimulation across more than 10 mm², while simultaneously recording from thousands of neurons distributed across multiple cortical areas.

While recent studies have initiated the use of two-photon optogenetics to probe inter-areal communication, they remain constrained to interactions between immediately adjacent cortical regions (Rowland et al., 2021; Fisek et al., 2023). One recent study investigated inter-areal communication using two-photon optogenetics on a conventional two-photon microscope by imaging across the boundary of two contiguous cortical regions (Rowland 2021), and another more recent study also enabled inter-areal interrogation between two contiguous areas, using a low NA 10x objective to achieve a 2mmx2mm FOV imaging FOV (Fisek 2023). In terms of imaging capability, our platform enables recording of 18-fold more cortical surface than Rowland et al., and significantly higher axial resolution with 4 times more cortical surface compared to Fisek et al. Importantly, the full 5mmx5mm nominal FOV of our platform enables random access selection of imaging ROIs anywhere within the 25mm^2^ FOV. In terms of photostimulation capability, our secondary large FOV design expands photostimulation FOV by tenfold compared to any other published work. A limitation of the current design is the trade-off between resolution and large field-of-view access. However, the observed resolution degradation is primarily due to off-axis scan aberrations and could be substantially mitigated with a custom large-aperture scan lens, offering a potential route for improving performance in future implementations.

Our large-scale holographic perturbation experiments provide a first demonstration that mesoscale two-photon optogenetics can be used to map functional connectivity across multiple cortical areas with near-cellular resolution. By selectively stimulating neural ensembles in V1 and LM while recording activity across the dorsal visual cortex, we revealed distinct patterns of excitatory and inhibitory influence that vary with both source and target area. Notably, while local networks showed net suppression, downstream areas often exhibited net excitation, suggesting divergent circuit motifs within and across areas. These findings establish mesoscale holographic optogenetics as a powerful approach for dissecting inter-areal functional influences with high spatial precision, enabling systematic investigation of long-range cortical interactions.

While there is substantial work detailing how individual cortical areas represent and encode information, how neural representations are transformed across cortical areas remains elusive. Most studies addressing cortico-cortical communication combined multi-site electrode recordings with correlative analyses (Bressler 2011, Semedo 2019, Siegle 2021), or electrical or one-photon optogenetic stimulation. However, electrical microstimulation provides limited spatial control and often (and perhaps preferentially) activates axons of passage (Histed 2009). One-photon optogenetic stimulation has cell-type specific control but cannot readily activate functional defined subsets of neurons (such as neurons with the same feature preference). Molecular approaches such as c-fos and TRAP (Liu et al., 2012; Guenthner et al., 2013; Girasole et al., 2018), when combined with one-photon optogenetics, offer some degree of functional specificity but within a far less flexible framework than two-photon optogenetics, as they do not permit the selection of cells based on multiple functional features. Neither microstimulation nor 1P optogenetic stimulation can recreate precise spatiotemporal sequences of activity, nor can they drive different user-defined firing rates into different neurons to recreate specific population activity vectors that might preferentially drive cortico-cortical communication (Semedo 2019). 2p holographic optogenetics can achieve these goals (Bounds 2017, Mardinly 2018), and this new 2p holographic mesoscope can implement these perturbations while simultaneously measuring both local and downstream neural activity. Thus, 2p mesoscale holography is set up to experimentally test major hypotheses and theories of interareal communication, such as how cortico-cortical feedback implements generative models of the external world such as in the framework of predictive coding (Rao and Ballard 1999, Bastos et al. 2012, Markov and Kennedy 2013, Hertag and Clopath 2022).

The 2P holographic mesoscope could also guide the design of more effective brain-machine interfaces (BMIs, Ersaro 2023). Access to intermediate processing stages between targeted neural perturbations and behavioral output will be essential for interpreting cases where stimulation fails to produce the expected outcome. Understanding and optimizing signal propagation across areas may represent a critical strategy for enhancing the efficacy of holographic interventions in driving behavior (Marshel et al., 2019; Bounds and Adesnik, 2025; Rowland et al., 2023; Russel et al., 2024).

Finally, the large accessible FOV of the mesoscope will help scale two-photon optogenetics to species with larger brains such as primates. This platform could also enable new experimental paradigms in smaller model organisms, such as larval zebrafish, with minimal beam shaping to adapt the holographic spot size—allowing targeted photostimulation in the central nervous system while simultaneously recording from both central and peripheral systems. It could even allow photo-stimulation and recording from multiple small animals simultaneously, such as flies or worms, engaged in social behaviors. Overall, we expect mesoscale two-photon holographic optogenetics to become a key technology in systems neuroscience because it can establish whole new classes of perturbative experiments to motivate and test previously untestable theories of brain function.

## Acknowledgements

We thank Paulo Chaves (Thorlabs), Stephen Prieur (Thorlabs) and Nikhil Bhatla for technical assistance. We thank the members of the Adesnik lab and Eirini Papagiakoumou for comments and discussions.

This work was funded by NIH grants R01NS128772, UF1NS107574, R01MH117824 and R01EY023756. H.A. is a Chan Zuckerberg Biohub Investigator. This work was supported by a Weill Neurohub fellowship to L.A and by the Burroughs Wellcome Fund with a Career Award at the Scientific Interface (1243586) to L.A.

## Author contributions

L.A. and H.A. conceived of the project. L.A. designed and built the 2p holographic system with assistance from Thorlabs. L.A and U.K.J characterized the system. L.A., H.S. and U.K.J. performed experiments. M.O contributed software. L.A. and H.A. wrote the paper with assistance from H.S. and U.K.J.

## Methods

### Animals

All experiments on animals were conducted with approval of the Animal Care and Use Committee of the University of California, Berkeley. All experiments were performed in mice of both sexes, aged two months and older. Three transgenic mouse lines were used: either Ai203 transgenic lines (Bounds et al., 2023) crossed in-house with Vglut1-Cre mice, or double (CaMK2-tTA;tetO-GCaMP6s) or triple (emx1-Cre;CaMK2-tTA;tetO-GCaMP6s) mice, although no Cre-dependent viruses were used in this study, except in Supp. Fig 19. Transgenic mice were obtained by crossing the corresponding lines in-house (JAX stock# 005628, Jax stock# 003010 and Jax stock# 024742). Mice were housed in groups of five or fewer in a reverse light: dark cycle of 12:12 hours. Experiments were conducted during the dark phase.

### Surgery

Mice were anesthetized with isoflurane (2%) and given buprenorphine as an analgesic (0.05mg/kg) and dexamethasone (2mg/kg) to reduce brain swelling. Mice were then placed in a stereotaxic frame (Kopf) over a heating pad. The scalp was removed, the fascia pushed to the sides, and the skull lightly scratched for better cement adhesion. A 4 or 5mm craniotomy centered around V1 was made using a biopsy punch (Robbins Instruments) and/or a dental drill (Foredom). Bleeding was controlled with cold phosphate-buffered saline and Gelfoam (Pfizer Inc.) For mice that required viral injection (double and triple transgenics), the virus preparation (pAAV-hSyn-GCaMP6m-p2A-ChRmine-Kv2.1-WPRE or AAV9-CAG.DIO.ChroME2s-FLAG-ST.P2A.H2B.mRuby3.WPRE.SV40) was injected using a beveled glass pipette at 4-5 locations across visual cortex. A cranial window, made of 7 mm and 5 mm diameter glass coverslips or 6 mm and 4 mm diameter glass coverslips, for 5 and 4 mm craniotomies respectively, was then placed onto the craniotomy and held in place with Metabond (C&B). Finally, a titanium headplate was fixed to the skull with Metabond (C&B0 and Ortho Jet (Lang). All craniotomies were performed on the left hemisphere of the mouse brain. Animals were allowed to recover in a heated recovery cage before being returned to their home cage.

### 2P-RAM mesoscope with a temporally-focused holographic path

The mesoscale read/write platform was custom-built around a 2P random-access fluorescence mesoscope previously described in detail (Sofroniew 2016) and now commercialized by Thorlabs Inc. The system had a nominal 5mm FOV accessed through a 0.6 NA objective that can be rapidly and flexibly scanned-across with a set of four conjugated scanners, one 12kHz resonant scanner and three large-angle galvanometers. The system was also equipped with a voice-coil remote-focusing module (Botcherby 2008) for 3D multi-plane imaging. In addition, the mesoscope was built on a vertical breadboard and the whole system was motorized in 4 dimensions (X, Y, Z translation and one axis of rotation) to provide maximal flexibility for the user. Since the microscope moved with respect to the incoming laser beam, the beam was injected through a six-stage periscope for alignment invariance.

The 2P mesoscale imaging path relied on a Ti:sapphire laser (Mai Tai, Spectra Physics) used at 920 nm for calcium imaging. External power control was achieved through a Pockels cells (Conoptics, Inc). A prism-based group delay dispersion compensation module (Akturk 2006) was installed to compensate for dispersion introduced by the mesoscope imaging path optics (estimated at about 25,000 fs^2^, Sofroniew 2016). The 2P holographic path relied on a 1030 nm femtosecond fiber laser (Aeropulse 50, NKT Photonics) with internal power control.

The holographic path was built on a 33”” × 24”” breadboard attached to the main mesoscope frame with three custom mounts (see Supplementary CAD). The holographic beam was merged with the imaging beam with a 1000 nm cutoff dichroic (DMSP1000R, Thorlabs) and rigorously co-aligned with the imaging beam through the system’s periscope to maintain alignment invariance with translation. The beam was then injected onto the 3D-SHOT optical path (see Supplementary partlist and supplementary CAD) with specific care to maintaining beam horizontality to achieve alignment invariance with rotation (see Supplementary note).

To expand the holographic FOV while preserving hologram resolution at the center (Fig. 4), a compact 1:2 Galilean telescope was inserted into a collimated section of the optical path after the SLM zero-order (between lenses L5 and L6). Large-aperture 30 mm XY galvanometer scanners (QS30XY-AG, Thorlabs) were added to maintain optical conjugation with both the SLM plane and the objective’s back aperture. The original L6 lens was replaced with a 100 mm f-theta scan lens (FTH100-1064, Thorlabs) to restore axial hologram resolution, and the recombination dichroic (D3) was replaced with a larger version to accommodate the large beam scanning. Finally, the remote-focusing voice coil used for imaging was adjusted to achieve coplanarity between the imaging and holographic planes at the center of the FOV.

The imaging path hardware was controlled with ScanImage software (Vidrio Technologies,LLC). Custom Matlab code was used for control of the photostimulation path hardware, synchronization with imaging, synchronization of galvo scanners and SLM phase masks and control of the visual stimulation.

### SLM nominal FOV calculation

The maximal lateral deflection from the SLM is achieved for the smallest effective ‘slit’ size on the SLM which would correspond to 2 pixels (the pattern on the SLM would then be a black and white grating pattern with a pixel-sized stripe pattern). Based on the grating equation and taking into account that the effective pixel size at the objective back aperture is magnified by the microscope optics, the nominal maximal FOV is given by:

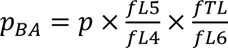, where λ is the laser wavelength, 𝑓_𝑜𝑏𝑗_ is the objective focal length and 𝑝_𝐵𝐴_ is the effective SLM pixel size at the back aperture. 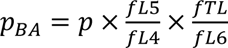 where p is the SLM pixel pitch and f_L4_ f_L5_, f_L6_ and f_TL_ the focal lengths of L4,L5,L6 and the tube lens respectively.

### Hologram characterization and system calibration

To characterize the generated holograms, 2P fluorescence was recorded from a thinly coated fluorescent slide with a substage objective (×4 Zeiss) coupled to a camera (Basler) with a f=125 mm achromatic doublet which provided a 1.3 × 1.3 mm characterization FOV. The whole substage system (fluorescent slide, objective tube lens, camera) is mounted on an XYZ motorized stage (Sutter Instruments) to enable hologram characterizations without moving the mesoscope. For successful multiphoton optogenetics experiments, precise co-alignment of the imaging and the photostimulation beams is required. To achieve this, hundreds of holograms were generated in 3D and any arbitrary curvature or inhomogeneity on the imaging remote-focusing planes or the SLM planes was taken into account in the computation of the hologram to real space transform matrix (Supp Fig. 2), in a fully automated procedure (Oldenburg 2024). The holographic FOV was then remapped onto the entire mesoscopic FOV so that photostimulation targets could be flexibly selected in any imaging ROI configuration. Given that the nominal accessible FOV of the mesoscope spans 5 mm × 5 mm and acquisitions are performed in a random-access manner, appropriate coordinate transformations are required to convert relative positions within a given imaging ROI to absolute spatial locations within the full mesoscopic FOV. The recorded holograms were also used as a reference to correct for diffraction efficiency fall-off and deliver homogeneous powers across the FOV *in-vivo*. To characterize hologram position invariance with translation and rotation, 2D point-cloud holograms were generated to “burn holes” i.e photobleach a fluorescent slide. The 2P fluorescence signal on the slide was then recorded with the mesoscope photomultiplier tubes (PMTs).

To characterize the impact of galvanometer scanning on hologram calibration for large FOV photostimulation, we first recorded 3D point-cloud holograms at the (0 V, 0 V) galvanometer configuration (central FOV), and then re-recorded the same holograms while varying galvanometer positions (supp. Fig. 15) and translating the substage system accordingly. We found that the absolute hologram coordinates shifted linearly with galvanometer deflection, indicating that the SLM-to-real-space calibration established at the central FOV can be extended to other mesoscopic FOVs with a simple global offset adjustment. For each off-axis FOV, lateral offsets were determined by targeted photobleaching on a fluorescent slide (hole burns).

### Visual stimulation and retinotopy

Visual stimuli were displayed on a 2048 × 1536 Retina iPad LCD monitor (Adafruit Industries) placed 10 cm from the right eye of the mouse and computed using Psychtoolbox-3 and custom MATLAB code. Visual stimuli were presented to map retinotopy and to identify functionally-defined ensembles (Fig3d-h). In Figs 3a-b,4 and 6 no visual stimulus was presented to the mouse during simultaneous imaging and photostimulation experiments. In the functional connectivity experiments (Fig. 5), half the datasets were collected without concurrent visual stimulation and the other half with simultaneous visual and holographic stimulation.

Retinotopy mapping was used primarily to identify visual cortical area boundaries. To map retinotopy, we adapted the procedure used in Marshel et al., 2011 and Zhuang et al., 2017 for the 2-photon mesoscope (supp. Fig. 17). Briefly, we imaged a 3.1 x 3.1mm FOV 100-200um below the dura at 3Hz when mice were viewing noise bars drifting in the horizontal or vertical directions. The bars were 10’ in width and spanned the vertical or horizontal extents of the screen respectively. The noise was drawn from a gaussian distribution with spatial frequency 0.03 cycles/deg and temporal frequency 3 cycles/deg. Each type of bar swept the screen 30 times with a period of 9s, for a total stimulation duration of 10 minutes. We smoothed the recorded 2-photon calcium frames with a 4-8 pixel square filter and downsampled them by 4-8x. For each pixel in the reduced image, we computed the preferred azimuth and elevation by identifying the average horizontal and vertical positions of the bar at which the pixel’s response was maximal. This generated azimuth and elevation preference maps for the full FOV. We then computed the sign of the gradients between the two maps to generate a visual field sign map (Sereno, et al., 1994) and used sign flips in this map to identify area boundaries as outlined in (Garrett, et al., 2014, Juavinett, et al., 2017).

### In vivo mesoscale two-photon imaging and photostimulation

For *in vivo* 2-photon mesoscale calcium imaging in vivo, mice were head-fixed and allowed to run on a freely moving circular treadmill. Mesoscale Imaging ROIs were set to 2.4 × 2.4 mm to 3.6 mm × 3.6 mm FOVs with recording rates between 3 and 5Hz. The imaging laser was set at 920 nm. Opsin expressing cells were automatically segmented online based on either the green channel signal (Ai203 mice or transgenic mice virally injected with ChRmine-GCaMP6m) or the red nuclear mRuby2 (virally injected mice with ChroME2s-mRuby2). To minimize the photostimulation artefact induced by the stimulation laser on the imaging, we use an Arduino Mega 2560 board to gate the photostimulation laser on the imaging resonant galvanometer and only stimulate at the edge of the resonant galvanometer scan. The corresponding edge stripes are excluded from analysis. Despite this gating strategy, given the large detection etendue of the 2P-RAM mesoscope, we have noticed that multi-target photostimulation generates an artefact on the imaging outside of the gating window when using mice expressing red fluorescent proteins. For these experiments (Fig3), we have conservatively subtracted post-stimulus frames for analysis and only used the tail of the calcium traces for analysis. This caveat is completely solved when using mice not expressing any red fluorescent protein (Figs4-6) and the full trace can be used for analysis. Experiments in Figs2,3 were performed with a standard Hamamatsu H10770A photomultiplier tube (PMT). Experiments in Figs4,5,6 were performed with a gated PMT (H11706-40, Hamamatsu) used in non-gated mode.

### Optical physiological point-spread function measurements

To estimate the PPSF in the Ai203×VglutCre mouse, we first identified photoactivatable neurons near the center of the holography FOV and selected 8 targets for multi-target stimulation. We generated 50 8-target holograms around and including the central holograms, 17 for the lateral PPSF and 33 for the axial PPSF. For the lateral PPSF, each hologram was offset in the [-40,40] um range with 5um increments along the x-axis. For the axial PPSF, each hologram was offset in the [-80,80] um range with 5 um increments along the z-axis. Each hologram was illuminated with 5 x 5ms pulses @ 30Hz and 25mW/target, with 10 repeats per hologram. The responses of the targeted neurons were averaged over each condition’s repeats to compute the PPSF.

For the PPSF experiment sessions in mice virally expressing the ChRmine opsin, we first ran a photoactivatability experiment at a series of powers (range 0.5-10 mW/target) on a set of 40-80 putative opsin-expressing neurons with the multi-target hologram centered, i.e., lateral and axial offsets set to 0. These experiments were performed with the 2P-RAM configuration with galvos set at the central (0,0) position, which we have shown generates similar optical PSFs to the no galvo 3D-SHOT implementation (Fig. 4). The stimulation was delivered as a train of 10 x 10 ms pulses at 30 Hz. We computed the power-response relationship for individual neurons and selected 10-20 neurons activated at intermediate powers and exhibiting a high dynamic range of power responses. We then performed the PPSF experiment on this subset of neurons with the corresponding holograms shifted laterally (range -40 to 40 µm in 5 µm increments) or axially (range -80 to 80 µm in 10 µm increments), and at low (0.5-2 mW/cell), intermediate (2-5 mW/cell), and high (5-10 mW/cell) powers, repeated 10 times for each offset/power combination. For each neuron’s ROI, the extracted calcium fluorescence traces were baseline-subtracted, and the evoked response was computed by averaging the deltaF in a 1.5 s window following stimulation onset for each condition. Computed responses at different spatial offsets were also normalized to the centered (zero offset) condition at each power and averaged across neurons. We then fit a weighted gaussian to the mean spatial profile of responses to compute the PPSF metrics at each power.

### Multi-area targeting with single field-of-view 3D-SHOT

For the multi-area holography demonstration in Figure 2f, we first identified area boundaries using retinotopy, and picked 4 visual cortical areas (V1, PM, RL, and LM) for same-session sequential targeting. For each area, we moved the objective along both lateral axes until the holographic FOV (fixed with respect to the objective) was positioned over the center of the area. We then adjusted the 2P-RAM scan fields to image a contiguous 3.1 x 3.1 mm area covering a fixed portion of the window using the vasculature as fiduciary markers. As a result, at the beginning of each area’s experiment, the imaging FOV was identical, but the holography FOV covered a different cortical area. In each experiment, we identified 10-15 of the most photoactivatable neurons with sequential single-neuron targeting. We then targeted these neurons (V1-13, PM-12, RL-11, LM-12 neurons) in each area using a multi-target hologram, driving them with a single 250ms pulse and 80 mW per target. This configuration requires repositioning the mouse and post hoc registration of the imaging FOV, making it well suited for experiments that involve probing different cortical areas across separate sessions.

### Holographic stimulation of random multi-target ensembles

For the targeted photostimulation experiments presented in Figure 3a-b, we used Vglut1-Cre;Ai203 mice. We recorded 4 sessions from 2 female mice (n=2 sessions each). The holographic FOV was placed over LM. The imaging FOV encompassed V1, LM, RL, AL, A and AM (3000 X 1800 µm2). Each target received 50mW, and each stimulation consisted of 10 pulses of 10 ms duration at 16 Hz. In each session, 4 distinct holograms were used in addition to a zero power condition, with 80 trials per condition.

### Decoding photostimuli pairs

To discriminate between pairs of distinct photostimulation patterns, a gradient boosting decision tree algorithm was used. The classifier was trained on 70% of the trials (n=112 trials per pair labeled as either photostimulus 1 or photostimulus 2) and tested on the remaining trials (n = 48 trials). The decoding accuracy was defined as the number of correctly predicted trials over the total number of test trials. For each pair of photostimuli, the final reported decoding accuracy result was the average across 10 independent draws (cross-validation). For the shuffled control decoding, the classifier was trained on randomly mislabeled trials and tested on the 30% held-out trials.

### Holographic Stimulation of Functionally Defined Ensembles

For co-tuned ensemble stimulation experiments presented in Figure 3, we used Vglut1-Cre;Ai203 mice (Ai203 refers to a transgenic line expressing the transgene TITL-st-ChroME-GCaMP7s-ICL-nls-mRuby3-IRES2-tTA2; Bounds et al., 2023). We recorded 11 sessions from 2 female mice (5 and 6 sessions each mouse). The holographic FOV was placed over V1 (white rectangle in Figure 3a). The imaging FOV encompassed V1, LM, RL, AL, PM and AM (2400 X 2400 µm^2^). The imaging and holographic FOV was fixed throughout each recording session.

To holographically stimulate neural ensembles defined by their visual response properties, each recording session consisted of 3 steps. In Step 1, we recorded visual responses of neurons in the FOV while the mouse viewed various images on the monitor. To gauge the orientation preference of each neuron, we presented static gratings in 4 different orientations (0, 45, 90, 135°), at a contrast of 0.75 and spatial frequency of 0.04 cycles per degree. Gray screen ‘blank’ trials, as well as static gratings in each orientation, were repeated 50 times, with randomized trial order. Each presentation, or trial, lasted for 1 s with 0 s inter-trial intervals. As such, the static grating block lasted 250 s (5 trial types X 50 repeats X 1s duration). In Step 2, we analyzed the 2p imaging data collected in Step 1 by running Suite2p on the computer where ScanImage was running. Before running Suite2p, we converted the ScanImage tif files into an h5 file as follows: 1) loading the ScanImage tif files; 2) cropping it into region corresponding to the holographic FOV; 3) keeping only the frames corresponding to the functional channel (e.g., every other frame if both red and green PMTs were recorded); and 4) saving as an h5 file. It was important to convert at least 5000 time points, because Suite2p identifies many more spurious ROIs with shorter recordings. Next, we ran Suite2p on the cropped h5 file. Because there is insufficient time to manually curate the ROIs during the online analysis, we analyzed tuning curves for every ROI returned by Suite2p.

In Step 3, we holographically stimulated neural ensembles defined in Step 2. Each target received 50mW, and each stimulation consisted of 10 pulses of 10 ms duration at 16 Hz. In addition to the co-tuned ensemble stimulation at 4 orientations, 0-power ‘blank’ trials and mixed-tuned ensemble stimulation trials were interleaved (i.e., 6 trial types). Each trial type was repeated 20 times with an inter-trial interval of 5 s.

### Functional connectivity experiments

In the functional connectivity experiments (Fig. 5), three datasets were acquired without visual stimulation (gray or black screen), and three with visual stimulation consisting of full-screen Gaussian-modulated noise patterns at 50% contrast. For these experiments, visual stimulation was triggered synchronously with optogenetic stimulation and was kept on the screen for the remaining trial duration. Data was collected across four independent sessions in two mice. In two of these sessions, photostimulation was performed under both visual and non-visual conditions in a randomized trial structure. For analysis, visual and non-visual trials were treated as separate sessions. For these experiments, we used virally injected transgenic CaMK2-tTA;tetO-GCaMP6s mice injected with pAAV-hSyn-GCaMP6m-p2A-ChRmine-Kv2.1-WPRE. Cells with elevated green baseline fluorescence were identified as putative opsin-expressing neurons and selected as photostimulation targets. Compact neural ensembles were selected within LM and V1, with each ensemble assigned a specific SLM phase mask and a set of discrete XY galvanometer positions. The corresponding phase mask and galvo coordinates were updated prior to each trial start. Imaging was performed at 3 Hz over a 3100 × 3100 µm² FOV encompassing eight contiguous visual areas (V1, LM, AL, RL, PM, AM, MMA, MMP), as well as surrounding regions. For functional connectivity experiments presented in supp. Fig. 19, we used virally injected transgenic mice (EMX1-Cre;CaMK2-tTA;tetO-GCaMP6s injected with AAV9-CAG.DIO.ChroME2s-FLAG-ST.P2A.H2B.mRuby3.WPRE.SV40). The holographic FOV was placed over LM. The imaging FOV encompassed V1, LM, RL, PM and AM (2400 X 2400 µm^2^). Stimulation consisted of long 250ms pulses, at zero power, P=30mW and P=50mW, with 200 trials per power condition to maximize detectability of long-range responses. P=30mW and P=50mW trials were averaged as the stimulation condition.

### Near simultaneous photostimulation of remote ensembles

In near-simultaneous remote photostimulation experiments (Fig. 6), putative opsin-expressing cells were segmented across the full 3.1 mm × 3.1 mm mesoscopic FOV, as in the inter-areal functional connectivity experiments. Each cell was assigned a specific XY galvanometer position, and neural ensembles were then defined and paired with a corresponding SLM phase mask and XY galvo coordinate. For near-simultaneous stimulation, two spatially remote ensembles were targeted by synchronizing SLM phase mask updates with XY galvo movements. The minimal galvo travel time between positions was calculated based on the specified galvo speed (1.2 ms per 0.4° optical deflection for QS30XY-AG scanners). An additional 5 ms buffer was added to define the final time interval between phase mask switches. Photostimulation consisted of 10×10ms pulses at 10Hz with 50 trials per condition.

### Offline analysis of mesoscopic two-photon calcium data

Motion correction and source extraction were done using Suite2p (Pachitariu, 2016), run through GPU for faster processing. Calcium sources were manually curated based on morphologically identifiable neurons. In Figs 2-3 experiments, frames with stimulation laser artefacts were discarded for downstream analysis and neuropil fluorescence was subtracted from the fluorescence traces using a 0.7 neuropil coefficient. Each cell’s fluorescence trace was then z-scored (Figure 2) or the dF/F was computed prior to z-scoring (z-scored dF/F). dF/F was defined as (F-Fmed)/Fmed where F is the neuropil corrected fluorescence trace and Fmed the median of the neuropil corrected fluorescence trace across all trials. A baseline of 2-5 frames before the stimulus onset was then subtracted for every trial (Figures 3,4). Photostimulation targets coordinates were remapped into the stitched mesoscale FOV and assigned to the closest suite2P identified calcium source. In Fig 4-6 experiments, there was no stimulation laser artefact except at the scanning stripe edges therefore all frames were kept for analysis. Fluorescence traces were z-scored across the entire experiment and 2 frames before the stimulus onset were subtracted for every trial. We found that neuropil subtraction had minimal impact on the causal influence maps (supp Fig. 23). For functional connectivity mapping experiments (Figs5,6), no neuropil correction was used.

### Definition of Modulated and Follower Cells

To quantify the impact of holographic perturbation across cortical areas, we classified cells based on their trial-by-trial modulation following photostimulation. Modulated cells were defined as cells located >100 µm from the nearest photostimulation target that exhibited a significant change in post-stimulus activity compared to no-power (0 mW) control trials (two-sided Mann–Whitney U test, *p* < 0.05). This category includes both strongly and weakly affected cells and captures the broader network influence of the stimulation. Follower cells were defined as a subset of modulated cells exhibiting high response consistency across trials. To identify them, photostimulation trials were randomly split into two non-overlapping subsets. Modulated cells were independently identified within each split using the same significance criterion (*p* < 0.05). Follower cells were defined as the intersection of the significantly modulated populations across both trial splits. This approach selects cells with consistent stimulus-evoked modulation and minimizes the inclusion of false positives driven by trial-to-trial variability.

### Functional connectivity metrics

The modulation strength metric computed in Fig. 5d quantifies the average amplitude of photostimulation-induced responses among follower neurons. It is computed as the sum of the absolute modulation amplitudes divided by the number of follower cells Nmod. Modulation Strength = 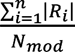. This provides a polarity-independent measure of the average effect size per modulated neuron, allowing comparison of response strength across conditions or cortical areas regardless of the number of recruited cells. The net influence metric computed in Fig. 5 quantifies the net directional impact of photostimulation on a downstream area. It is calculated as the sum of the signed modulation amplitudes across all follower cells, divided by the total number of follower cells. Net influence = 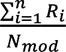. Positive values indicate a net excitatory effect, while negative values indicate a net inhibitory influence. Additional functional connectivity metrics are defined in Supp. Fig 22. These metrics were also calculated using a broader pool of putative connections (supp. Figs. 20,22).

### Statistics

All statistical analyses were performed using Python or Matlab. The analyses performed were paired t-test, Wilcoxon sum-rank and Mann-Whitney U test and tests were two-sided unless stated otherwise. No statistical method was used to predetermine sample size. Trial randomization was used in all applicable experiments (Figures 2,3,4,5,6).

**Supplementary note 1: Hologram axial resolution**

The 2P fluorescence signal of a temporally focused gaussian spot at the focus of the objective can be written as:

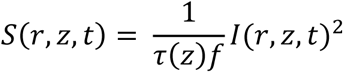

With τ the pulse width, f the repetition rate, I the intensity profile at the focus, r the radial dimension 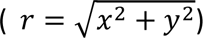 and z the axial dimension, both defined with the center of the gaussian spot as origin. Under the approximation that the linear chirp at the objective back aperture (αΩ, with α a constant proportional to the grating groove density and Ω the FWHM of the pulse frequency spectrum) is significantly smaller than the size of each monochromatic beamlet s 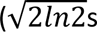 the FWHM of each monochromatic beamlet), the pulse width as a function of the axial depth z can be written as (cf. Papagiakoumou et al, 2020):

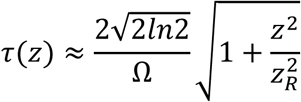

With 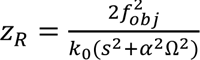 , f_obj_ the objective focal length, and k_0_ the wavevector of the central frequency of the pulse.

The 2P fluorescence signal can then be written as:

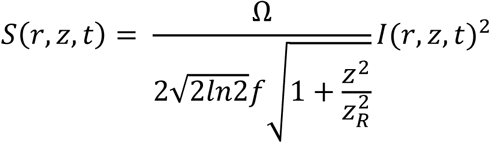

When injecting into this model numerical values corresponding to a conventional 2P objective (here Olympus 20X 1.0 NA f_obj_ = 9mm and 18mm back aperture size) and values corresponding to the 2P-RAM mesoscope objective (0.6 NA f_obj_ = 21mm, 25mm back aperture size), we were able to obtain, for a 400 fs pulse width at 1 MHz and at 1.03 μm wavelength, a cut-off threshold at the tail of the distributions corresponding to empirically observed axial hologram sizes: ∼ 30𝜇𝑚 for the 2P-RAM mesoscope and ∼ 20𝜇𝑚 for the conventional 2P microscope (cf. Pégard 2017, Mardinly 2018). The large back aperture of the mesoscope objective seems to partially compensate for the lower NA regarding axial temporal focusing resolution.

**Supplementary note 2: Physiological point-spread function**

The physiological point-spread-function (PPSF) is the main metric for quantifying single-cell resolution in two-photon optogenetics experiments. It is usually reported as the FWHM (full-width half maximum) of the neuron’s response as the optical hologram is digitally displaced along the axial axis. In this manuscript, we use the GCaMP calcium response as a readout for the neuron’s response. The effective PPSF can be partially modeled as the convolution of the optical axial point-spread-function with the cell diameter and is affected by several parameters (stimulation power, opsin expression, potency and localization and cell intrinsic properties, cf. Lees et al. 2024). We have performed extensive PPSF characterizations in mice expressing the very potent opsin ChRmine (Fig.2, supp Figs. 8,9 and 15) to provide a range of PPSFs to be expected by experimenters using our platform. At photostimulation powers well below saturation, we were able to achieve *in vivo* near single-cell precision with 33-35µm axial FWHMs (supp. Fig. 8 and 15) in both central and off-axis holographic FOVs. At saturating powers, we measured ∼70µm axial FWHMs in the central holographic FOV and ∼80µm off-axis. These values fall on the higher end but are still within the range of previously reported PPSFs in the field (Daie et al., 2021, Fisek et al., 2023). While these high-power conditions were the ones used in the functional connectivity experiments in the present manuscript, it is important to note that for many cells these conditions were on the saturation part of the power curve and the same experiments can be done at lower powers below saturation. Depending on the biological question under investigation, experimenters can choose their photostimulation parameters to optimize the trade-off between larger PPSFs and the number of elicited spikes per neuron for a set stimulation duration. To optimize this trade-off, one can adjust the individual power delivered to each cell (supp Fig 9, see also Bounds et al., 2023), at the expense of increasing experimental complexity. Notably, earlier PPSF measurements in Ai203×Vglut-Cre transgenic mice revealed an axial PPSF of 49 µm, well within the range reported for ChRmine-based viral expression. Lateral PPSF measurements ranged from 24 µm to 38 µm across powers and opsins, with the upper end corresponding to saturating photostimulation powers with virally injected ChRmine. (Fig. 2, supp. Fig. 8).

**Supplementary note 3: Assembly and alignment guide**

Detailed documentation on mesoscope assembly and alignment is available on the Janelia open wiki https://wiki.janelia.org/wiki/display/mesoscopy/Documentation. We focus below on the specifics for adding a 2P holographic module to the Janelia/Thorlabs mesoscope.

Prerequisites:

Adding a 2P holographic photostimulation path onto the mesoscope required some custom parts and modifications to the original mesoscope parts. Items 1-3 below were designed in collaboration with Thorlabs who manufactured the final parts. All the part designs are available on the CAD file attached. We do however encourage scientists willing to upgrade their system to reach to Thorlabs for assistance with the items 1-3 upgrades.

1. The third scan relay (scan lens and tube lens assembly) needs to be opened to merge the imaging and photostimulation beams. See CAD file for aperture size and position.

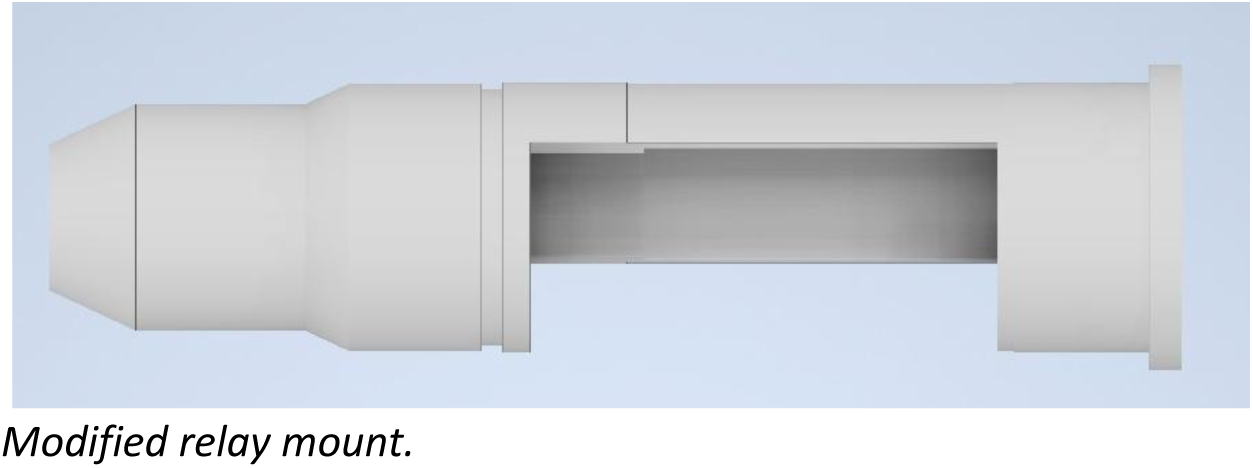

1. The 2P holographic path is set up on a horizontal custom 33”” × 24”” × 0.5”” aluminum breadboard (Thorlabs) attached to the main motorized mesoscope frame (gantry). The breadboard is attached to the gantry through two custom support clamps in the back (Thorlabs) and one custom L-bracket on the front (Thorlabs). Securing the L-shape bracket in place required drilling 4 ¼”-20 tapped holes on the mesoscope rotation hub.

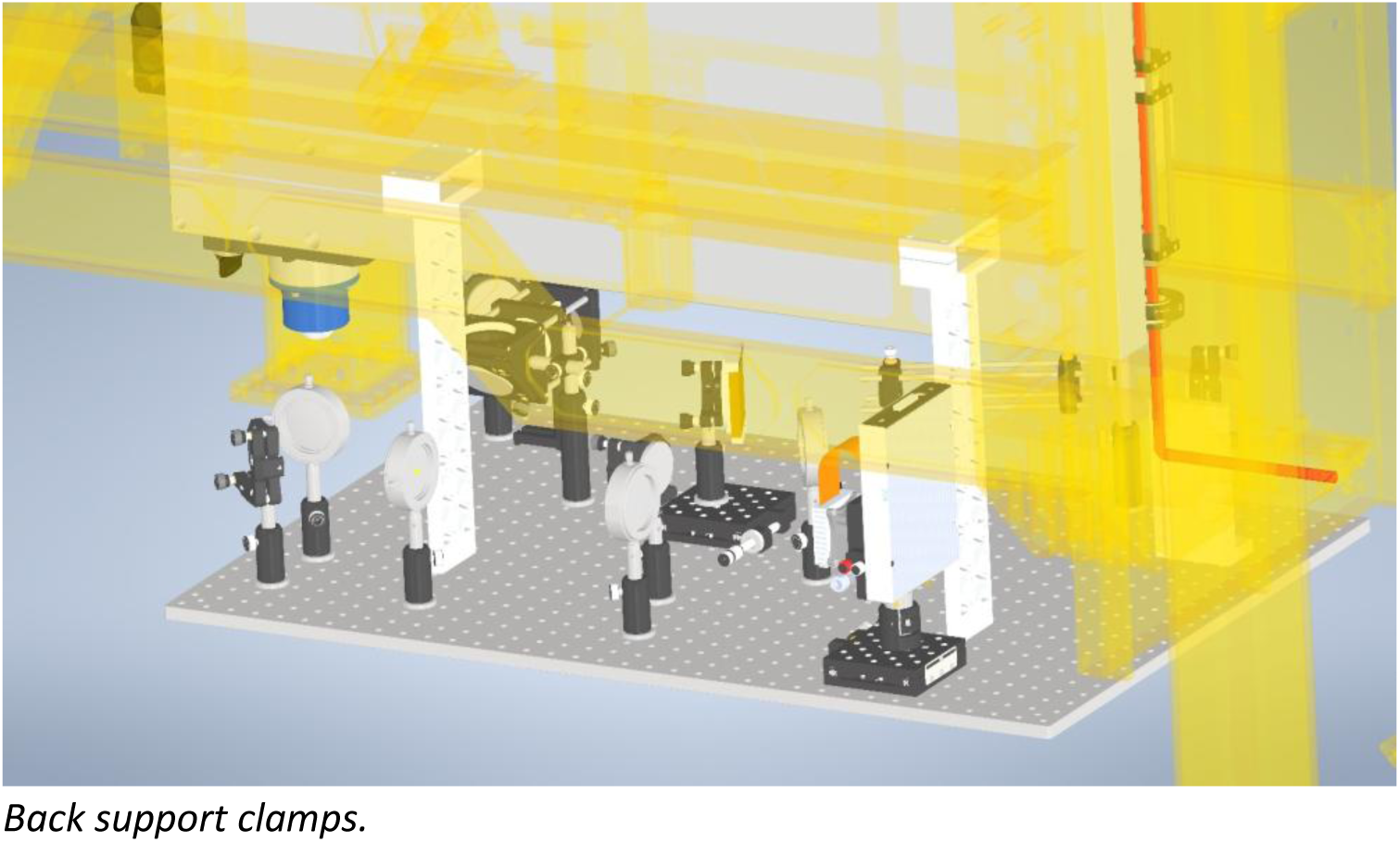

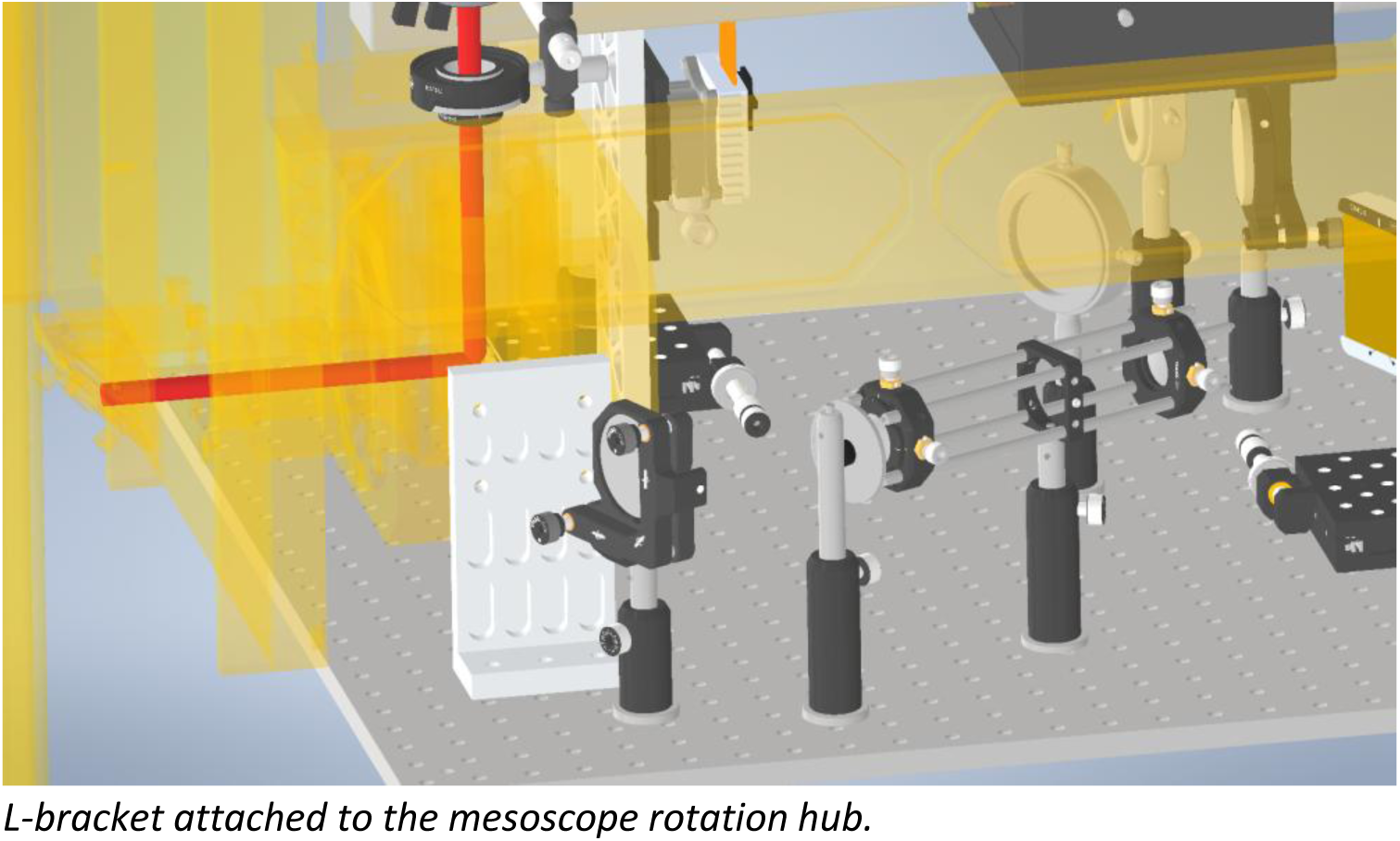

1. A custom threaded bracket (Thorlabs) was used to attach the recombination dichroic D3 onto the vertical mesoscope breadboard.

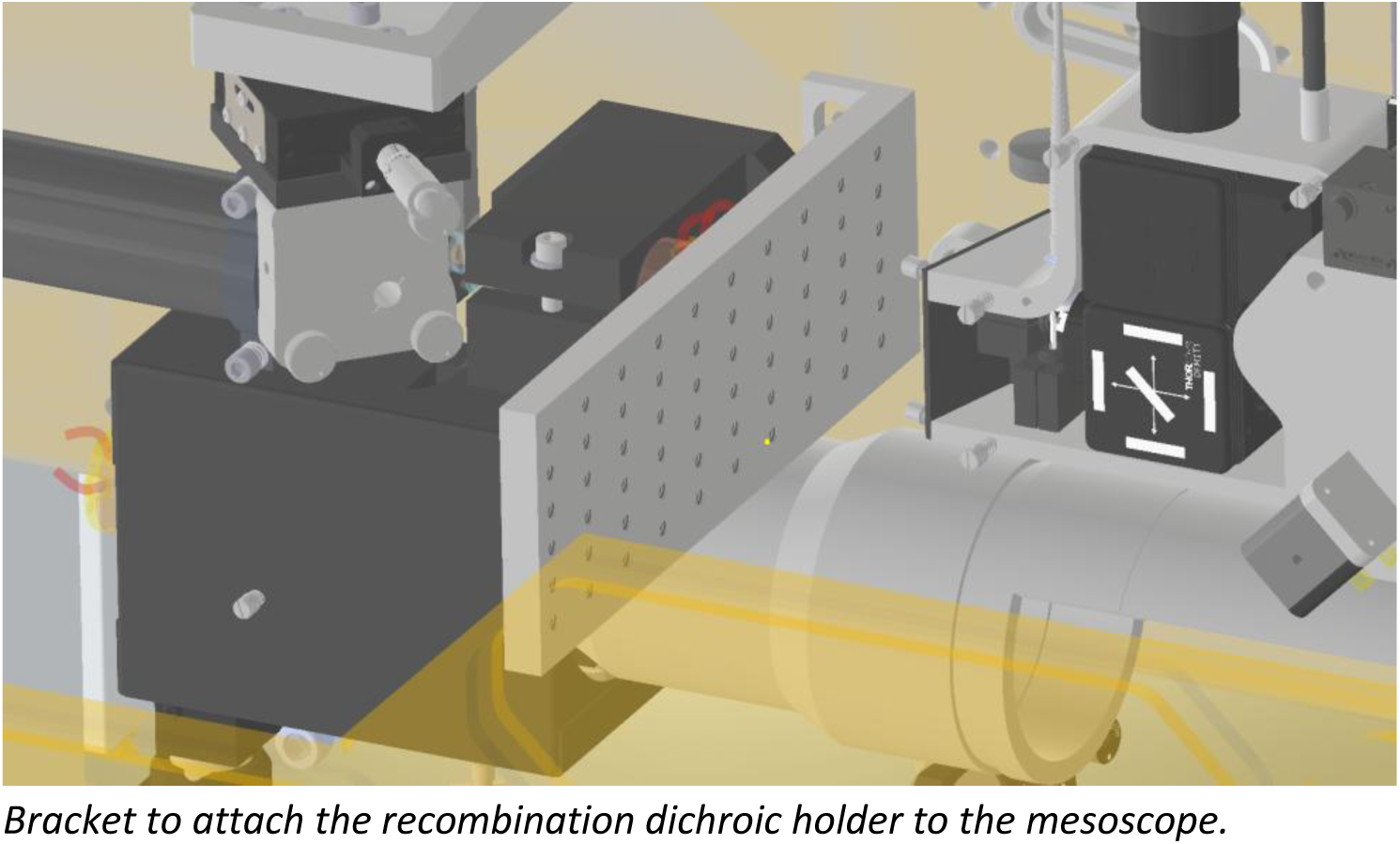

1. A custom 3D-printed holder is used to flexibly position the recombination dichroic D3 on the mesoscope scan path. The holder allows flexible positioning across 3 translation directions and one rotation axis and was designed to hold a standard ½” Thorlabs post. The dichroic is secured in place with a FFM1 holder (Thorlabs) secured on a B3C mount (Thorlabs).

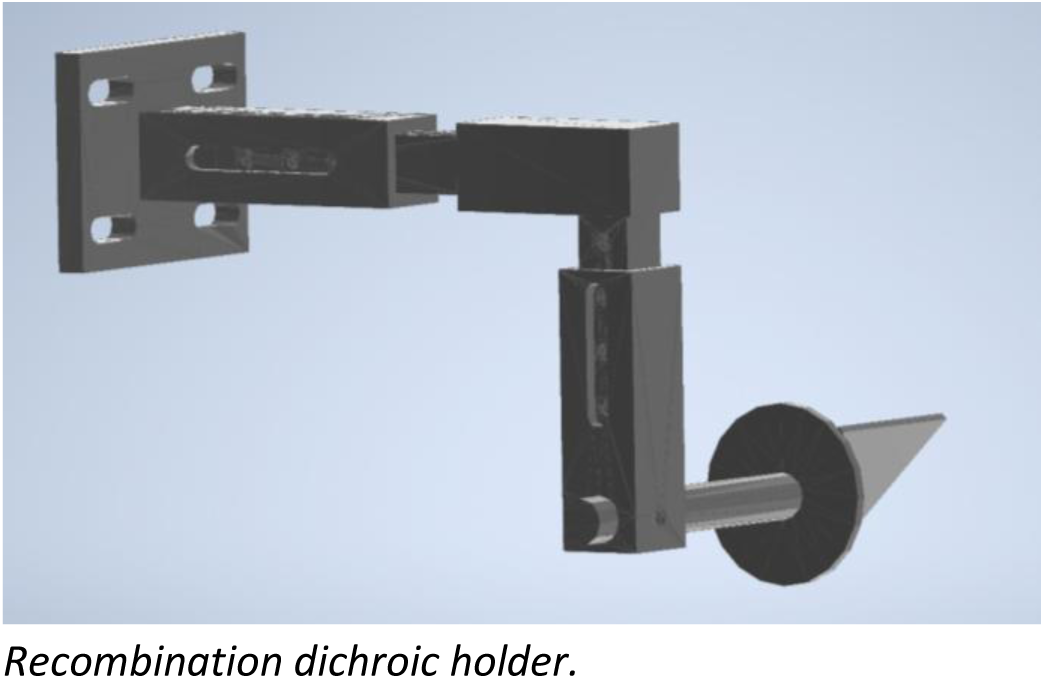

1. To maintain axial alignment, the motorized axial movement of the vertical breadboard relative to the gantry is locked and replaced with an external motorized lab jack (Thorlabs, MLJ150).
2. Assembly and alignment:
3. Once the breadboard is installed, pre-position all breadboard optics to match the 4f requirements using the CAD layout as a guide.
4. Align the imaging beam up until the rotation hub to be centered on each of the gantry periscope mirrors (M1,M2,M3,M4,M5) and perfectly stationary with the gantry’s X and Y translation.
5. Remove the last periscope mirror (M6) and place the provided rotation alignment tool. If step 2 is done properly, M5 is enough to steer the beam to be centered and leveled along the rotation axis and to have the beam position invariant with rotation.
6. Inject the stimulation beam onto the periscope so that the beam is co-aligned to the imaging beam i.e centered on all mirrors and invariant with X, Y and rotation. For this step only the D1 dichroic and mirrors external to the gantry periscope can be used. Consider having the recombination dichroic D1 installed on a lateral translation stage to have an additional degree of freedom for co-alignment.
7. Given the natural divergence of the laser beam and the long distances on the mesoscope, the beam should be quite expanded on the rotation alignment tool. To correct for that, we added a 1:1 telescope before the gantry (lenses L01 & L02) to have a more collimated beam impinging on the grating. The lenses of the telescope need to be placed centered on the beam so that the beam remains undeflected. L02 could be placed on a translation stage to allow for easier adjustment of the divergence while maintaining the alignment. Fine adjustments of the imaging/stimulation beam co-alignment can be made after the telescope is installed (again, only with mirrors external to the gantry or with the D1 dichroic).
8. Remove the rotation alignment tool and replace M6 with the D2 dichroic. At this stage, the imaging and stimulation beams can now be aligned independently.
9. Subsequent alignment of the imaging path on the vertical breadboard is now constrained since only D2 can now be used to route the beam straight on the pre-RF vertical axis. Therefore, depending on how the system has been originally set, the whole pre-RF vertical axis might need to be translated horizontally so that the beam can be straight and vertical along that axis.
10. After steps 4 and 5, the stimulation beam should be perfectly leveled impinging on the grating. Adjust the angle of the grating to minimize power lost at higher diffraction orders. Then iterate adjusting the height and position of the relay rail and adjusting the tip/tilt on the grating mount so that ultimately the beam is perfectly centered on L1 and L2.
11. Install the spinning diffuser and adjust the subsequent mirror to stay leveled at the beam height and to hit the SLM injection mirror while being centered and on-axis on the L3 lens.
12. Load a flat phase on the SLM (or simply turn it off). Use the pre-SLM injection mirror and the tip-tilt knobs on the SLM to make the beam centered and leveled on the L4 axis. Having the SLM set on a translation stage can be helpful at this stage since it allows to correct for any clipping issue without changing the angle of the SLM. If the beam is properly centered on the SLM, the beam after L4 should look like a full rectangle beam on an IR card. With subsequent mirrors, the beam should then be aligned to be centered and leveled on the L5 and L6 axes. Prior to this, the cage rail system holding L6 need to be pre-positioned precisely under the imaging relay aperture so that a 90° beam exiting the cage system is centered on the imaging scan relay vertical axis.
13. Place the recombination dichroic D3 in order to have the stimulation beam centered on the mesoscope objective back aperture. Iterate between the position and angle of D3 and the tip/tilt knob of the last cage mirror to have the beam both centered on the back aperture of the objective and straightly vertical on the objective axis.

All lenses are oriented with the curved face towards the collimated beam to minimize cumulative spherical aberrations.

---

**Supplementary Figure 1.**
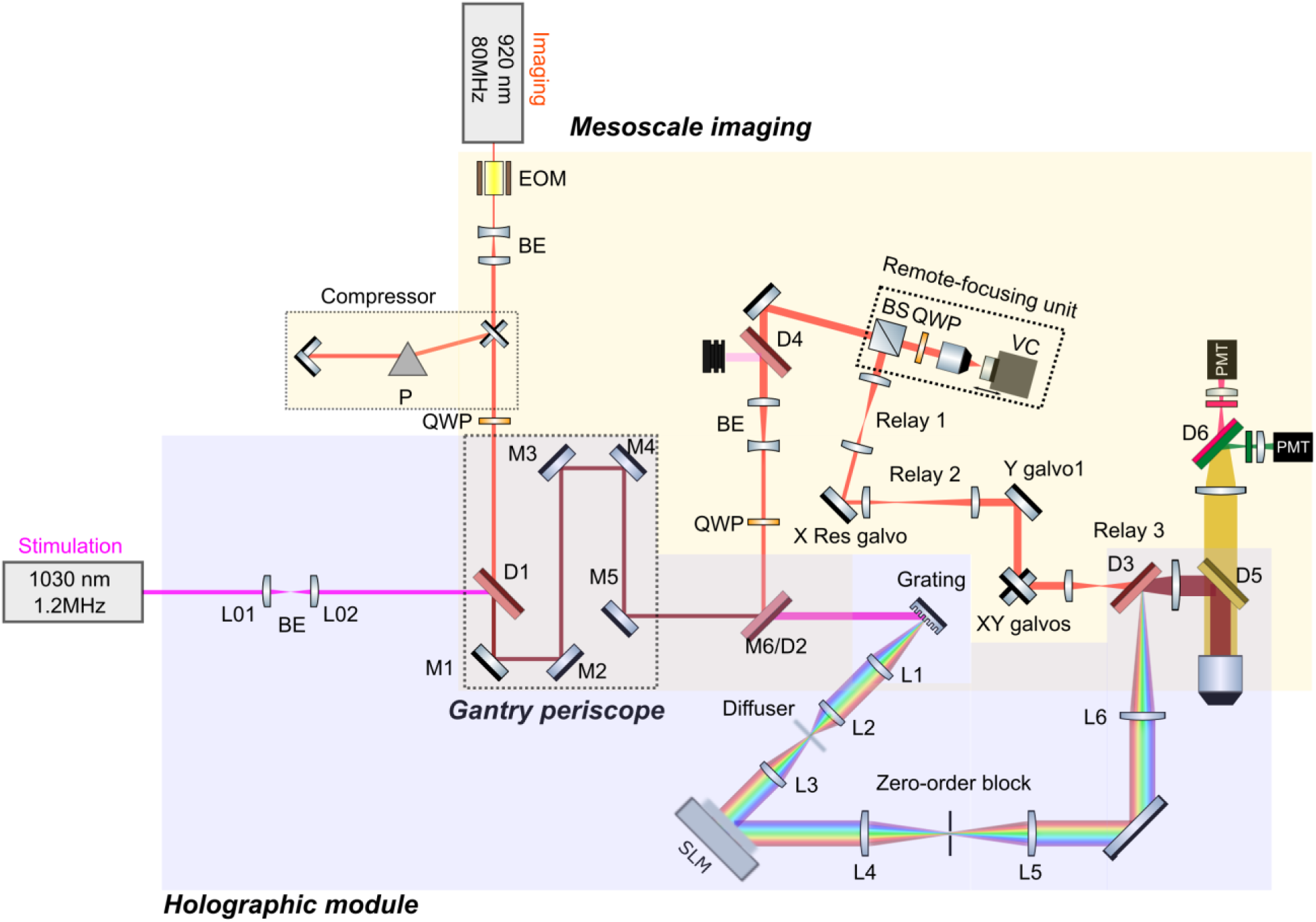
3D-SHOT two-photon holographic mesoscope optical setup. EOM: electro-optic modulator, BE: beam expander, P: prism, QWP: quarter waveplate, Dn: dichroic mirror, BS: polarization beamsplitter, VC: voice coil, SLM: spatial light modulator. M6 mirror in the original mesoscope design has been replaced with the D2 dichroic. Implementation in Figs.1,2,3.

**Supplementary Figure 2.**
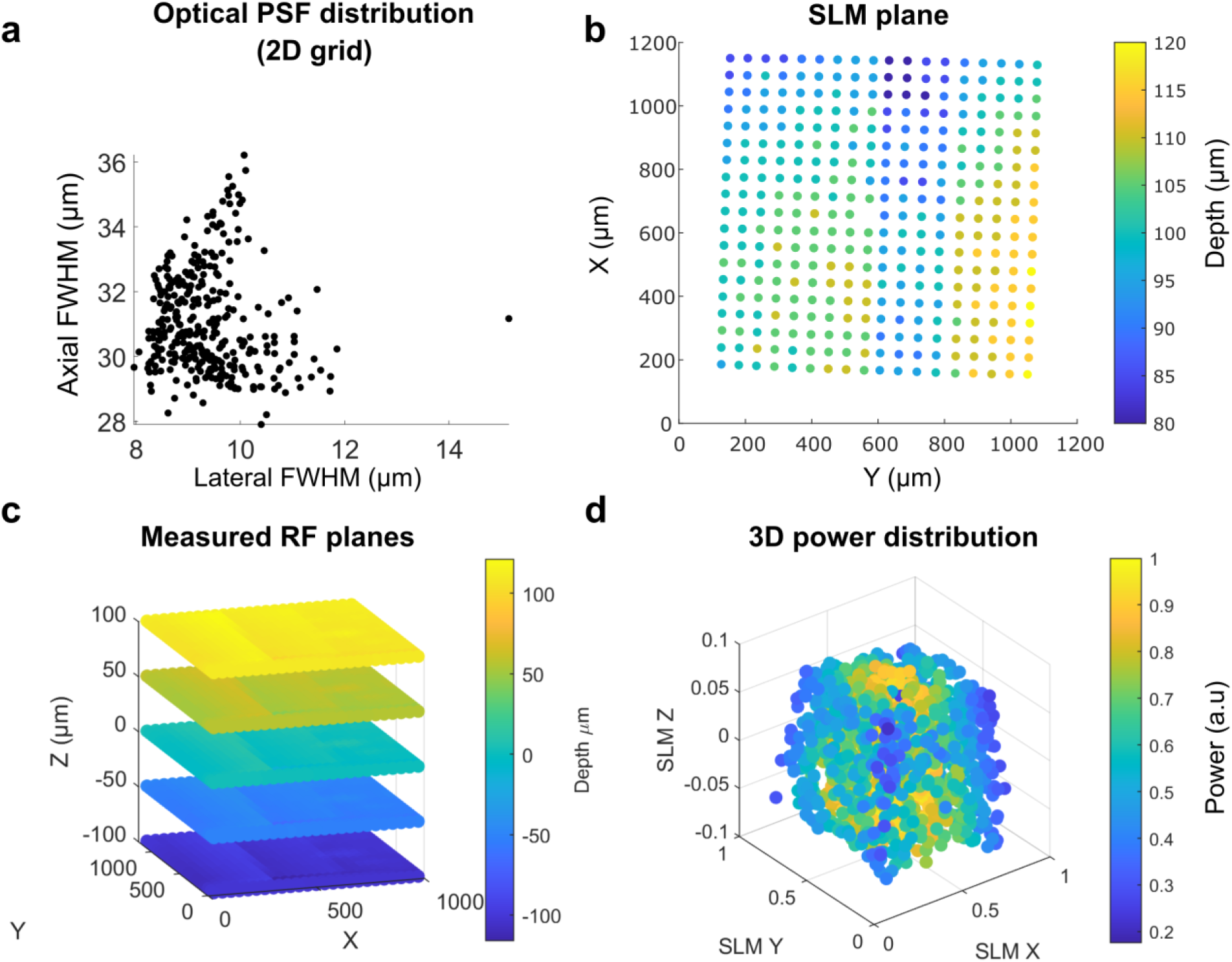
Optical characterization of the 2P holographic mesoscope. **a)** Distribution of optical point-spread-functions of holograms arranged in a 2D grid across the holographic FOV. **b)** Axial locations in real space (units: microns, arbitrary origin) of holograms with equi-Z SLM coordinates across the holographic FOV. **c)** 3D remote-focusing fluorescence imaging planes in real space measured with an inverted substage camera with voice-coil curvature correction on. The recorded holograms are registered to these recorded planes for accurate 3D calibration. **d)** Power distribution variation in 3D measured as the normalized square-root of the 2P fluorescence signal for a distribution of holograms randomly positioned in 3D space. This distribution is used to build a 3D model of diffraction efficiency that is used to dynamically correct the power allocated to each hologram as a function of its location in space, thus allowing to use a greater range of the SLM.

**Supplementary Figure 3.**
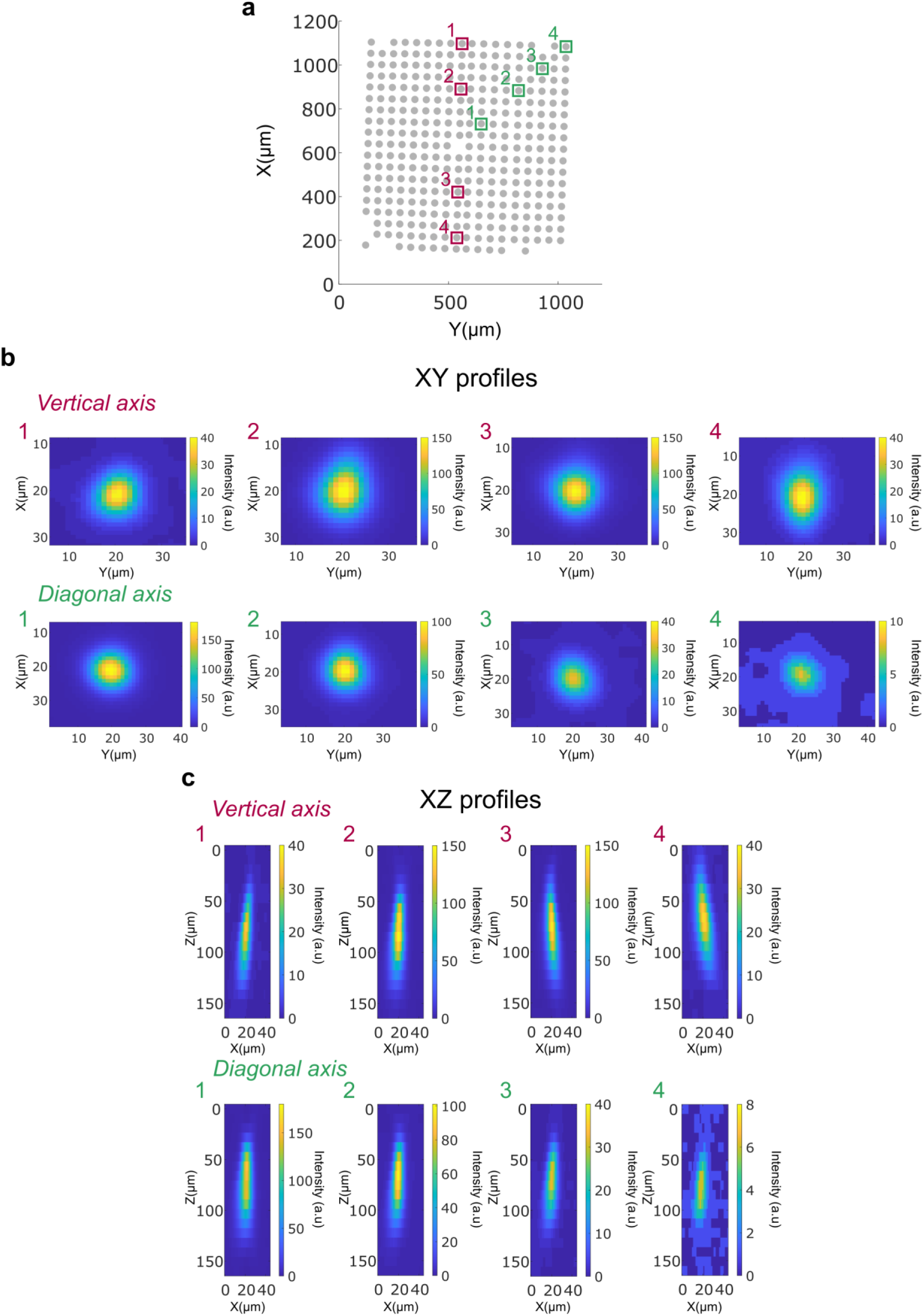
Example hologram intensity profiles across the SLM FOV. **a)** Detected 2D locations of holograms used for SLM FOV characterization. Cherry and green boxes indicate the locations of the select examples of off-axis holograms shown below. **b)** Lateral 2P fluorescence intensity profiles at select off-axis SLM FOV locations. **c)** Axial 2P fluorescence intensity profiles at select off-axis SLM FOV locations.

**Supplementary Figure 4.**
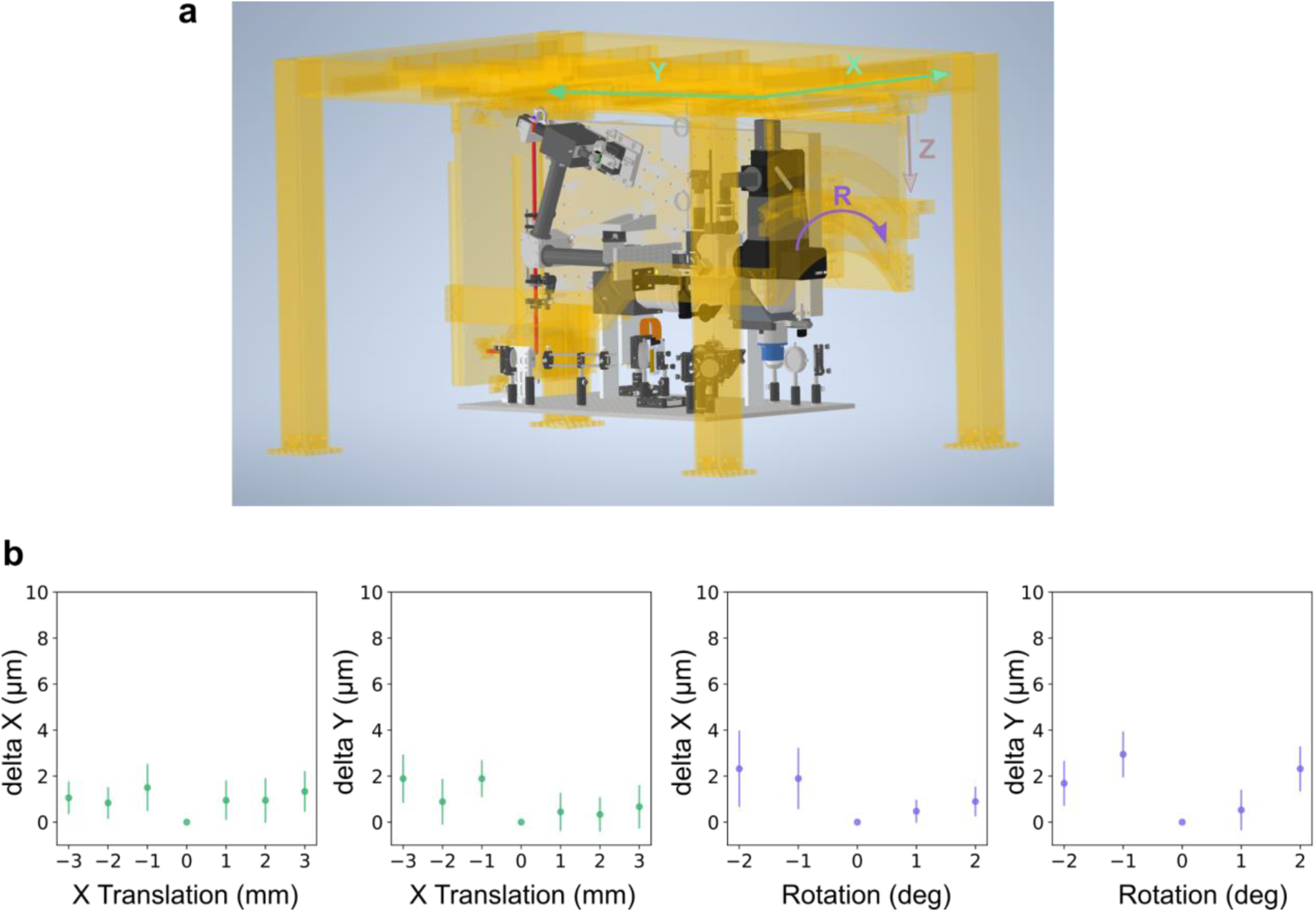
Targeting invariance with 2P-RAM mesoscope movement. **a)** The microscope main frame is motorized in X, Y translation and rotation. The holographic module was designed to be solidary to the main frame thus preserving motion along these axes. Axial Z translation in the 2P-RAM mesoscope is implemented by moving the vertical breadboard relative to the main frame. To maintain holographic invariance, that axis was locked and replaced by a motorized lab jack (MLJ150, Thorlabs) to translate the animal axially relative to the microscope. **b)** Targeting invariance measured for translation and rotation of the microscope around a central reference position. Delta X and Y are computed, respectively, as the difference between the X and Y location of the centroids of holographically burnt holes in a fluorescent slide at the translated or rotated position and the X and Y location of the centroids in the central reference position. Markers: mean value, error bars: standard deviation.

**Supplementary Figure 5.**
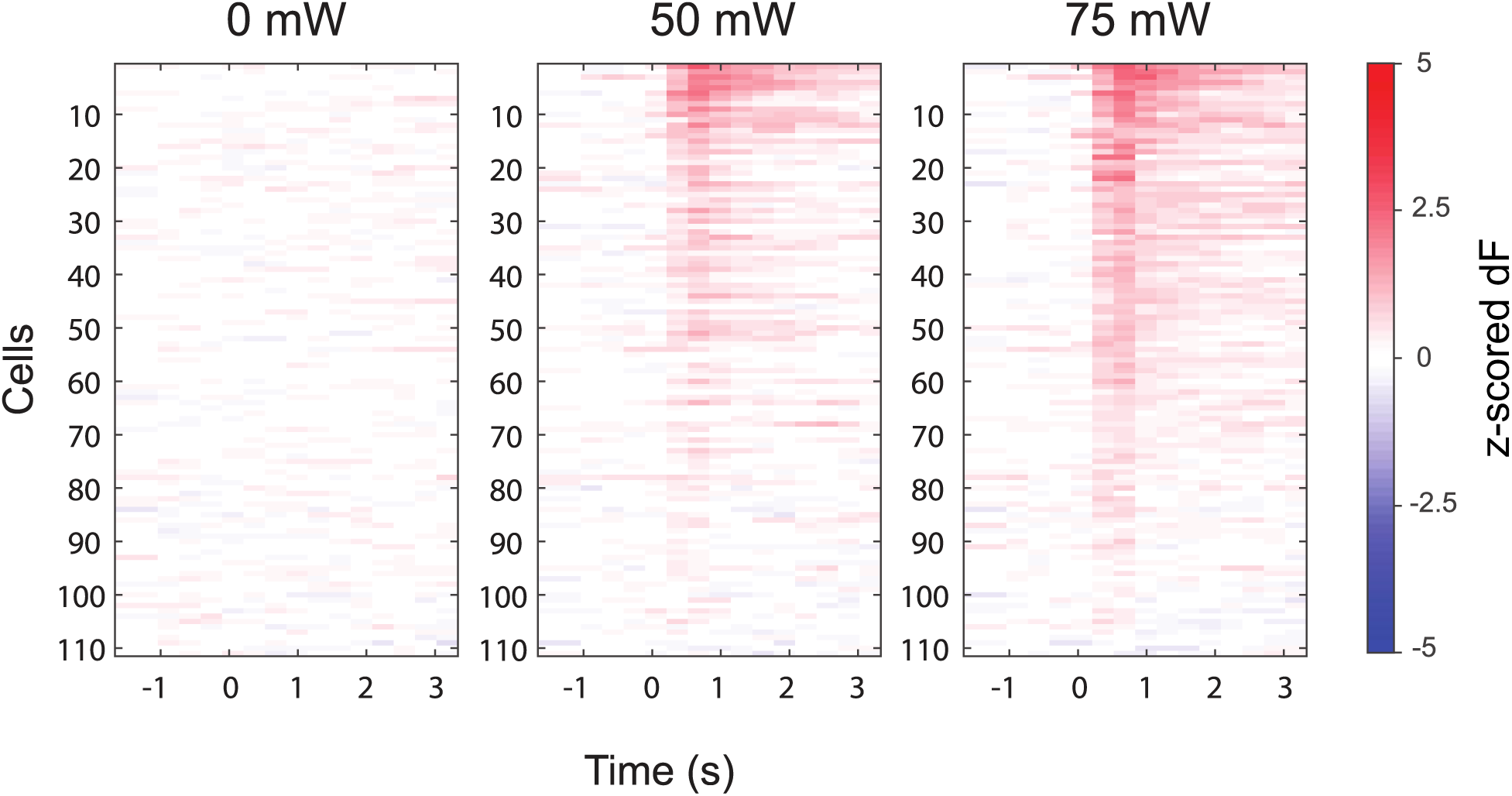
Sequential stimulation at different laser powers. Heatmaps of stimulus-aligned calcium responses (z-scored dF) for 111 targeted cells in an example session (Ai203 × VGlut Cre transgenic). 13, 58 and 75 were significantly activated at 0, 50, and 75mW, respectively, signed rank test, one tailed p<0.025).

**Supplementary Figure 6.**
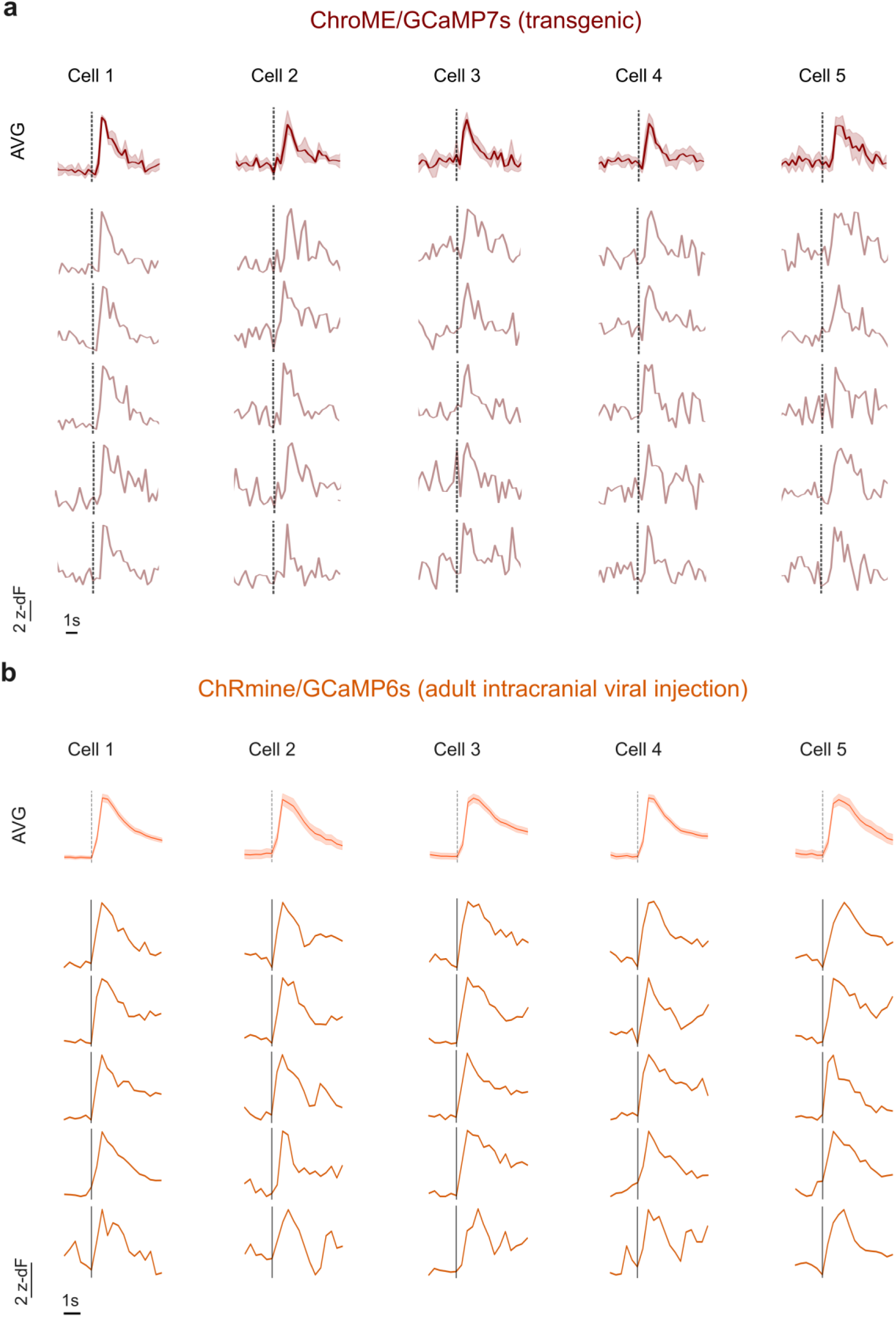
Reliable target photostimulation across trials. **a)** Example average (5trials) and single-trial traces across photostimulation repeats of target neurons co-expressing ChroME and GCaMP7s (Ai203xVglutCre transgenic). Photostimulation at 50mW. **b)** Example average (100 trials) and single-trial traces (5 randomly selected among 100) across photostimulation repeats of target neurons co-expressing ChRmine and GCaMP6s. Photostimulation at 10mW.

**Supplementary Figure 7.**
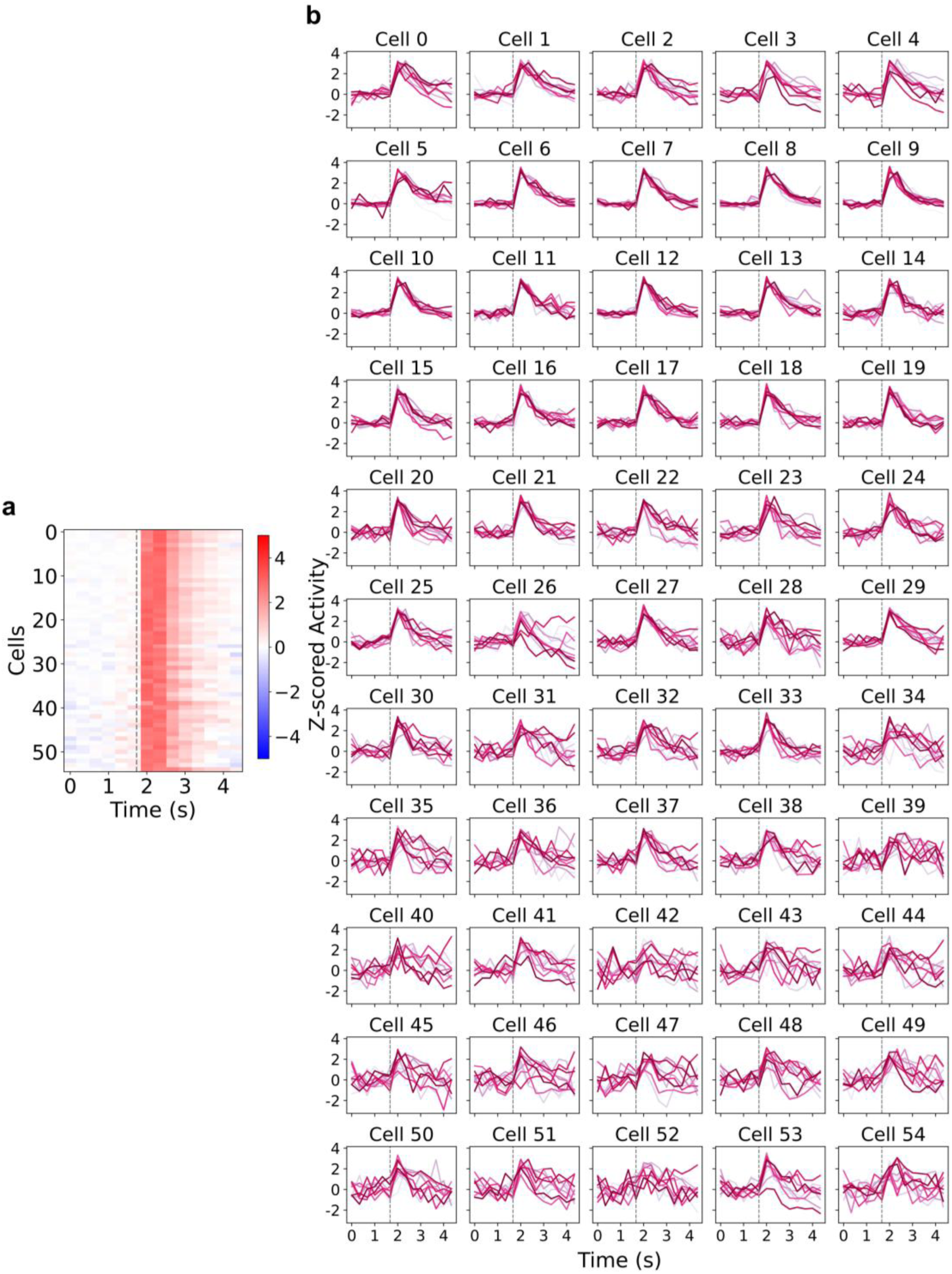
Reliable photostimulation of a large neural ensemble. **a)** Trial-average (n= 20 trials) post-stimulus time histogram (PSTH) of target neurons (n=55) simultaneously photostimulated. Dashed line indicates photostimulation start time. All neurons co-express GCaMP6s and ChRmine. **b)** Single-trial traces of individual target cells from the photostimulated neural ensemble.

**Supplementary Figure 8.**
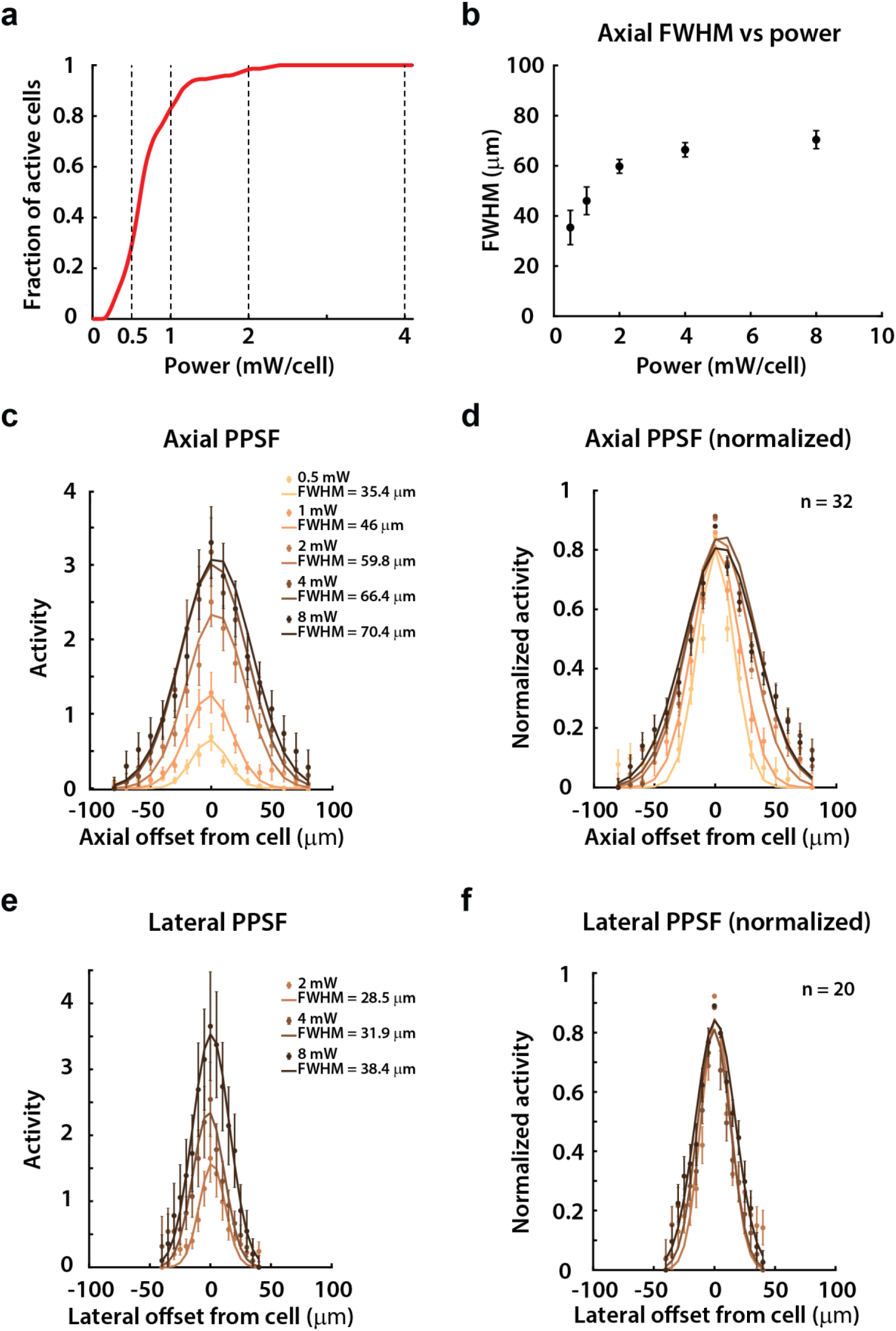
Physiological point-spread functions (PPSFs) as a function of power. **a)** Cumulative density histogram of the fraction of active cells used in the PPSF experiments (ordinate) that were active above a certain power (abscissa). **b)** Mean +/- s.e.m. axial FWHM across cells as a function of power. **c)** Axial PPSF profiles for each power condition. The filled circles and error bars are the actual mean +/- s.e.m. across cells, and the solid traces are weighted gaussian fits. **d)** Normalized axial PPSF profiles for each power condition, normalized to the centered hologram location (axial offset = 0). **e)** Similar to c), but for lateral PPSFs. **f)** Similar to d), but for normalized lateral PPSFs. All PPSF experiments were done in CaMK2-tTA;tetO-GCaMP6s mice injected with a pAAV-hSyn-GCaMP6m-p2A-ChRmine-Kv2.1-WPRE virus at discrete locations across visual cortex.

**Supplementary Figure 9.**
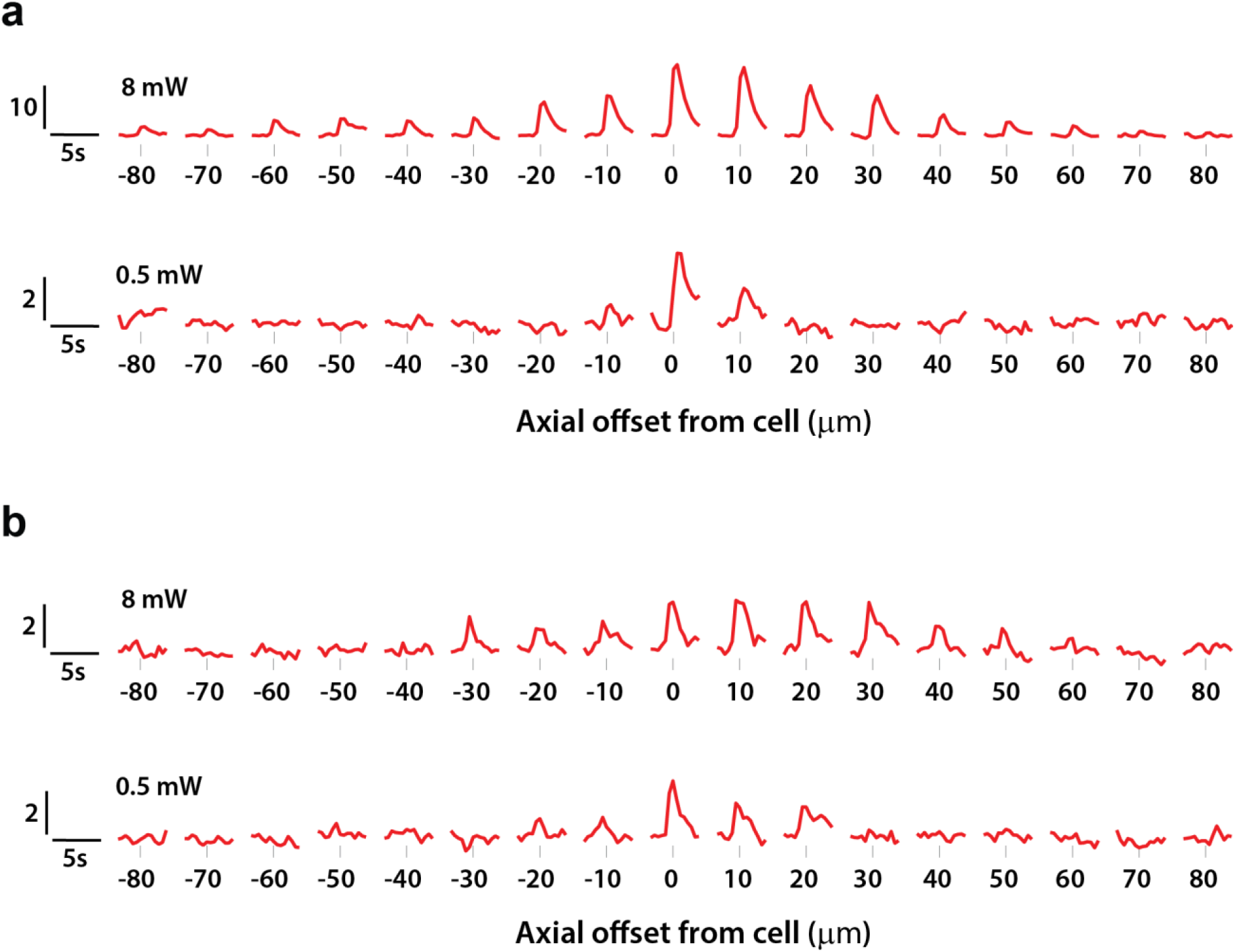
Example calcium traces from PPSF experiments. **a)** DF/F traces for an example cell at various axial offsets (columns), at high (first row) and low (second row) powers. **b)** Similar to a) for another example cell.

**Supplementary Figure 10.**
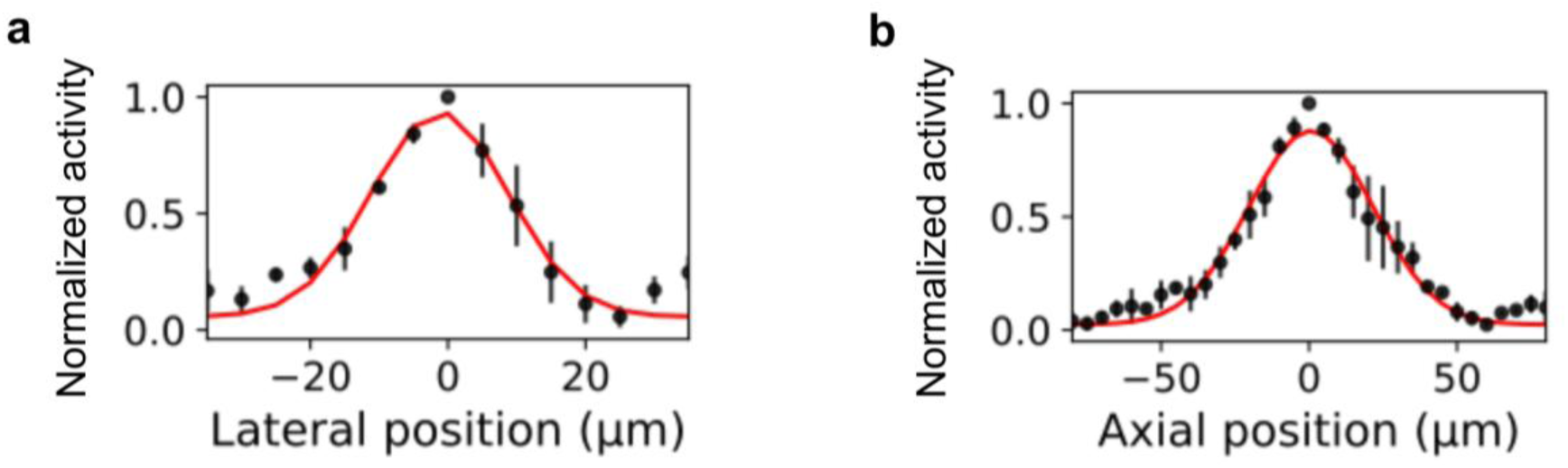
Physiological point-spread-function measured in Ai203;VGlut-Cre. *In vivo* optically measured physiological point-spread functions (n=3cells). Cells co-expressing GCaMP7s and ChroME. Error bars: S.E.M. Red curve: Gaussian fit, lateral full-width-half-maximum (FWHM) 23 µm, axial FWHM: 49 µm. Photostimulation with 5x5ms pulses at 30Hz at 25mW.

**Supplementary Figure 11.**
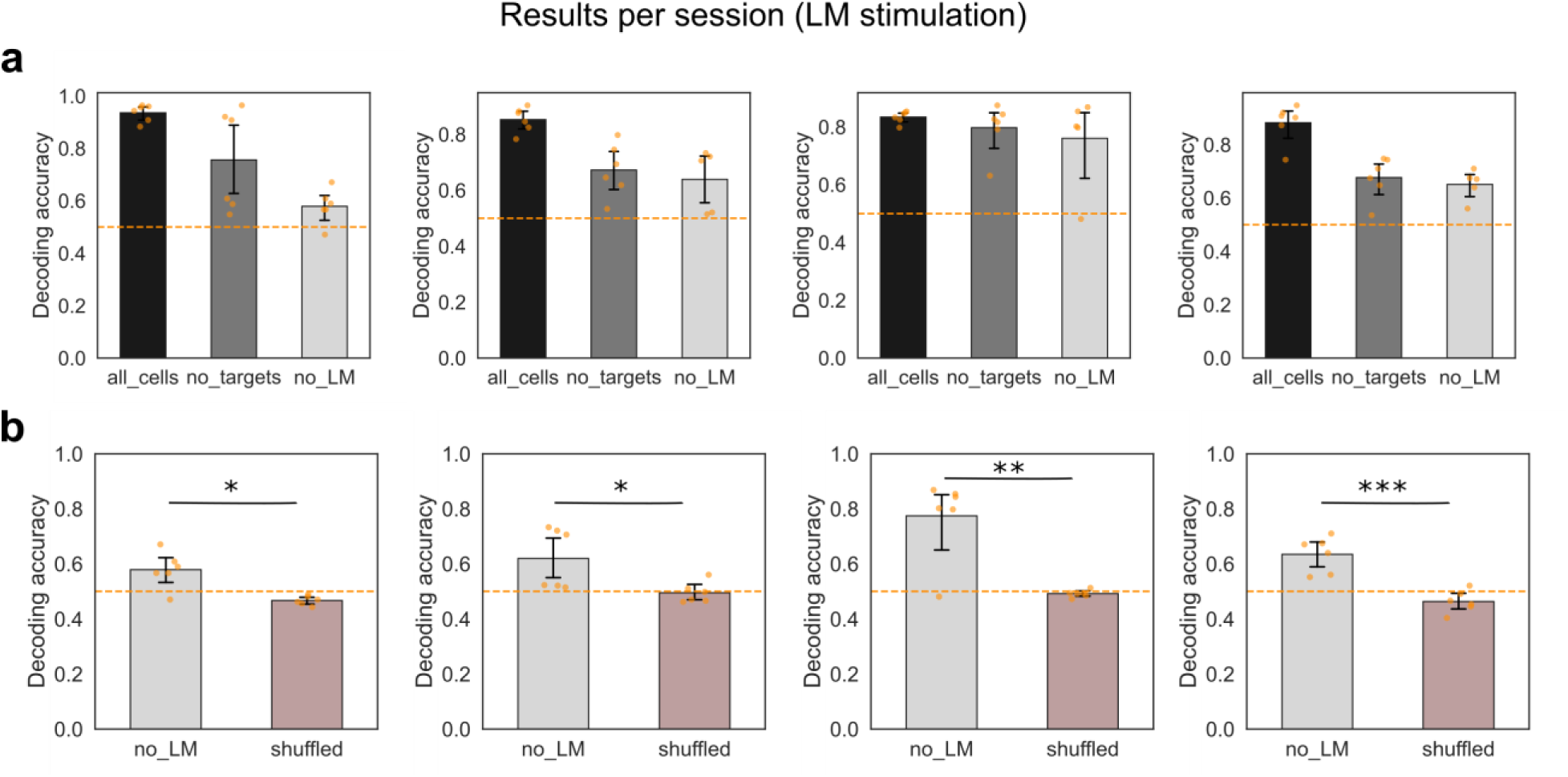
Photostimulus decoding results per session. **a)** Decoding accuracy of a classifier trained to discriminate the holographically induced patterns in LM from different groups of non-targeted cells in the FOV, split per session. 6 pairs of stimuli are presented for discrimination in each session. All cells: decoding from all neurons in the mesoscale FOV. No targets: decoding from all neurons except the targeted cells. No LM: decoding from all neurons outside area LM. b) Decoding accuracy per session of a classifier trained to discriminate the holographically induced patterns in LM from all neurons outside area LM with correct and shuffled labels in the training set. Session1/Mouse1: p = 0.013, Session2/Mouse1: p = 0.023, Session1/Mouse2: p=0.005, Session2/Mouse2: p = 0.0006.

**Supplementary Figure 12.**
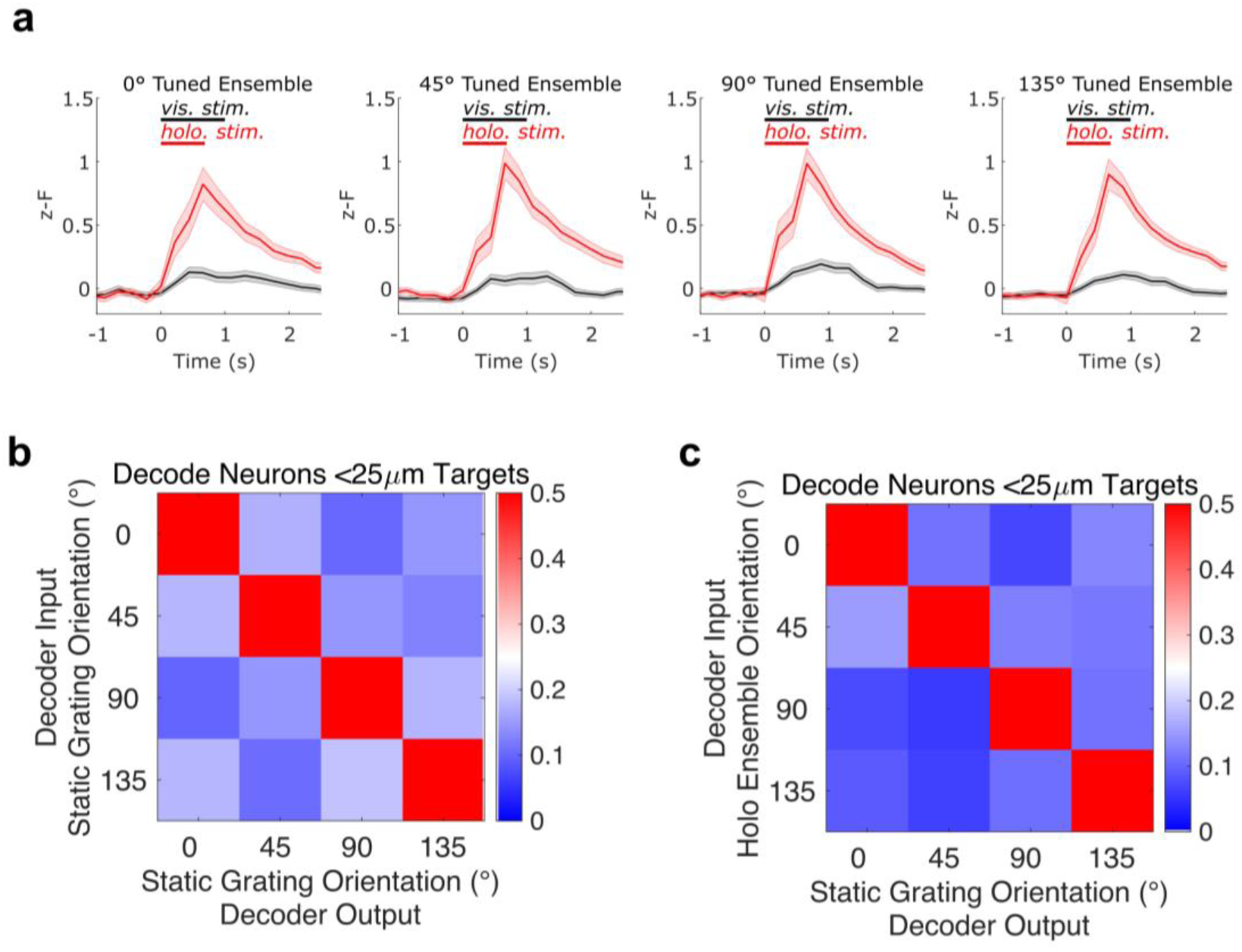
Stimulation of functionally specific neuronal ensembles sharing orientation preference in mouse visual cortex. a) Visual (black) and holographic (red) responses of holographically targeted neurons in each ensemble are shown (mean ± SEM across neurons, n=53/62/71/70 neurons in 0/45/90/135° ensembles respectively, aggregated across 11 sessions from 2 mice). Visual responses were driven by static gratings in each orientation, presented for 1s with no intervening period. Holographic responses were driven by synchronous photostimulation of the targets, where each target received 10 of 10ms pulses at 16Hz and 50mW power. b) Classification accuracy of a linear support vector machine (SVM) trained to classify visual response of 4 orientations, for neurons within 25 μm of each target in holographic ensembles (average 77.7 neurons per session). 10-fold cross-validated decoding accuracy was 0.557 ± 0.019 mean ± standard error mean (SEM) across N=11 sessions (p = 9.77 × 10^−4^ Wilcoxon signed-rank test compared to chance-performance of 0.25). c) Classification accuracy of the same SVM when probed with orientation-specific holographically evoked activity without any visual stimulus (0.696 ± 0.043 mean ± SEM, p = 9.77 × 10^−4^ Wilcoxon signed-rank test across N=11 sessions). Decoding accuracy was measured as the proportion of holography trials that were classified as the corresponding grating orientation.

**Supplementary Figure 13.**
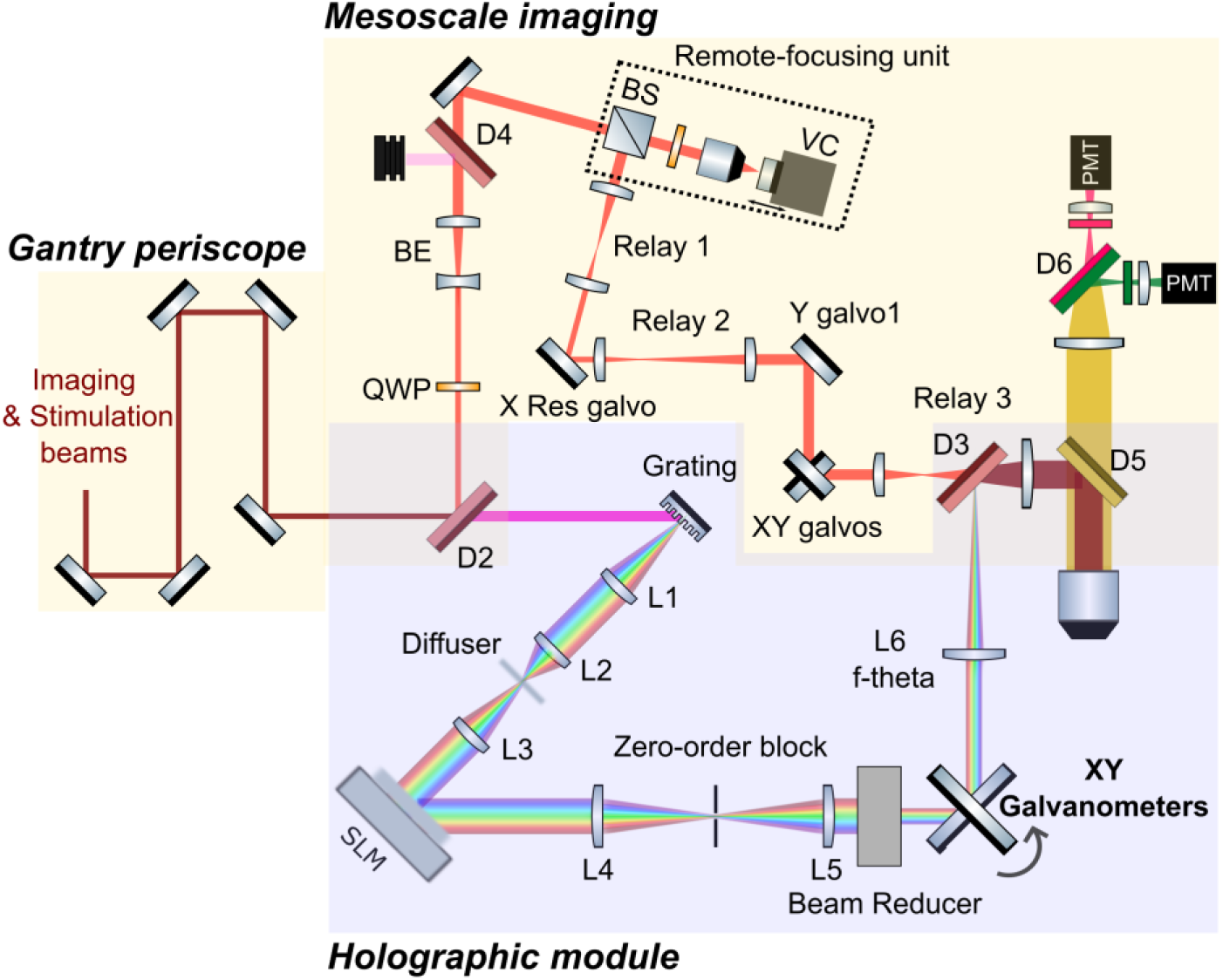
Random-access 3D-SHOT optical layout. A beam reducer and an XY galvo scanner pair are introduced in the collimated space between L5 and L6. L6 is replaced by an f-theta scan lens. The recombination dichroic D3 is replaced with a larger aperture mirror. Implementation in Figs.4,5,6.

**Supplementary Figure 14.**
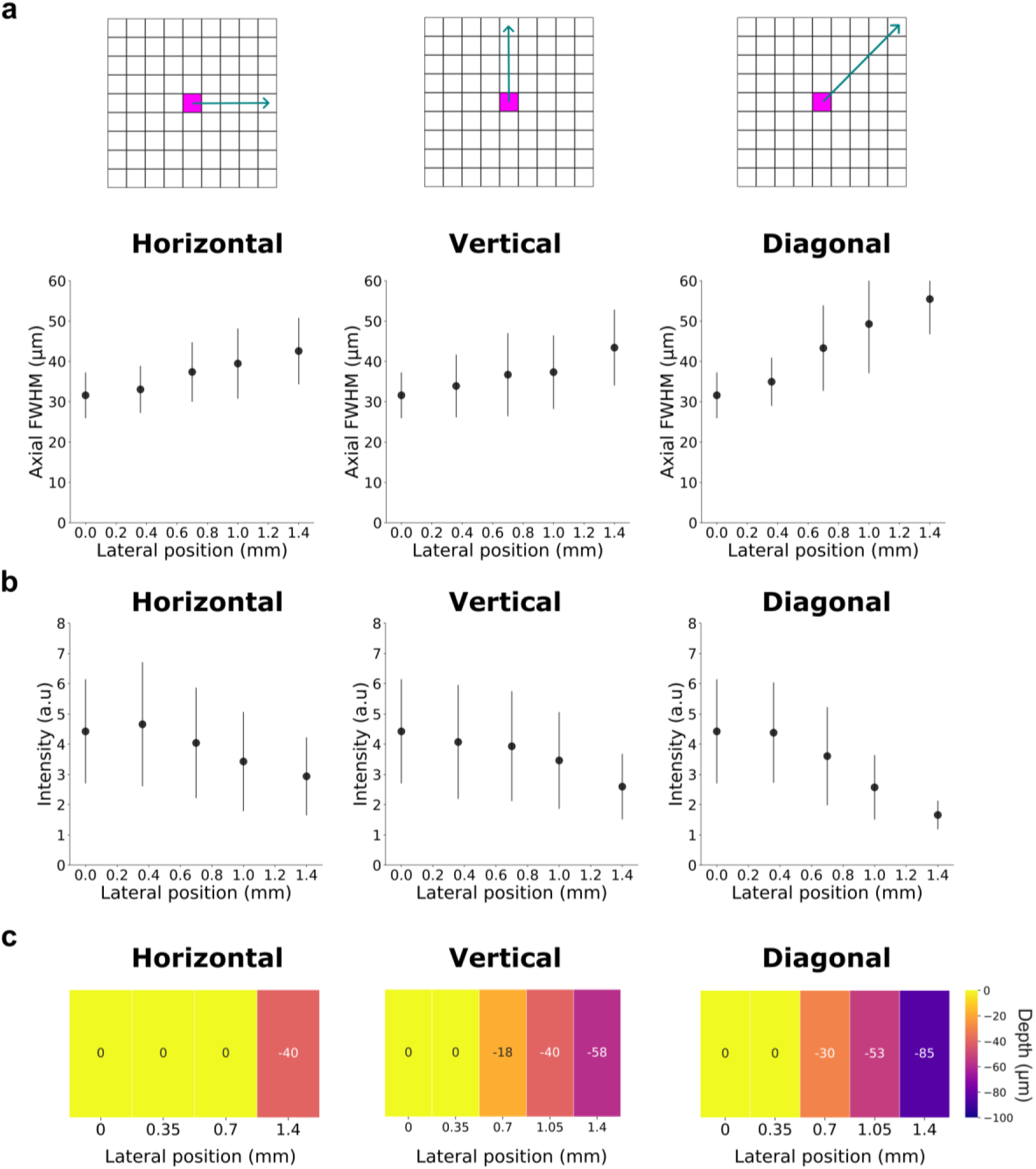
Optical characterization of random-access 3D-SHOT. a) Average axial resolution measured across 3D distribution of spots at different galvanometer scan positions across the horizontal, vertical and diagonal axes. Error bars indicate standard deviation values. N=70-380 holograms at each galvanometer position. b) Average single-hologram intensity measured across 3D distribution of spots at different galvanometer scan positions across the horizontal, vertical and diagonal axes. Error bars indicate standard deviation values. c) Axial offset between imaging plane and holographic plane z=0 plane (no defocus).

**Supplementary Figure 15.**
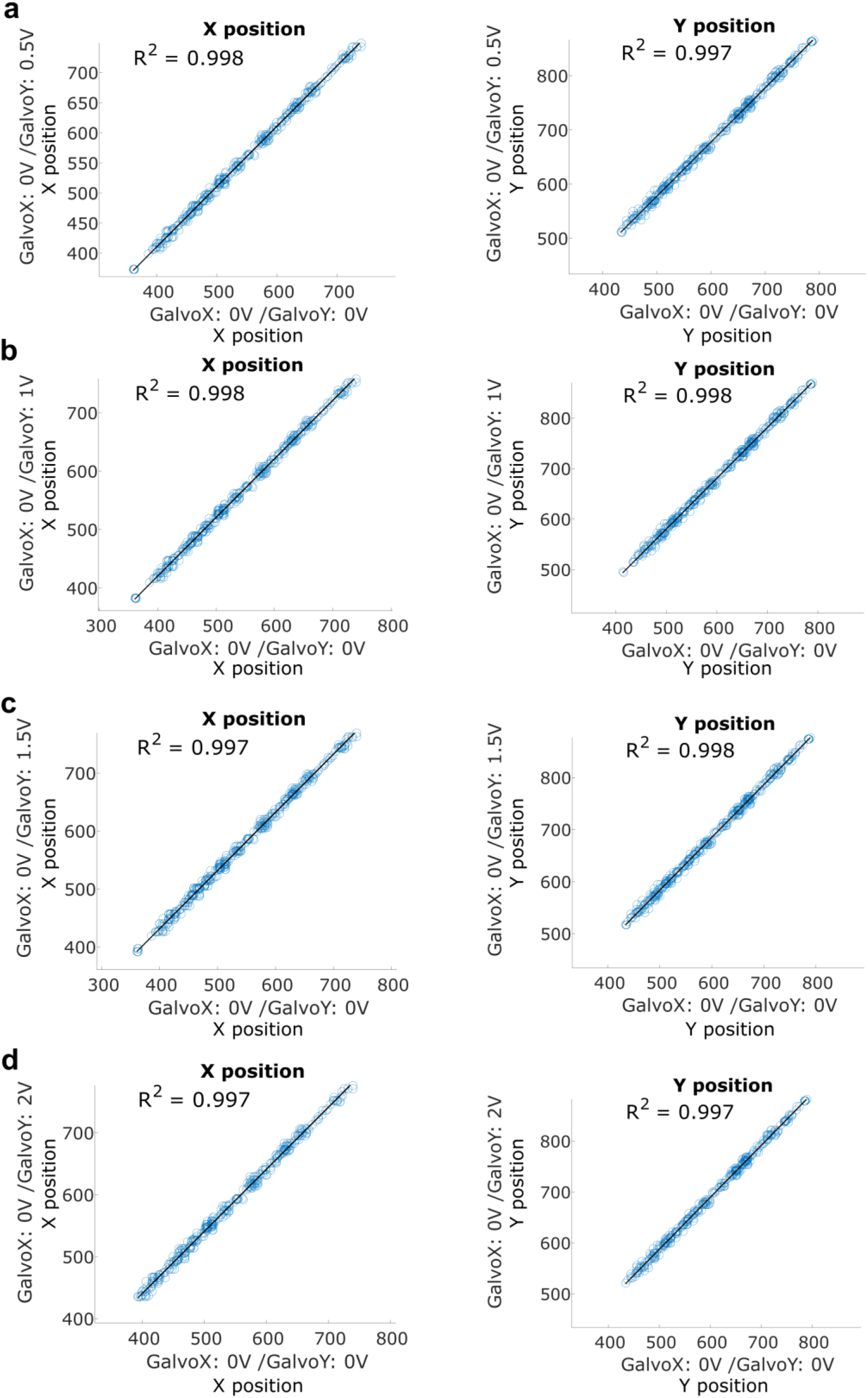
Calibration linearity across holographic field-of-views. Measured X and Y hologram coordinates when the galvo pair is at the central position (GalvoX:0V/ GalvoY:0V) plotted against X and Y coordinates of the same holograms when the galvo pair is at a different location. From top to bottom: n=354,364,362 and 338 holograms respectively. Linearity of these X and Y positions indicates that relative distances between holograms are maintained with galvo scanning. The SLM to real space calibration computed on the central FOV can therefore be linearly translated to other mesoscopic FOV.

**Supplementary Figure 16.**
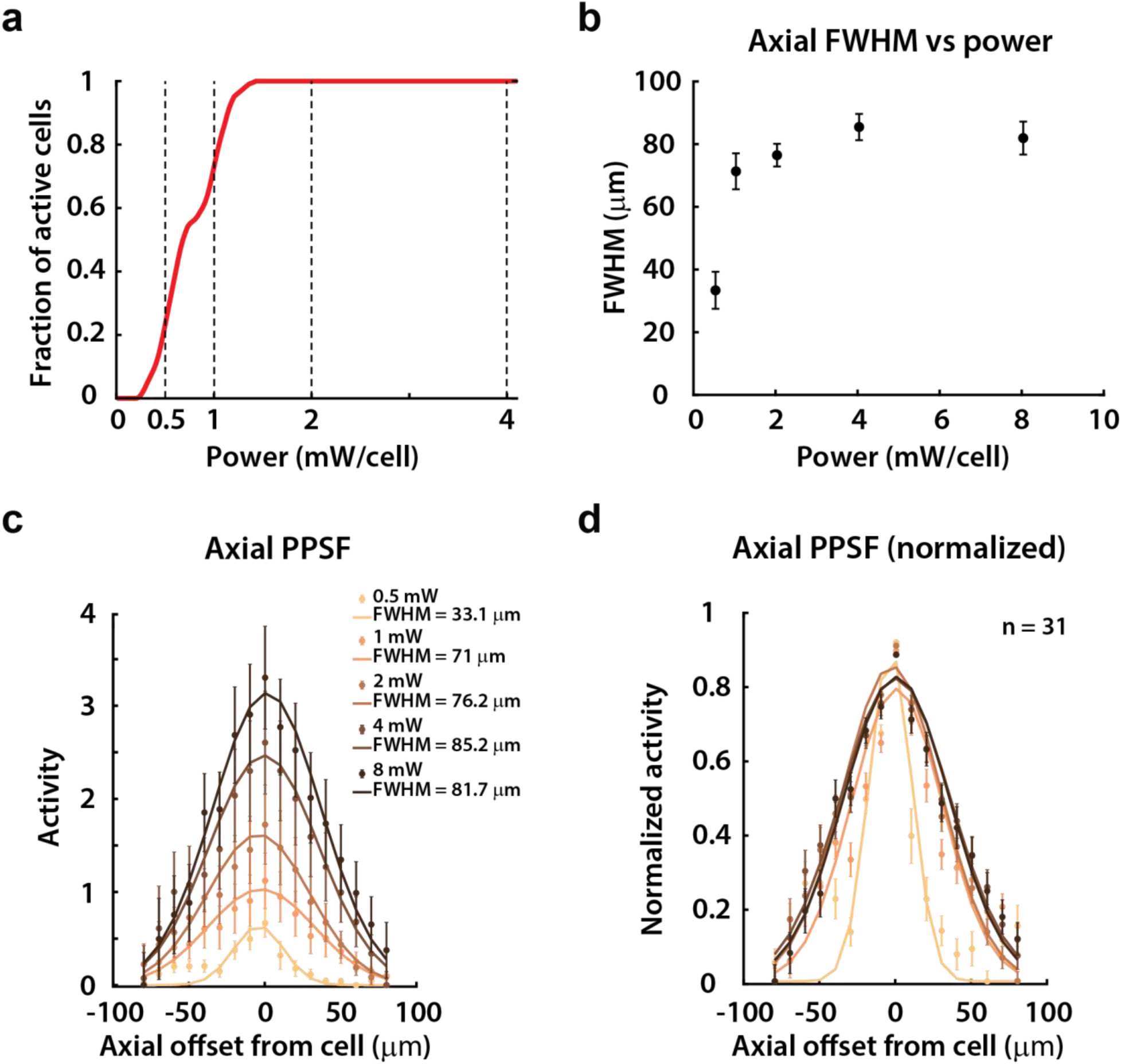
PPSFs as a function of power for off-center holography FOV (Galvo offsets: - 1V, 0V). **a)** Cumulative density histogram of the fraction of active cells used in the PPSF experiments (ordinate) that were active above a certain power (abscissa). **b)** Mean +/- s.e.m. axial FWHM across cells as a function of power. **c)** Axial PPSF profiles for each power condition. The filled circles and error bars are the actual mean +/- s.e.m. across cells, and the solid traces are weighted gaussian fits. **d)** Normalized axial PPSF profiles for each power condition, normalized to the centered hologram location (axial offset = 0). All PPSF experiments were done in CaMK2-tTA; tetO-GCaMP6s mice injected with a pAAV-hSyn-GCaMP6m-p2A-ChRmine-Kv2.1-WPRE virus at discrete locations across visual cortex.

**Supplementary Figure 17.**
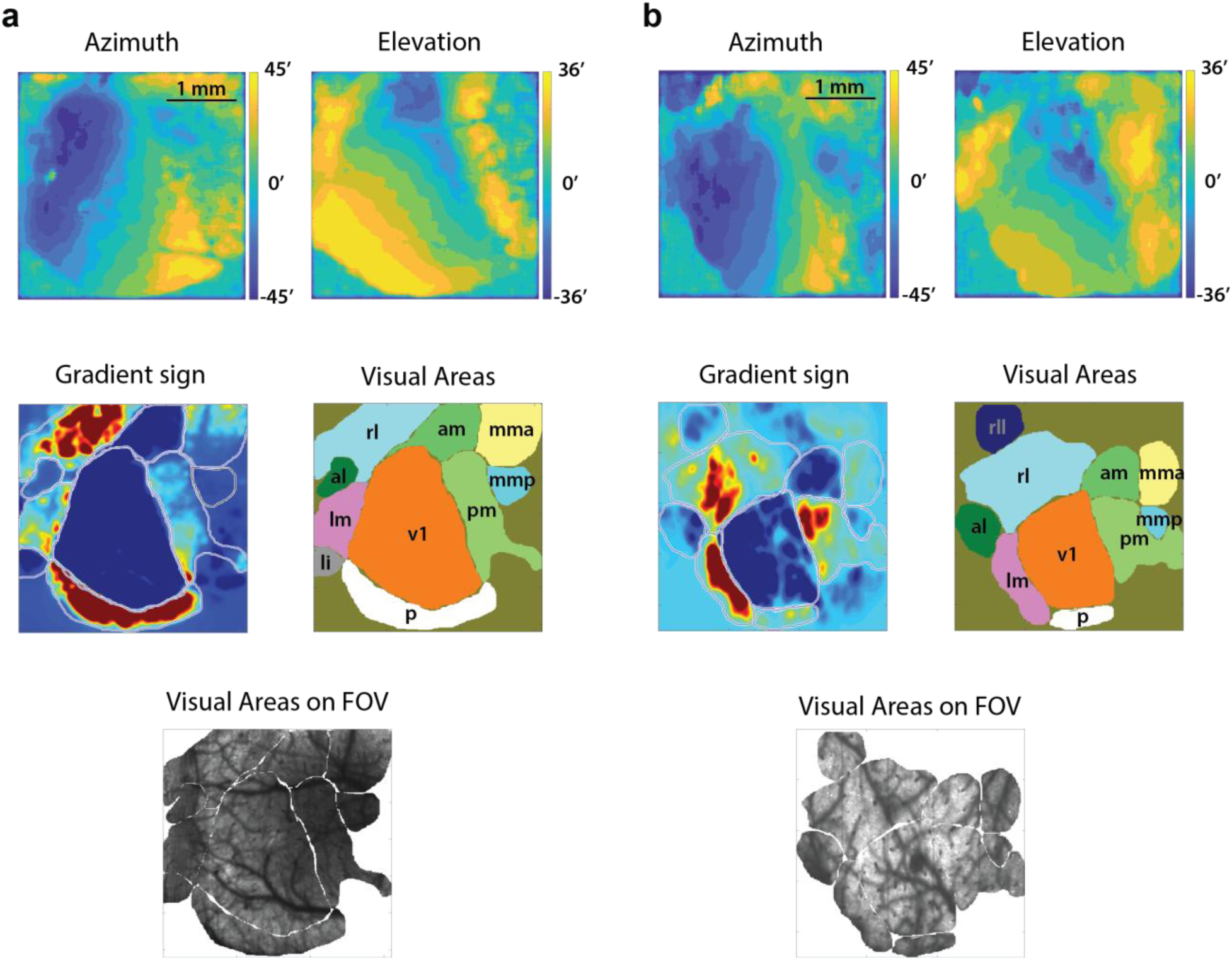
Retinotopic identification of visual areas with the 2P-RAM mesoscope. a-b) Two examples of visual areas identified using retinotopy. Top row: Average pixel-wise azimuth (left column) and elevation (right column) preference maps in response to a moving bar stimulus. Middle row: Gradient sign map computed from the retinotopic preferences (left column), and visual areas segmented manually based on transitions in the gradient sign (right column). Bottom row: Segmented visual areas overlaid on the 3 x 3 mm FOV for vasculature-based alignment across sessions.

**Supplementary Figure 18.**
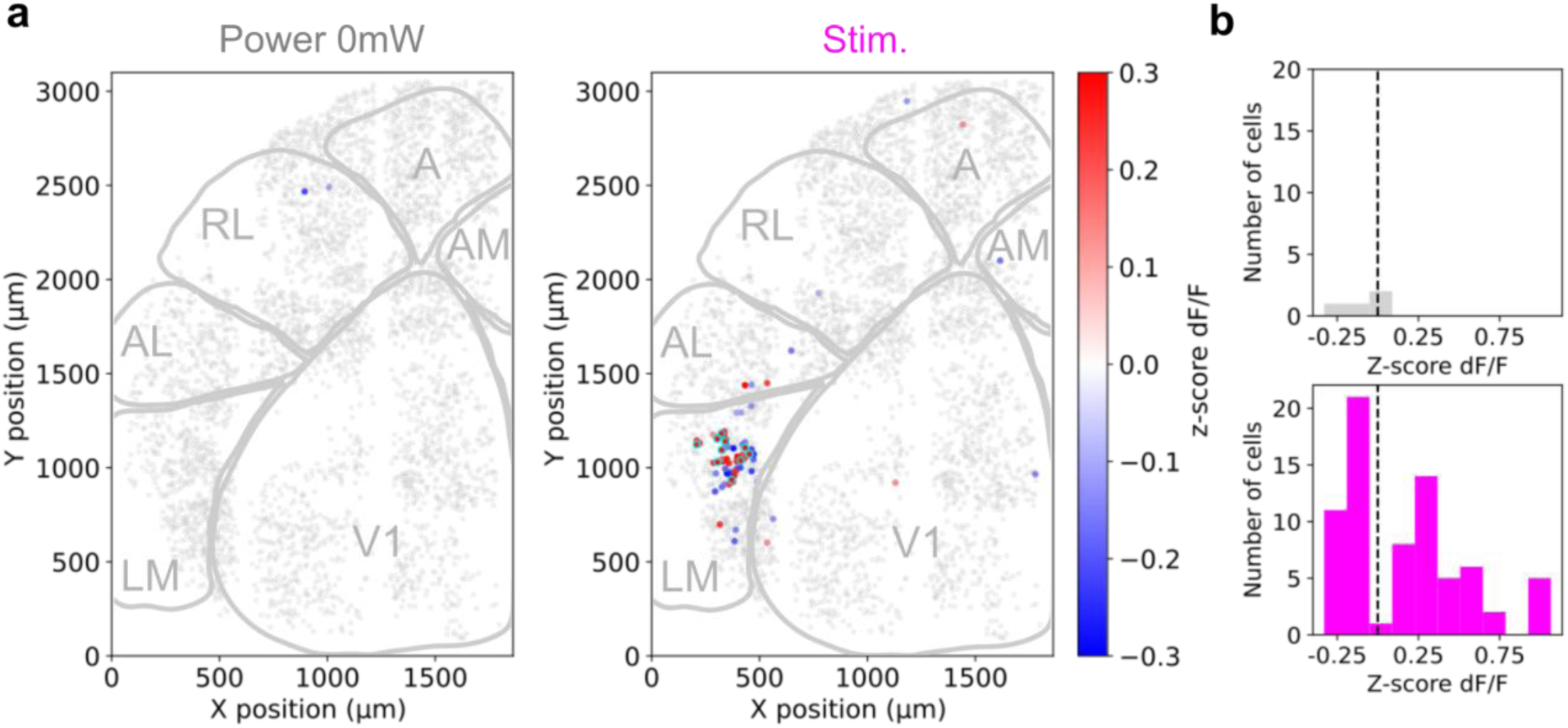
Cross-validated influence maps for holographic stimulation and no stimulation control. **a)** Example mesoscale holographic ‘influence’ maps measured in response to no holographic stimulation (left) and to activation of an ensemble of LM neurons (right). Significantly modulated cells are colored according to the amplitude of their calcium responses. Significantly modulated cells are selected as the intersection of significantly modulated cells (t-test p-value <0.05) across two random splits of the total number of trials. Non-modulated cells are displayed in gray. Holographic targets are circled in cyan. Mouse: Ai203xVGlutCre. **b)** Distribution of the amplitude of the post-stimulation calcium responses of significantly modulated cells in response to holographic activation (magenta) compared to control trials with no stimulation (grey).

**Supplementary Figure 19.**
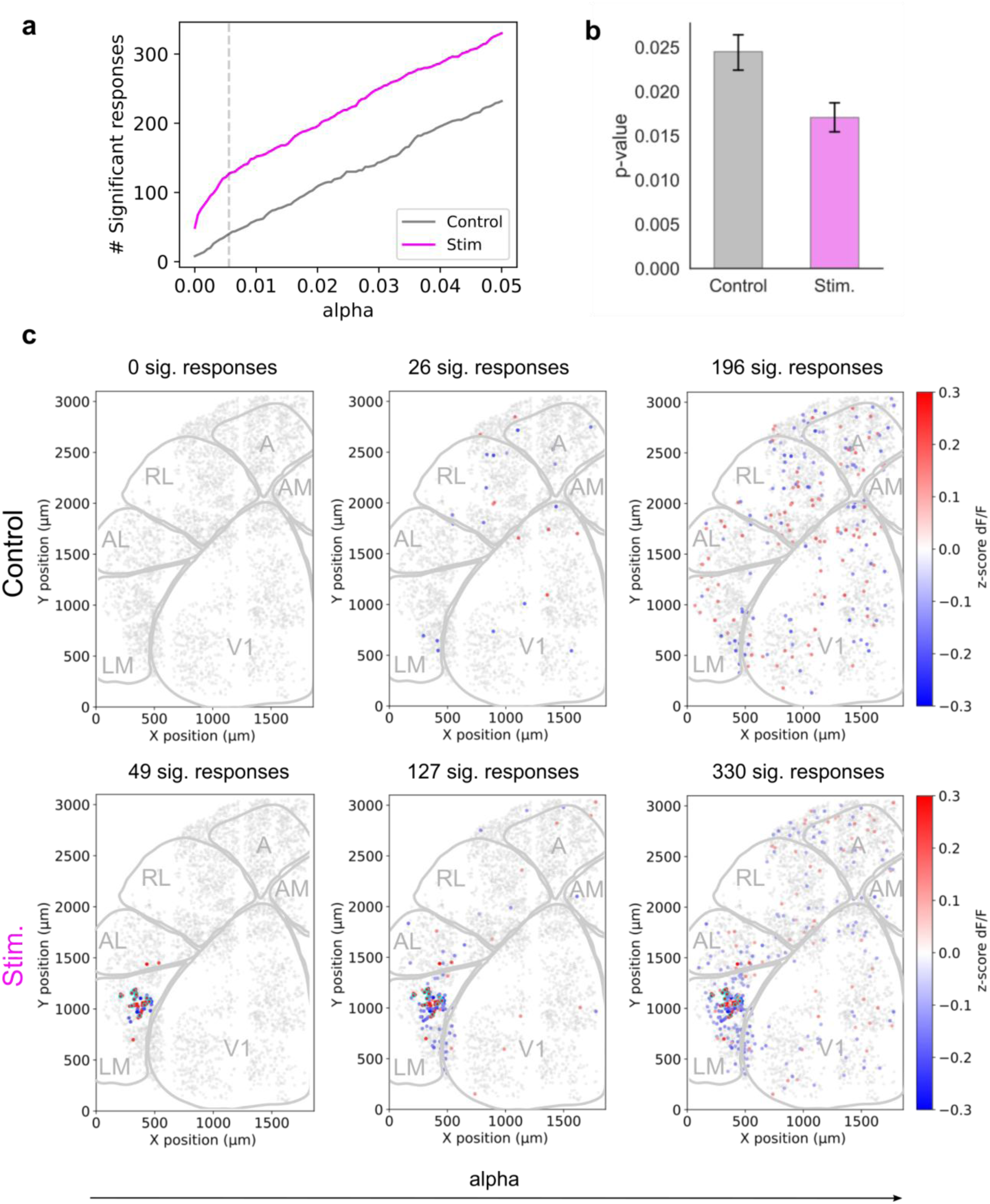
Significant modulations and false positive rates. **a)** Number of significantly modulated cells in response to holographic activation and in response to the no stimulation control as a function of the significance level parameter alpha. Holographic stimulation consistently elicits more significant responses than the false positive rate. Calcium indicator:GCaMP6s, Opsin: ChroME2s. **b)** Average of p-values for significantly modulated cells (t-test, p-value <0.05) in the stimulation and no stimulation control condition. **c)** Maps of significantly modulated cells as a function of the significance level alpha parameter. Alpha parameter from left to right: Bonferroni (1.35×10^-5^), Inflection point value in a) (5.6×10^-3^), and 0.05.

**Supplementary Figure 20.**
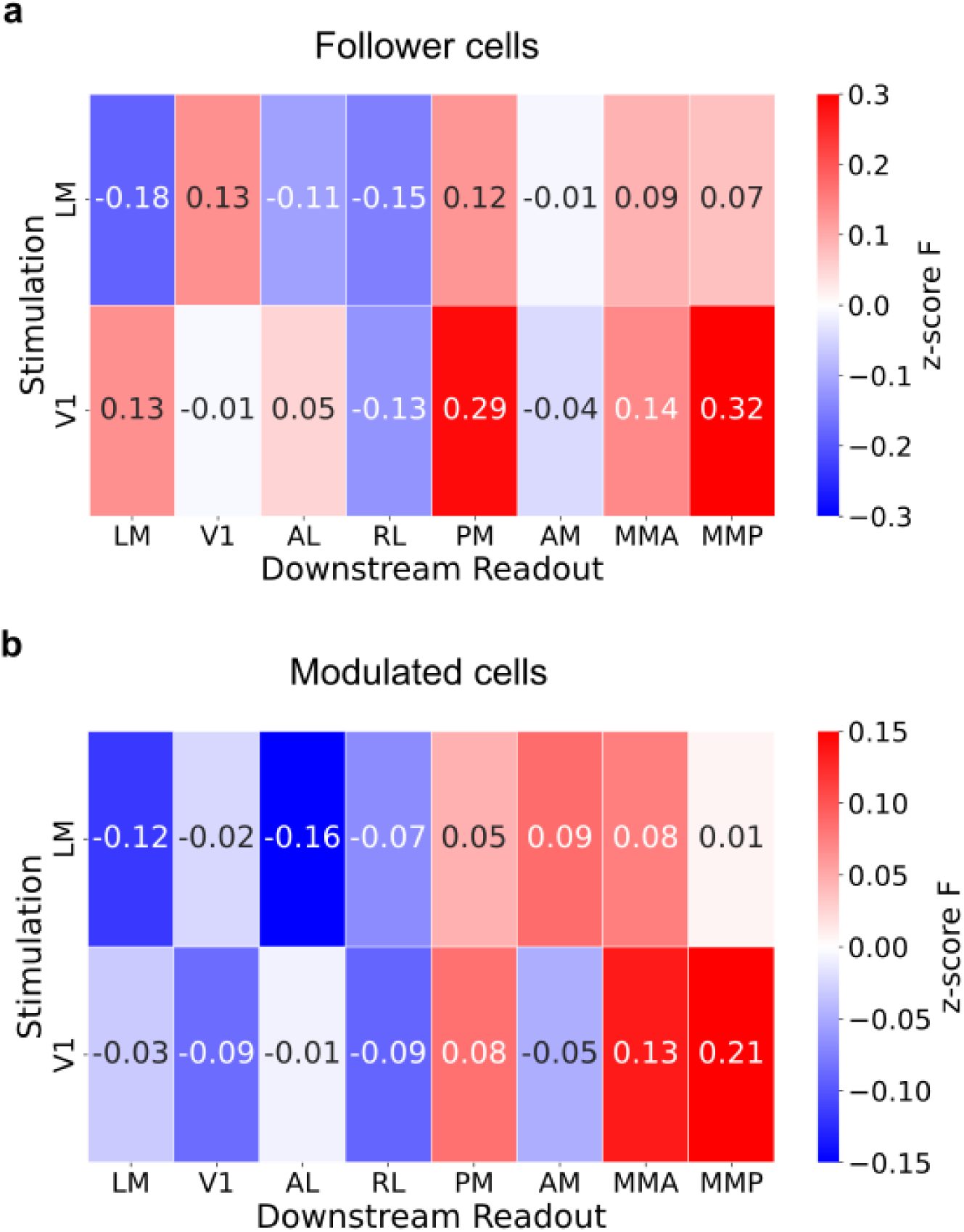
Inter-areal functional connectivity maps from follower cells and modulated cells. **a)** Inter-areal causal mapping of net influence across cortical areas computed from post-stimulation responses of follower cells (a subset of highly reliable modulated cells, see Methods). N=6 sessions, 2mice. Area boundaries are defined using online two-photon retinotopic mapping (see supp. Fig. 17 and Methods). This connectivity matrix hence corresponds to a functional connectivity matrix of strong and sparse connections. **b)** Same as a) but computed from post-stimulated responses of modulated cells (all cells significantly responsive to photostimulation, see Methods). This connectivity matrix hence by definition also includes weaker connections.

**Supplementary Figure 21.**
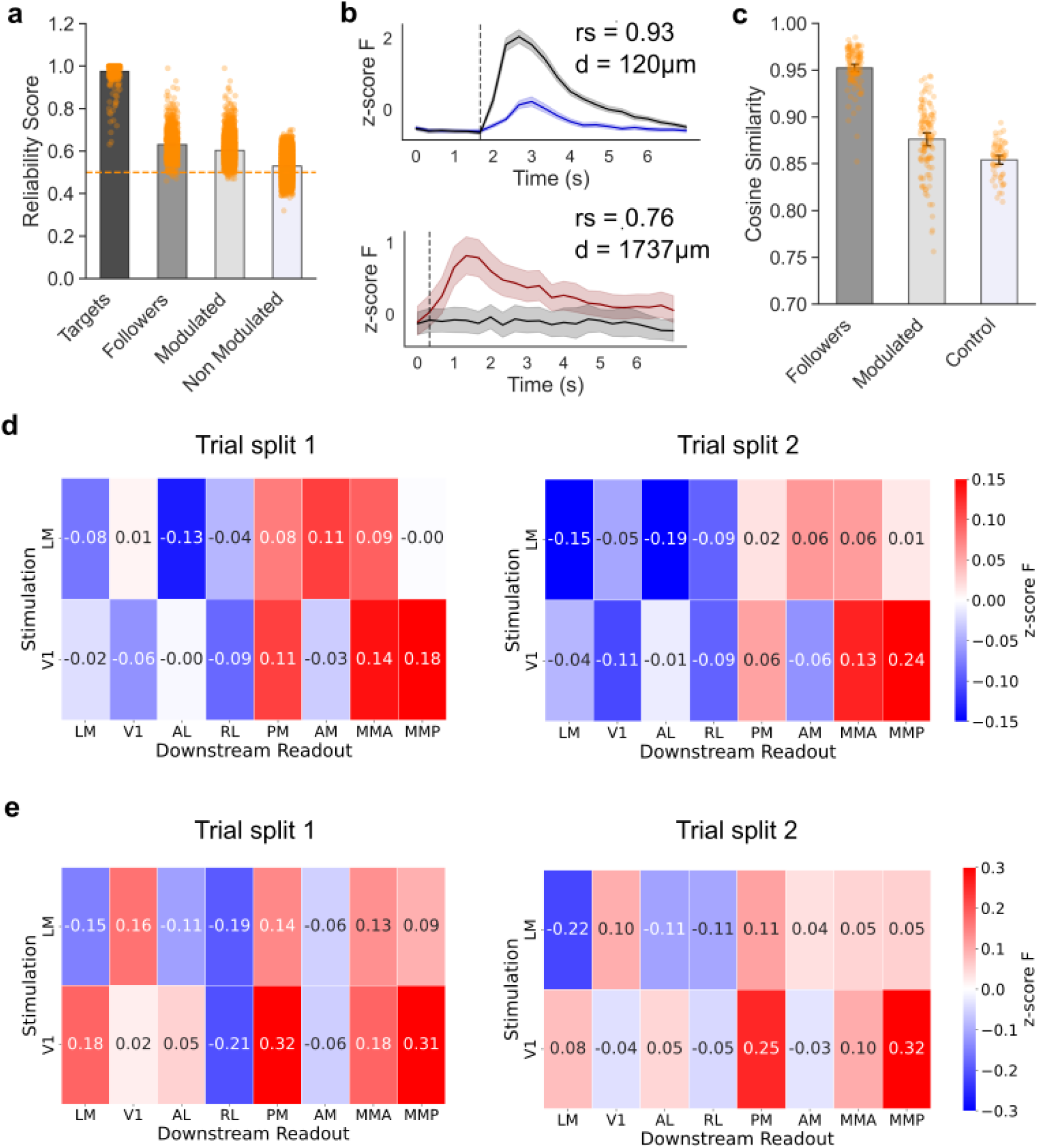
Reliability of post-synaptic responses across trials. **a)** Single-trial reliability of target, follower, modulated, and non-modulated cells. Modulated cells were located >100 µm from the nearest target and showed significant responses to photostimulation (p < 0.05; see Methods). Follower cells were defined as a subset of highly reliable modulated cells (see Methods). Reliability was quantified as the fraction of trials in which the sign of post-stimulus modulation matched that of the average modulation. Post-stimulus modulation was computed as the difference between responses in photostimulation and control (no-power) trials. *n* = 417 (targets), 931 (followers), 4,298 (modulated), and 50,629 (non-modulated), pooled across 6 sessions from 2 animals. To avoid sample size bias, 1,000 cells were randomly subsampled from each non-target group for statistical testing. Mann-Whitney U tests: Followers vs. Non-modulated, p = 3.14E-140; Followers vs. Non-modulated, p= 3.1×10^-225^; Modulated vs. Non-modulated, p = 6.2×10^-218^. **b)** Example calcium responses from follower cells with high reliability. Average calcium traces from two follower cells with reliability scores of 0.93 and 0.76. Blue or red traces show responses during photostimulation trials; black traces indicate responses during control (no-power) trials. **c)** Similarity of population response vectors across bootstrapped trial splits. For each experiment, photostimulation trials were randomly divided into two non-overlapping subsets, repeated over 10 bootstrap iterations. Population vectors of post-stimulus modulation were computed separately for each subset and condition: follower cells, modulated cells, and a control group consisting of significantly modulated cells in no-power trials. Both modulated and follower cells showed significantly higher cosine similarity than control cells (Mann–Whitney U test: modulated vs. control, *p* = 8.4×10^-7^; follower vs. control, *p* = 3.5E-27). *n* = 120 (modulated), 120 (follower), and 60 (control). The distribution of follower cell cosine similarity was entirely non-overlapping with control values. **d)** Inter-areal functional connectivity maps from trial-split population responses. Connectivity maps were computed from modulated cells using population vectors derived from two non-overlapping subsets of both photostimulation trials and control (no-power) trials. N=2 animals, 6 sessions. **e)** Inter-areal functional connectivity maps from trial-split population responses. Connectivity maps were computed from follower cells using population vectors derived from two non-overlapping subsets of both photostimulation trials and control (no-power) trials. N=2 animals, 6 sessions.

**Supplementary Figure 22.**
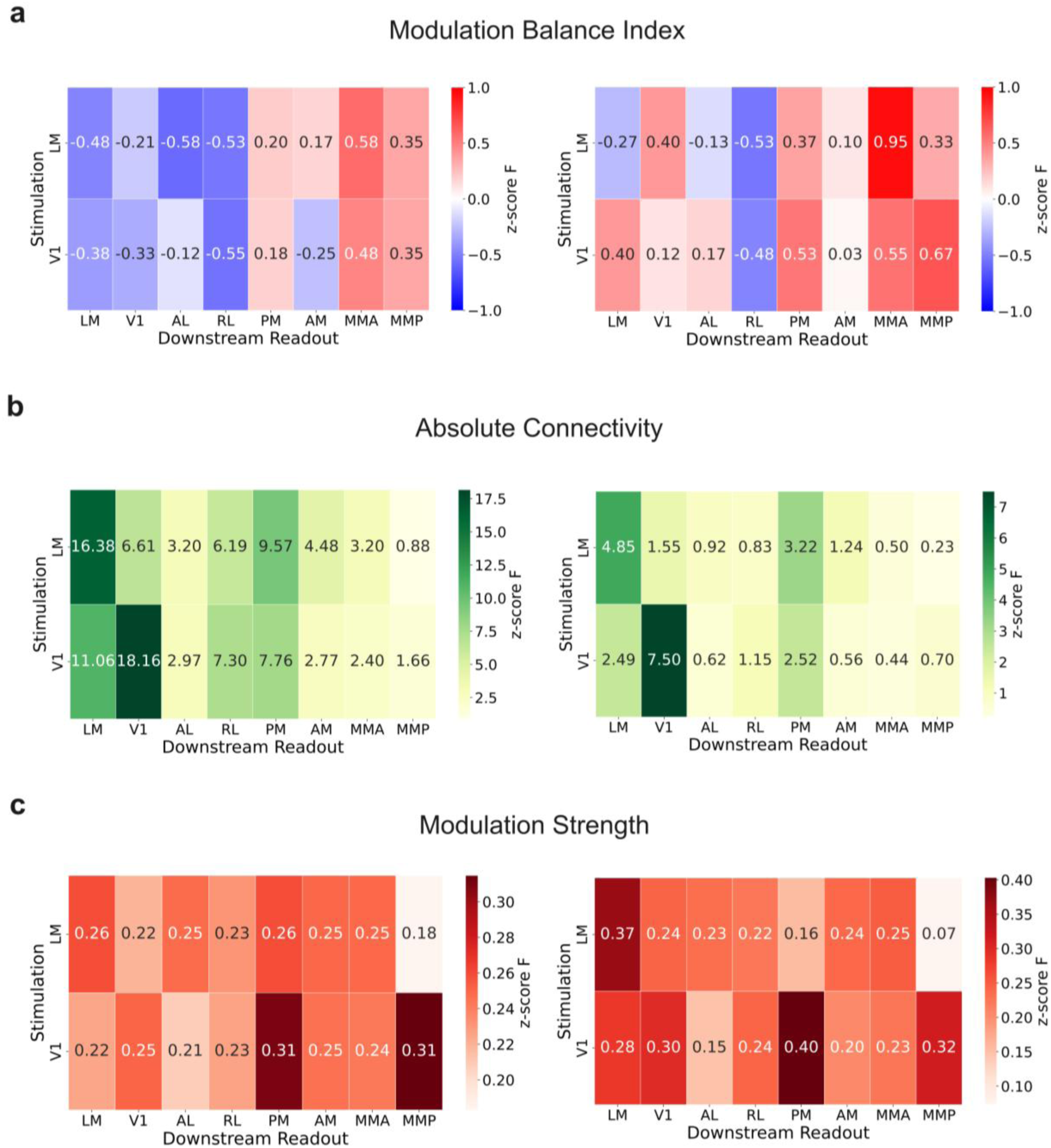
Inter-areal functional connectivity metrics for photostimulation in V1 and LM. **a)** The modulation balance index quantifies the relative dominance of excitation vs. inhibition among significantly modulated downstream neurons (left) or follower neurons (right). It is defined as Modulation Balance Index = 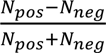 where N and N are the numbers of significantly positively and negatively modulated cells, respectively. The index ranges from -1 (pure inhibition) to +1(pure excitation), with 0 indicating a balanced mix. **b)** The absolute connectivity metric quantifies the total strength of downstream modulation elicited by photostimulation, regardless of polarity. It is calculated as the sum of the absolute modulation responses across all significantly modulated neurons (left) or follower neurons (left). Absolute Connectivity = 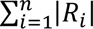 where the sum is taken over all n significantly modulated cells. This metric reflects the overall magnitude of the network response. **c)** The modulation strength metric quantifies the average amplitude of photostimulation-induced responses among significantly modulated neurons (left) or follower neurons (right). It is computed as the sum of the absolute modulation amplitudes divided by the number of significantly modulated cells Nmod. Modulation Strength = 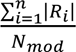. This provides a polarity-independent measure of the average effect size per modulated neuron, allowing comparison of response strength across conditions or cortical areas regardless of the number of recruited cells. All metrics were computed from significantly modulated cells within each visual area (n = 6 sessions, 2 mice).

**Supplementary Figure 23.**
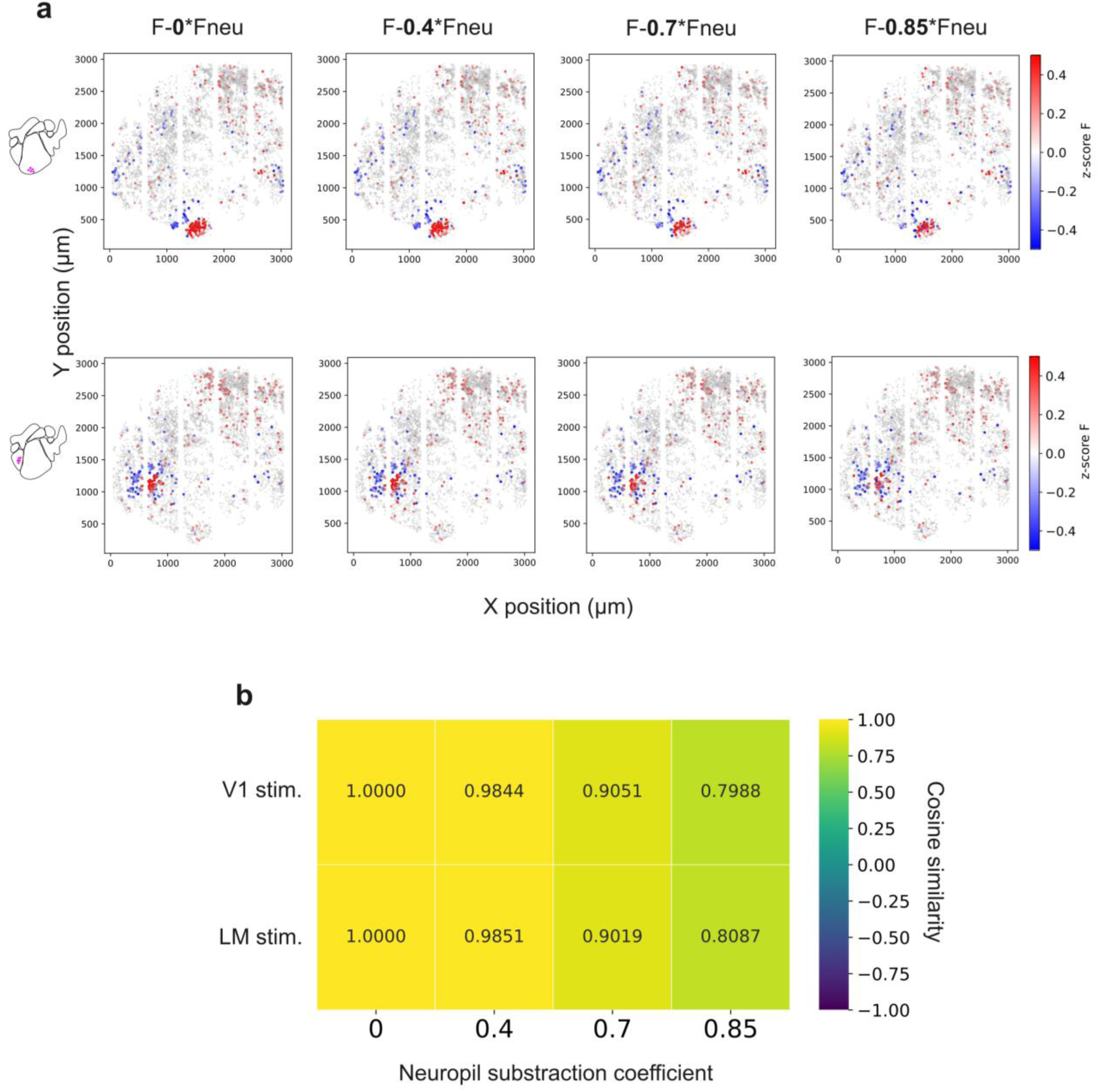
Impact of neuropil subtraction on modulated cell responses. **a)** Example influence maps for photostimulation in V1 (top) and LM (bottom) showing targets and significantly modulated cell responses with different neuropil subtraction coefficients. **b)** Cosine similarity of the population response compared to the uncorrected (no neuropil subtraction) condition.

**Supplementary table 1.** List of optical components (holography path)

|  |  |
| --- | --- |
| L01 | Achromatic doublet $f = 50$ mm, AC254-50-B. **Thorlabs, USA** |
| L02 | Achromatic doublet $f = 50$ mm, AC254-50-B. **Thorlabs, USA** |
| D1 | Shortpass Dichroic Mirror, DMSP1000R. **Thorlabs, USA** |
| D1 (RAM 3D-SHOT) | 1040 nm StopLine notch laser dichroic, $50 \times 50$ mm. **AVR Optics USA** |
| D2 | 980 nm laser BrightLine single-edge dichroic beamsplitter, 25 mm diameter. **AVR Optics, USA** |
| G | Diffraction Grating, Plane Ruled $50 \times 50$ mm, 1000nm, 600Gr/mm, 33010FL01-520R. **Newport Corporation, USA** |
| L1 | Achromatic doublet $f = 125$ mm, AC254-125-B-LM. **Thorlabs, USA** |
| L2 | Aspheric Lens $f = 20\text{mm}$, AL2520H-B. **Thorlabs, USA.** |
| D | 0.5° Diffusing Angle 50 mm, #47-989. **Edmund Optics, USA** |
| L3 | Achromatic doublet $f = 500\text{ mm}$, ACT508-500-B. **Thorlabs, USA** |
| SLM | 1920 × 1152 Phase Series Spatial Light Modulator, **Meadowlark Optics, USA\*** |
| L4 | Achromatic doublet $f = 150\text{ mm}$, AC508-150-B. **Thorlabs, USA** |
| L5 | Achromatic doublet $f = 300\text{ mm}$, ACT508-300-B. **Thorlabs, USA** |
| BRU/L1 (RAM 3D-SHOT) | Achromatic doublet $f = 150\text{ mm}$, AC508-150-B. **Thorlabs, USA** |
| BRU/L2 (RAM 3D-SHOT) | N-BK7 Plano-Concave lens $f = -75\text{mm}$, LC1315. **Thorlabs, USA.** |
| XY Galvos (RAM 3D-SHOT) | Beam Galvo system QS30XY-AG. **Thorlabs, USA** |
| L6 | Achromatic doublet $f = 200\text{ mm}$, ACT508-200-B. **Thorlabs, USA** |
| L6 (RAM 3D-SHOT) | F-theta scan lens FTH100-1064. **Thorlabs, USA.** |
| TL | Custom Tube Lens, $f=236\text{mm}$. **Jenoptik USA**<br>Optical design file can be found at:<br>https://wikis.janelia.org/display/mesoscopy/Optical+design+files. See also Sofroniew et al., 2016. |
| Objective | Custom Objective, $f=21\text{mm}$. **Jenoptik USA**<br>Optical design file can be found at:<br>https://wikis.janelia.org/display/mesoscopy/Optical+design+files. See also Sofroniew et al., 2016. |
| D3 | 1040 nm StopLine notch laser dichroic, 50 × 50 mm. **AVR Optics USA** |
| D3 (RAM 3D-SHOT) | 1040 nm StopLine notch laser dichroic, 75 × 75 mm. **AVR Optics USA** |
| D4 | Shortpass Dichroic Mirror, DMSP1000R. **Thorlabs, USA** |
| Mirrors | Broadband dielectric mirrors, BB2-E03. **Thorlabs, USA** |

Refer to CAD file for optomechanical parts.
\*: This part has unfortunately been discontinued and is no longer sold by the vendor. We briefly discuss below two alternative options from the same vendor:
2) 1024x1024 Spatial Light Modulator, Meadowlark Optics, 17 $\mu\text{m}$ pixel pitch. With the current optical design, this device can enable similar axial resolution performances, at the expense of a reduced holographic FOV. Adjustments on the relay lenses can be made to adjust the resolution/FOV trade-off, but having simultaneously the same resolution and FOV specs as presented in the manuscript will be challenging. This device has however a very high refresh rate (>500Hz). This device can therefore be used in the random-access 3D-SHOT configuration. Relay optics can be optimized for resolution and the limited FOV can be compensated for by the galvanometer system.

**Supplementary table 2.** Groupe delay dispersion estimation (holography path)

| | Part # | Material | Thickness (mm) | GVD at 1.03 $\mu\text{m}$<br>(fs<sup>2</sup>/mm) | GDD at 1.03 $\mu\text{m}$<br>(fs<sup>2</sup>) |
| --- | --- | --- | --- | --- | --- |
| L01 | AC254-50-B | N-LAK22 | 7.5 | 43.78 | 328.35 |
|  |  | N-SF6HT | 1.8 | 241.67 | 435.006 |
| L02 | AC254-50-B | N-LAK22 | 7.5 | 43.78 | 328.35 |
|  |  | N-SF6HT | 1.8 | 241.67 | 435.006 |
| L1 | AC254-125-B | N-BK7 | 6 | 25.123 | 150.738 |
|  |  | N-SF8 | 6 | 88.483 | 530.898 |
| L2 | AL2520H-B | S-LAH64 | 7.6 | 63.455 | 482.258 |
| L3 | ACT508-500-B | N-SK2 | 6 | 41.15 | 246.9 |
|  |  | N-SF5 | 6 | 155.86 | 935.16 |
| L4 | AC508-150-B | N-LAK22 | 8.2 | 43.78 | 358.996 |
|  |  | N-SF6HT | 5 | 241.67 | 1208.35 |
| L5 | ACT508-300-B | N-SK2 | 8 | 41.15 | 329.2 |
|  |  | N-SF11 | 6 | 125.35 | 752.1 |
| L6 | ACT508-200-B | N-SK2 | 8.2 | 41.15 | 337.43 |
|  |  | N-SF57 | 5 | 271.33 | 1356.65 |
| Tube lens | Jenoptiks/Thorlabs |  |  |  | 1.20E+03 |
| Objective | Jenoptiks/Thorlabs |  |  |  | 5775 |

The total group delay dispersion (GDD) introduced by the holographic path is estimated at 15190 fs<sup>2</sup>.

